# High-quality genome assembly for the genetically improved Abbassa Nile tilapia enables the reconstruction of X and Y haplotypes

**DOI:** 10.1101/2025.02.28.640774

**Authors:** GJ. Etherington, A. Ciezarek, T. Mehta, T. Barker, A. Durrant, F. Fraser, S. Henderson, N. Irish, GG. Kaithakottil, V. Knitlhoffer, S. Ali, T. Trong, C. Watkins, D. Swarbreck, K. Gharbi, JAH. Benzie, W. Haerty

## Abstract

The success of the Nile tilapia (*Oreochromis niloticus*) as an aquaculture species is partly the result of continuous selective breeding leading to high performing strains. These elite strains have been derived from breeding populations of diverse origins and crosses with other *Oreochromis* species. Owing to the complex and unique evolutionary histories of each strain, existing reference genomes of wild populations are unsuitable to implement genomic selection for beneficial traits such as growth or environmental resilience in aquaculture programmes. Here we generated a high-quality genome assembly and annotation of the WorldFish Genetically Improved Abbassa Nile tilapia (GIANT) elite strain using a combination of PacBio HiFi, and Omni-C Illumina sequencing. As a male Abbassa Nile tilapia was used for the generation of the genome assembly, we reconstructed both X and Y haplotypes, identifying both *amhY* and *amhΔy on* LG23 indicating that Abbassa likely shares the same sex determination system as GIFT, and thereby differs from the existing reference genome, whose sex determination loci are located on LG1.

## Background

For the first time, aquaculture production exceeded wild capture output in 2022 reaching 51% of the total fish industry yield (185 million tonnes)^1^. Inland finfish production represented nearly 53 million tonnes in 2022, with the Nile tilapia (*Oreochromis niloticus*) being the third most important species, representing 10% of the global production (5.3 million tonnes)^1^. The success of the Nile tilapia is mainly due to active breeding programmes with a strong focus on improving growth rate and weight at maturity. Those programmes led to the generation of several elite strains including the FaST (Bolivar, 1998), GET-EXCEL^2^ as well as the WorldFish GIFT (Genetically Improved Farmed Tilapia) and GIANT (Genetically Improved Abbassa Nile Tilapia, hereafter referred to as ‘Abbassa’)^3^ strains. The Abbassa programme was developed independently from the GIFT strain in Egypt in 2001^4,5^ using a different and diverse initial set of wild (Aswan, Zawia, and Abbassa) and farmed Egyptian populations to initiate the breeding programme. The programme led to a strain with increased yield and lower feed conversion ratio relative to other strains^6^.

To accelerate the improvement of yield traits, there is a strong incentive to better understand the genetic bases underlying those traits enabling genomic selection. Additionally, there is increasing pressure to secure yield through the development of strains with increased resistance to pathogens and diseases (e.g., Tilapia Lake Virus^7^), and enhanced resilience to environmental variation due to climate change such as increased temperature and salinity. The availability of high-quality genomic resources, including an accurate genome assembly and annotation, is key to enhancing breeding of desirable traits in genomic selection programmes through the identification and utilisation of genome-wide marker information.

The current reference genome for the Nile tilapia was generated from a homozygous clonal line produced at the University of Stirling (Scotland) and initiated from individuals originating from Lake Manzala (Egypt)^8,9^. In contrast, the vast majority of the elite *O. niloticus* strains include significant introgression from other *Oreochromis* species during their breeding process^3^ and thus, do not share the same evolutionary history as the reference genome individual, significantly limiting the use of this reference genome for global breeding purposes. As such, we recently demonstrated substantial introgression with *O. mossambicus* during the breeding process of the GIFT strain, leading to the identification of over 11Mb of *O. mossambicus* genomic sequence, harbouring 283 genes, within this elite strain^10^. Similarly, introgression of *O. aureus* during the breeding process of the Abbassa strain has been previously reported^11^, but the extent to which has not been previously described. Moreover, loci involved in sex determination in Nile tilapia have been identified on different linkage groups (LG1, LG20, or LG23) depending on the origin of samples^12–17^. Using bulk segregant analysis, sex determination loci were identified on LG1 for the reference strain^18^ and LG23 in GIFT^19^ implying again that the current reference genome might not be suitable for the development of genomic selection for this strain.

To enable the future implementation of genomic selection approaches to the Nile tilapia Abbassa strain, we generated a high-quality genome assembly and annotation allowing for a quantification of the level of introgression with other *Oreochromis* species. Most importantly, the high-contiguity of the genome assembly allowed for a full reconstruction of the X and Y haplotypes for this strain, identifying the same genetic loci and variation associated with sex determination previously identified in the GIFT strain through bulk segregant analysis and differing from the one reported for the main reference assembly^19^. These observations once again stress the importance of the generation of strain specific genomic resources to facilitate the future application of genomic selection in tilapia breeding.

## Results

### Genome assembly

Using PacBio Hifi reads and Omni-C Illumina short reads generated from the same animal donor, we produced a high-quality genome assembly for the Abbassa strain (Figure 1A). The total assembled genome size of Abbassa is 1.095 Gb across 223 scaffolds. The assembly size is slightly larger than those of the current reference Nile tilapia genome assembly (Ensembl v110, hereafter UMD) and GIFT^10^ (1.01 Gb and 1.07 Gb respectively) and contained in fewer contigs (2460 in UMD, 344 in GIFT). The N50 statistics show that Abbassa has a slightly superior contig N50 compared to GIFT and notably higher than UMD (Figure 1B). The scaffold N50 is slightly lower than UMD and GIFT, reflecting the lack of the existence of a genetic map enabling anchoring the scaffolds to linkage groups as previously done for UMD^20^ and GIFT^10^. In the Abbassa assembly, 76.7% of the assembly is contained in the largest 23 scaffolds, 90% within the 41 longest scaffolds and 95% within 58 scaffolds.

**Figure 1.**
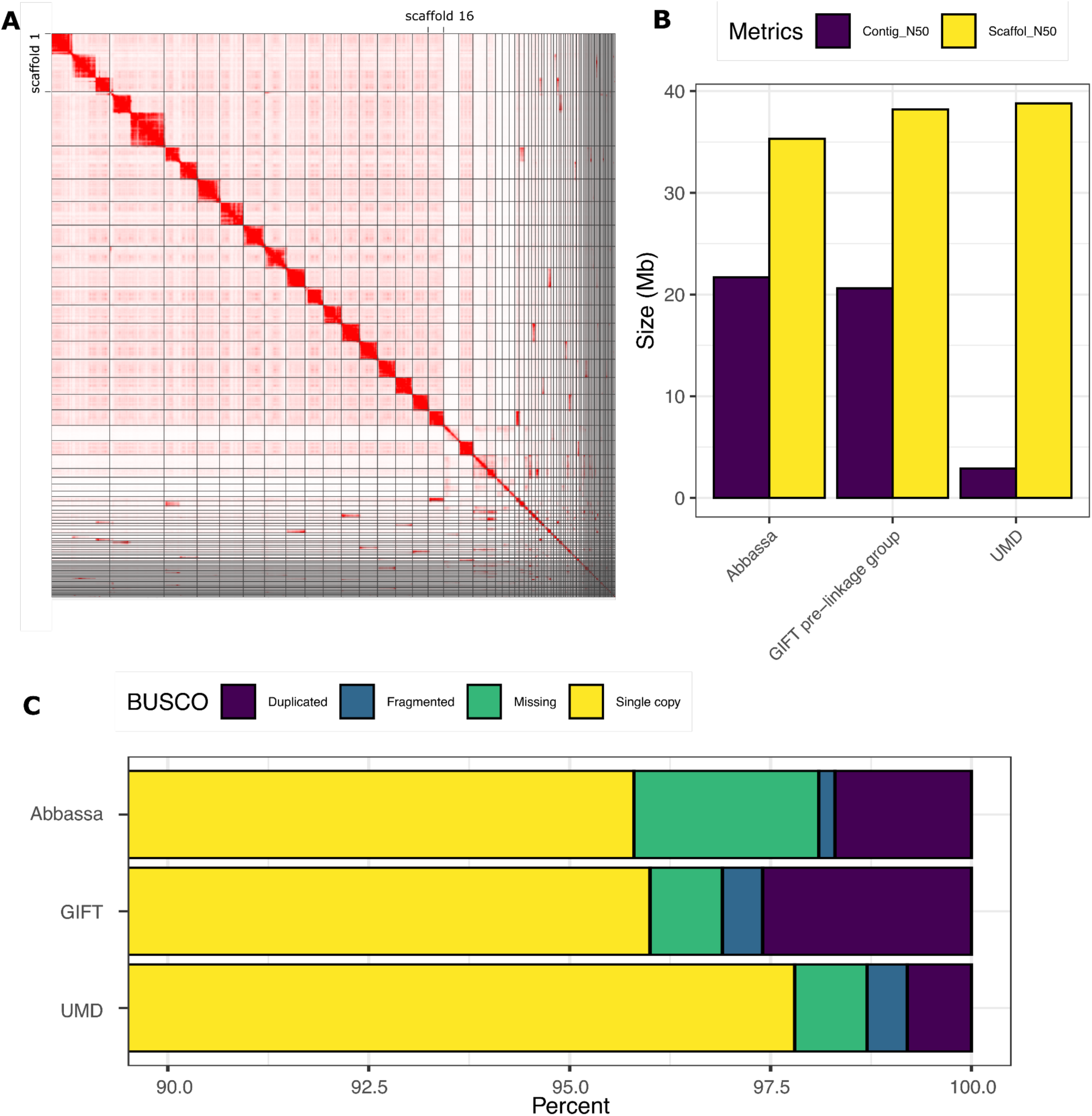
Assembly of the Abbassa genome. **A**. HiC contact map for the Abbassa genome. **B.** Comparison of Contig and scaffold N50 (Mb) for the Abbassa, GIFT, and UMD genome assemblies. For consistency, the assembly statistics for GIFT are prior linkage group mapping. **C.** Number of BUSCO single-copy orthologs identified in the Abbassa, GIFT, and UMD genome assemblies. The x-axis starts at 90% reflecting the high number of orthologs recovered in single copies across all genomes The quality of the assembly is further supported by the high proportion of single-copy orthologs recovered (Figure 1C, Supplementary table 1), that is similar to the GIFT^10^ and UMD^20^ assemblies (95.8% in Abbassa versus 96% in GIFT and 97.8% in UMD respectively). Notably, Abbassa has slightly more missing orthologs (2.3% compared with 0.9% in UMD^20^ and GIFT^10^).

### Genome annotation

The annotation of the Abbassa genome was based on PacBio Iso-Seq libraries from male gonads, gills, heart, and kidney, complemented with existing genes models from 9 closely related cichlid fish species using the REAT pipeline (https://github.com/EI-CoreBioinformatics/reat), that we previously used to annotate the GIFT genome^10^. We report a total of 40,301 high confidence gene models (96,317 transcripts), including 36,288 protein coding genes encoding 76,722 protein coding transcripts (Table 1). To assess the accuracy and completeness of our annotations, we ran BUSCO on the annotated genes identifying 97.97% of complete BUSCOs, and only 0.27% missing (Supplementary table 1) demonstrating that our annotation is of high-quality. These outputs suggest that the increased number of gene models identified in the Abbassa annotation is not associated with more fragmented gene models compared to the GIFT or UMD annotations. Instead, the increased number of identified Abbassa genes and associated transcripts relative to GIFT and UMD likely stems from the number of tissues and the technologies used to generate the transcriptomes (long-read RNA sequencing for Abbassa, short reads for UMD, computational predictions for GIFT) and the annotation pipelines (REAT for GIFT and Abbassa; NCBI RefSeq for UMD) used to generate them.

**Table 1.** Number of genes and transcripts annotated in the Abbassa, GIFT, and UMD assemblies.

| Metric | Abbassa | GIFT | UMD |
| --- | --- | --- | --- |
| Number of genes | 40,301 | 37,789 | 33,162 |
| Protein coding genes | 36,288 | 31,508 | 28,198 |
| Number of Transcripts | 96,317 | 65,649 | 82,099 |
| Protein coding transcripts | 76,722 | 59,220 | 75,555 |

### Genomic variation between assemblies

We previously reported substantial differences in genomic content between the UMD and the GIFT genome assemblies^10^. Here, we apply similar analyses comparing our novel Abbassa assembly to the GIFT^10^ and UMD^20^ assemblies respectively (Figure 2). Genome synteny dot plots of the Abbassa genome against GIFT (Figure 2A) and UMD (Figure 2B) linkage groups show that a chromosome-scale resolution assembly has been achieved for Abbassa with little or no missing genomic regions. As previously described for GIFT^10^ we identified low homology between strains for LG3, which is most likely the consequence of its previously described high repetitive content in UMD^20^, and ancient chromosome fusion event in Nile tilapia^8,21^ (Supplementary table 2).

**Figure 2.**
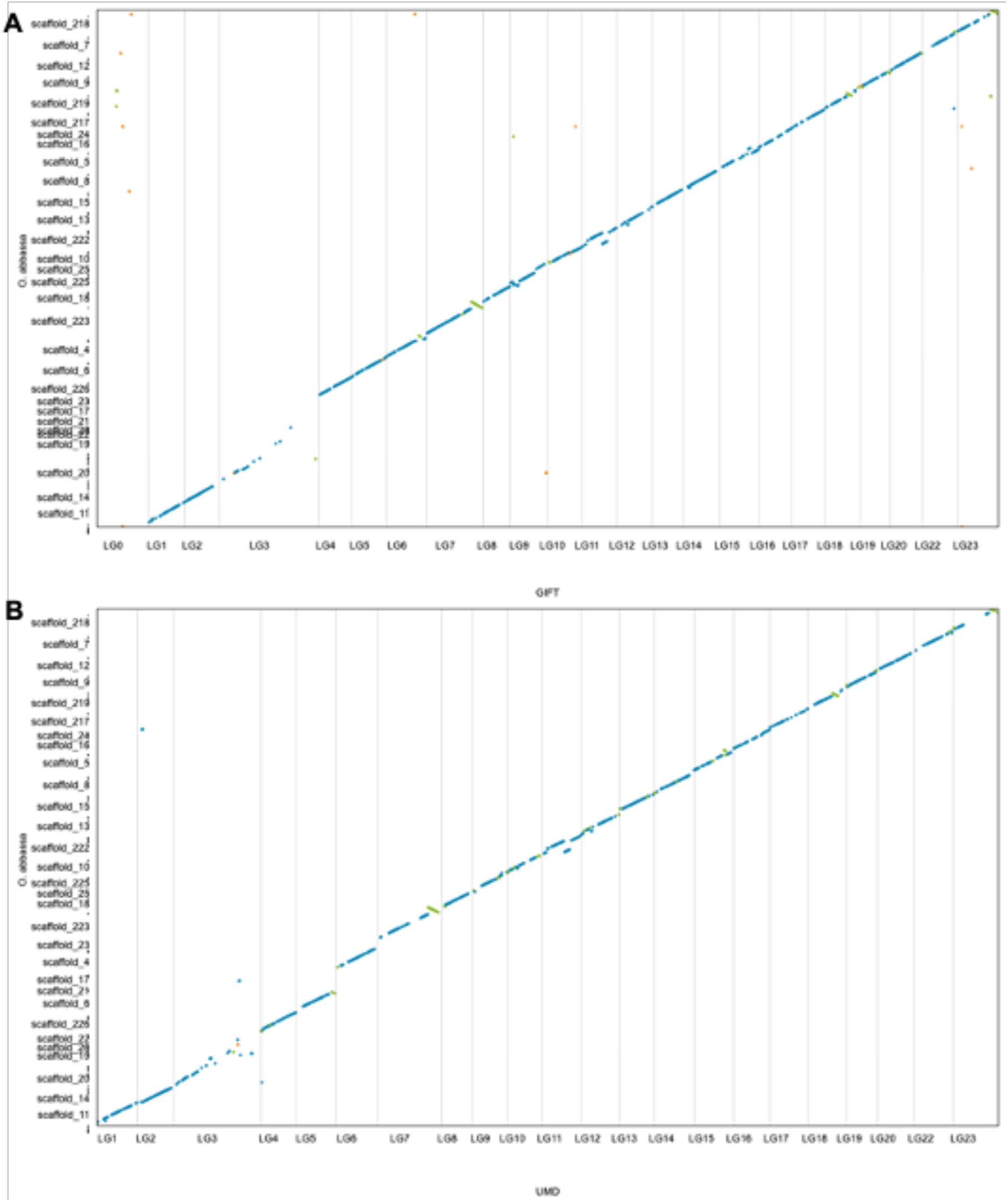
Synteny conservation between Abbassa and GIFT assemblies (A) and between Abbassa and UMD assemblies (B). Blue lines represent unique forward alignments, whilst green lines represent unique reverse alignments. LG0 in the GIFT genome assembly includes all the unplaced sequences.

### Introgression with *O. aureus* during Abbassa breeding

To investigate and characterise the previously reported introgression with *O. aureus*, we called a total of 5.6M SNPs by mapping genomic reads from diverse individuals (GIFT, wild-type *O. niloticus* from lake Albert, *O. aureus*, *O. mossambicus, O. urolepis* and *M. zebra*) against the Abbassa genome (see *Methods*). We applied f-branch analysis to detect admixture and footprints of introgression, and TWISST to quantify taxa relationships.

All except one of the *D* statistics was significantly different from 0 (the trio testing for introgression between either GIFT or Abbassa and *O. aureus*). This is likely as *O. aureus* has introgressed into both GIFT and Abbassa; the *f*-branch analysis showed introgression from *O. aureus* into Abbassa as well as the previously recorded introgression from both *O. aureus* and *O. mossambicus* into GIFT (Figure 3). TWISST analysis shows that the species tree is by the far the most common across the scaffolds, but there are some regions showing an excess of the ancestry where Abbassa and *O. aureus* are most similar to each other, which may represent introgressed regions (Figure 3). The detected regions encompass a total of 40.94Mb, Supplementary Table 3). This extensive signal reflects both the close phylogenetic relationship between *O. niloticus* and *O. aureus*^22^ as well as the existing introgression event during the breeding of Abbassa. However, scaffolds 17,19, 20, 21, 22 and 23 show substantial phylogenetic discordance. These linkage groups also had significantly lower mapping quality and genotype quality than the other scaffolds. The synteny plots (Figure 2), showed that these scaffolds fall in the repeat-rich mega LG3 on GIFT and the UMD reference, which is a fusion of an autosome with a repetitive B-chromosome, rich in retroviral elements, long non-coding RNAs and immunoglobulin genes^8^. This may explain the rampant discordance and poor mapping.

**Figure 3.**
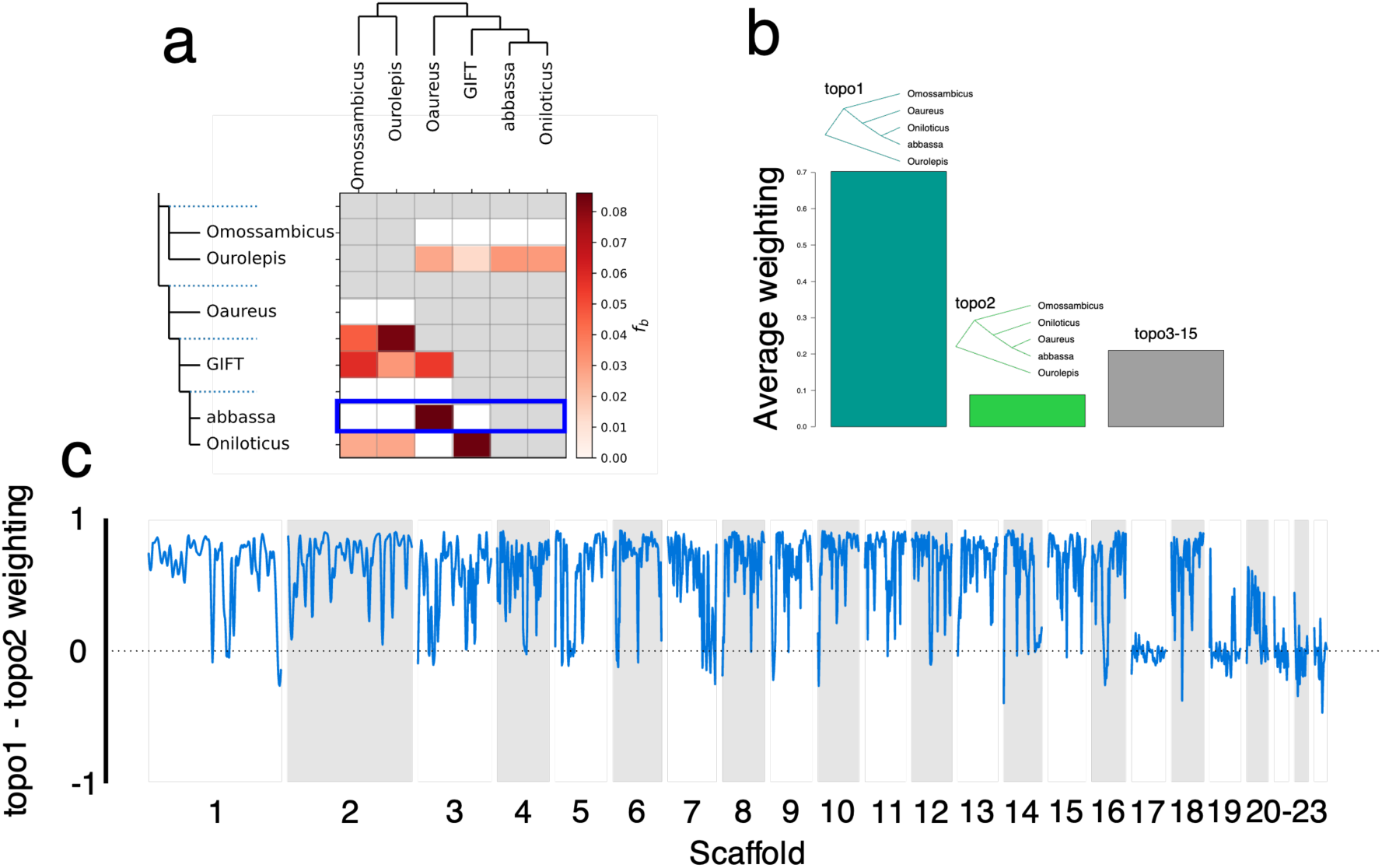
Introgression of *O. aureus* into the Abbassa genome. Phylogenetic representation across genomes of species as estimated by TWISST. **A)** Normalized weights across the *O. niloticus* Abbassa genome assembly for the three different phylogenies. The blue box highlights the row demonstrating the *O. aureus* – Abbassa introgression **B**) Genome-wide average weighting of the species tree, where the reference *O. niloticus* is sister to Abbassa (topo1), the putative introgression tree, where the reference *O. aureus* is sister to Abbassa (topo2) and all other topologies in total (topos3-15) **C)**. Relative weighting topo1 – that of topo2– Negative values reflect regions showing potential introgression as topo2 is more frequent.

### Identification of sex determination locus in Abbassa and comparison with major references

Interestingly, both Abbassa and GIFT show good contiguity on LG 23, while both strains show limited homology for this linkage group with the UMD reference (Figure 2A and 2B). This chromosome is of specific interest as genes on LG23 had been found to be associated with sex determination in GIFT through bulk segregant analysis^19^.

Therefore, we further investigated the Abbassa genome and we confirmed the presence of *amh* and its tandem duplicates, *amhy* and *amhΔy* locus previously identified respectively on the X and Y^14,19^, on LG23 based on breakpoints between the *oaz1* and *dot1l* flanking genes in the Abbassa and GIFT genomes (Fig. 4, a).

**Figure 4.**
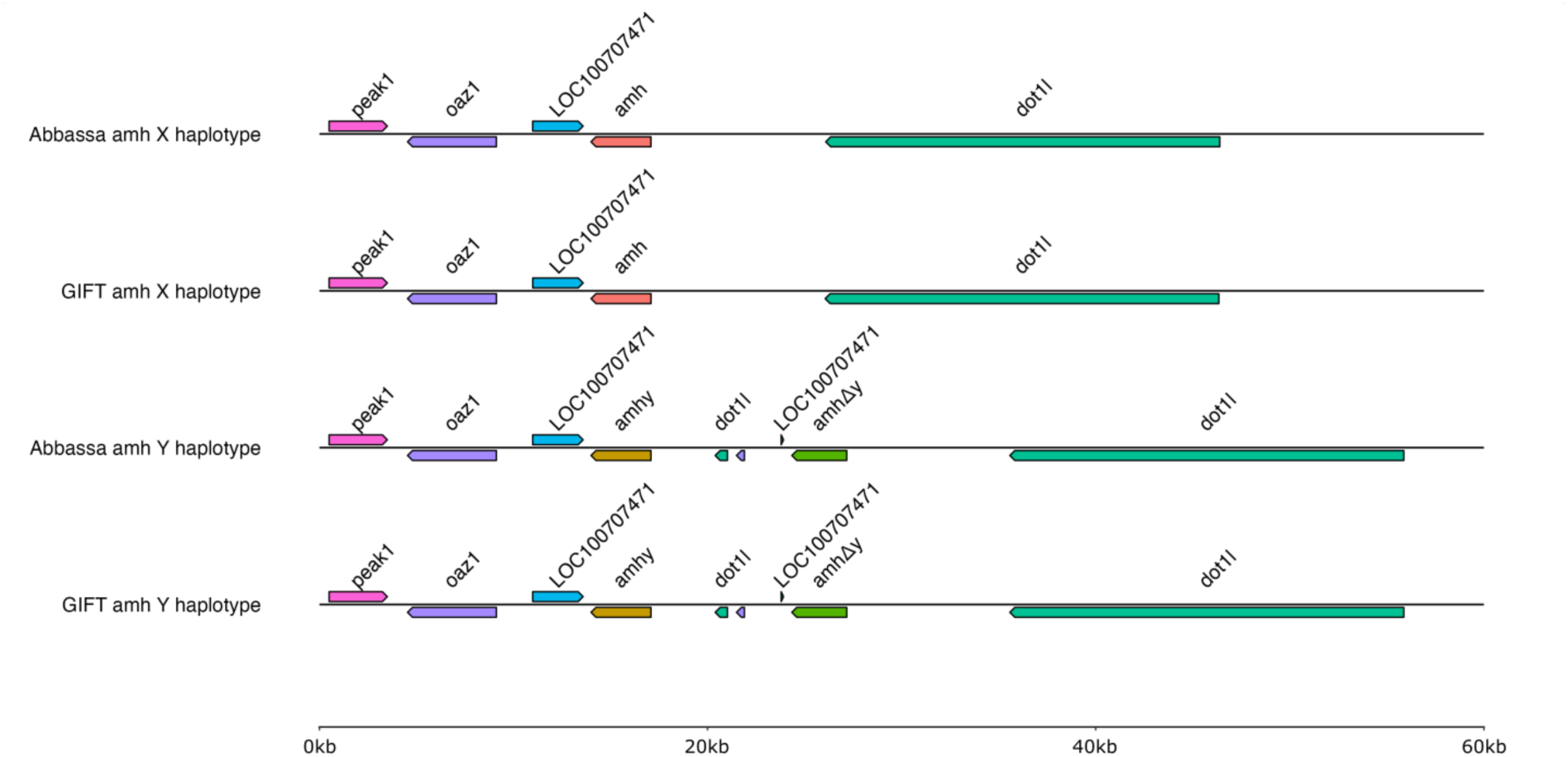
*amh* loci and flanking genes in X and Y haplotypes of Abbassa and GIFT genomes. Alignment of the X-linked *amh* locus in Abbassa and GIFT identifying flanking genes and Y-linked tandem duplicates (*amhy* and *amhΔy*) based on breakpoints between the *oaz1* and *dot1l* flanking genes.

Abbassa and GIFT *amh/amhy* have an ORF of 1,545 bp, encoding putative proteins of 514 aa with the TGF-β domain (Fig. 4, Supplementary Figures 1 and 2). Like previously described for Nile tilapia^14^, both Abbassa and GIFT *amhΔy* have a 777bp ORF encoding a 258aa protein, owing to a 5 bp (ATGTC) insertion in exon VI that creates a premature stop codon, truncating the protein of its TGF-β receptor binding domain (in exon VII) (Fig. 4, Supplementary Figures 1 and 2). We also recovered the previously identified^14^ 346 bp 5’ UTR and 782 bp 3’ UTR of *amh/amhy,* and 46 bp 5’ UTR and 1,242 bp 3’ UTR of *amhΔy* based on alignments of the sequences reported by Li et al^14^ against the Abbassa (Supplementary Figure 3) and GIFT (Supplementary Figures 4) sequences. Most importantly, we identified all previously described genetic variation in the *amhy* and *amhΔy* locus^14,19,23^, including the 5608 bp *amhy* promoter deletion associated with sex determination^14^, the 3 bp ‘AAG’ insertion in the *amhy* gene promoter^19^, and 161 bp deletion in *amhΔy* 5’ UTR^14,19^ (Supplementary Figures 3 and 4). Like previously shown for GIFT populations, neither GIFT nor Abbassa had the C>T substitution previously identified in exon II of Nile tilapia *amhy*^14^, and critical for male sex determination. We also identify several other SNPs and short indels in the gene promoter and 5’ UTR regions of both *amhy* and *amhΔy*, including six SNPs in the 5’ UTR of *amhy* of both Abbassa and GIFT (Supplementary Table 3, Supplementary Figures 3 to 4).

By comparing to the X-linked *amh* gene in Abbassa and GIFT, we identified that *amhy* and its tandemly duplicated gene, *amhΔy*, have genetic variation in associated coding and noncoding regions (Supplementary Table 4, Fig. 5). Whilst there are several conserved coding and noncoding variations in the *amhy* and *amhΔy* genes of Abbassa and GIFT (Supplementary Table 4, Fig. 5, Supplementary Figures 1 and 2), there are also notable unique genetic variations between orthologous genes of the two species. This includes a SNP (at 23 bp G>A) uniquely identified in the 5’ UTR of the Abbassa *amhΔy* gene (Supplementary Table 4, Fig. 5). In the Abbassa genome, the coding sequence of *amhy* is identical to the *amh* gene except for a SNP (A>C) in exon VII, which changes an amino acid (Thr>Ala) in the downstream sequence of the TGF-β binding domain (Supplementary Table 4, Fig. 5, Supplementary Figure 1). Similarly, in the GIFT genome, the coding sequence of *amh* and *amhy* are identical except for a SNP (C>T) in exon VII, that also changes an amino acid (Ala>Val) in the upstream sequence of the TGF-β binding domain (Supplementary Table 4, Fig. 5, Supplementary Figure 2).

**Figure 5.**
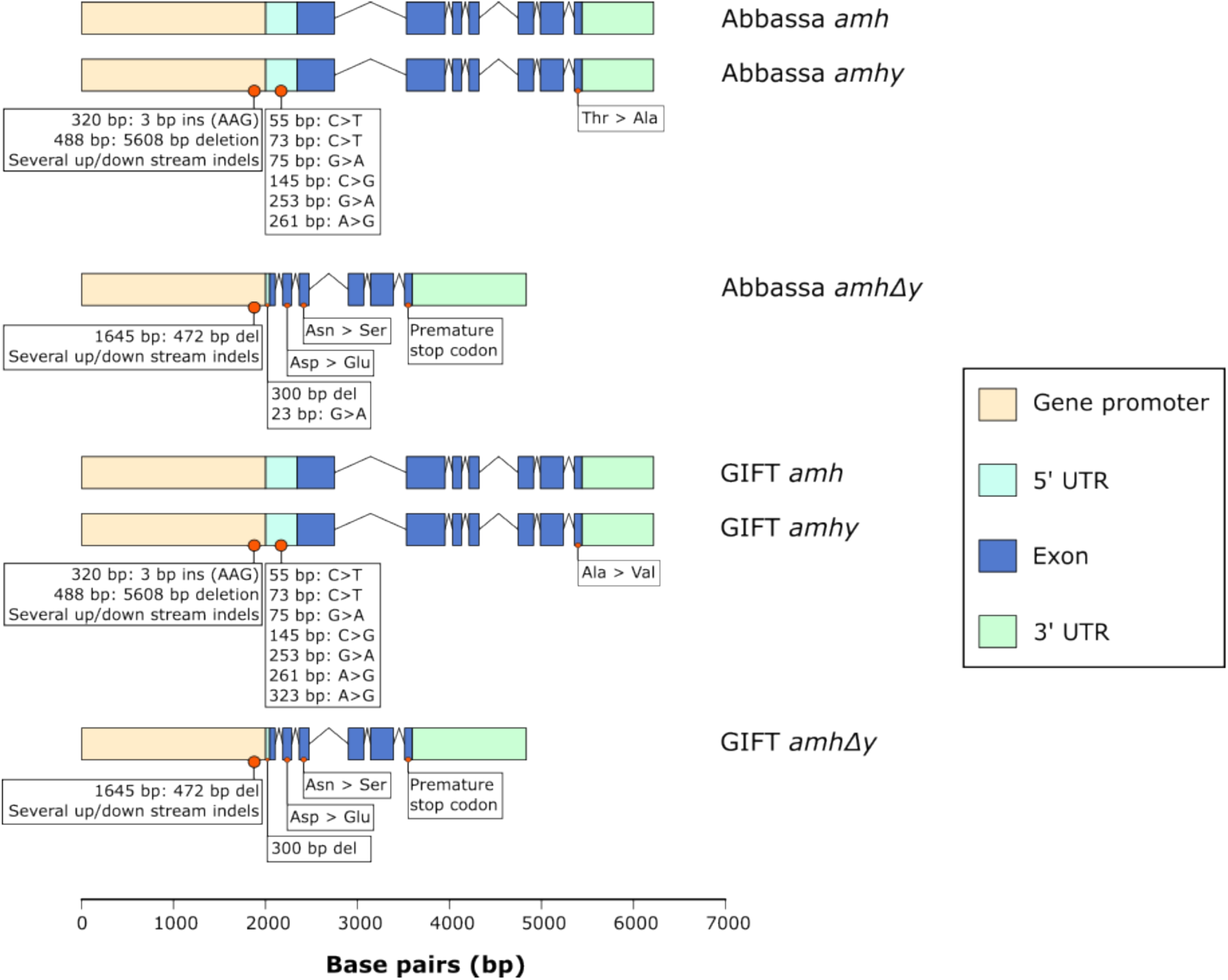
Gene structure and genetic variation of *amh*, *amhy*, and *amhΔy* in Abbassa and GIFT genomes. Similarities and differences in observed genetic variation of the Abbassa and GIFT *amhy* and *amhΔy* genes as compared to *amh.* Observed genetic variation is marked beneath gene structures and further described in Table 1.

## Discussion

Selective breeding has been a major factor in the development of aquaculture, and the availability of high-quality genome assemblies with comprehensive annotations is at the core of enhancing genomic selection of important traits such as environmental resilience and pathogen resistance. Whilst high-quality reference genomes are available for key aquaculture species such as the Nile tilapia^8^, they were generated from tissues and cell lines derived from wild individuals with different evolutionary history to the elite strains. Those elite lines have a complex breeding history involving a mix of wild and farmed populations, with closely related tilapia species also contributing to the gene pool^3^. As such, not only was the sex determination of the GIFT strain suggested to differ from the Nile tilapia UMD reference^19^, our recent work demonstrated extensive introgression with *O. mossambicus* and *O. aureus* during the breeding process, leading to substantial differences in the genomic content of the respective assemblies^10^.

To further enable the selective breeding of the Abbassa strain established in Egypt, we generated a high-quality assembly and associated annotation, that is of a similar quality to the one we recently generated for the GIFT strain^10^. The observation of large variation between the Abbassa genome assembly and the UMD reference, both in terms of gene content (40,301 vs 33,162 genes for Abbassa and UMD respectively), but also genomic content where we detected introgression with *O. aureus*, stresses the importance of generating strain specific genome assemblies.

Given the complex sex-determining system in Nile tilapia, where males are heterogametic (XX/XY), and controlled by both environmental and genetic factors^24^, it is important for the continued success of genomic selection programmes to characterise the genetic loci likely associated with sex determination and differentiation. It was previously shown that sex determination in the reference UMD strain was associated with LG1^20^, whereas LG23 was identified as the major loci in GIFT^19^. Here, we report the identification of *amhy* and *amhΔy* on the Abbassa scaffold (scaffold 1) homologous to the GIFT LG23. Both Abbassa and GIFT share the deletion events reported in the Nile tilapia founding strain from Egypt^14^, including the 5608 bp deletion within the promoter of *amhy* that is conserved in different Nile tilapia strains and associated with sex determination^14,23,25,2614,19,23^. This has been suggested to occur through earlier and higher expression of *amhy* in XY gonads during sex determination^14,19,23^. Whilst we did not perform association analyses to confirm the direct association of the *amhy* loci to sex determination in Abbassa, we identified high sequence identity between Abbassa and GIFT in these loci, where bulk segregant^19^ and molecular profiling^14,19,23^ analyses performed in multiple GIFT populations identified direct associations of these loci to sex differentiation. Taken together, this strongly suggests that Abbassa shares the same sex determination system as GIFT, as well as certain Nile tilapia populations and strains^14,19,23^.

Sex reversal only occurs in the presence of *amhy* mutations in XY Nile tilapia fish^14,17,19,23^. Whilst the missense SNP in exon II of *amhy*, was previously proposed to be key in male determination in Egyptian Nile tilapia^14^, it was more recently found to be absent in Nile tilapia strains where *amhy* was identified as the sex determiner^27,28^. Since this missense SNP is a polymorphism existing at different frequencies in Nile tilapia populations/strains^14,19,23^, and absent in the Abbassa and GIFT *amhy* genes, other genetic variations in *amhy* and possibly, *amhΔy* could determine male sex in Abbassa or GIFT^19,27^. This could include the 5608 bp *amhy* promoter deletion, previously associated with sex determination^14,19,23^, or the conserved missense SNPs in exon II and exon III of Abbassa and GIFT *amhΔy,* or variation that differs between the two strains, like the missense SNPs in different regions of the TGF-β binding domain of *amhy* exon VII. Since members of the transforming growth factor beta (TGF-β) signalling pathway e.g., *amhy* are associated with sex determination in several teleost fish^29–31^, including Nile tilapia^14^, we suggest that much like that proposed for the GIFT strain^19^, *amhy* and either of its tandemly duplicated copies (*amhy* and *amhΔy*) are likely to be the candidate genes for male sex determination in the Abbassa strain. However, no direct efforts were made to confirm this, and further investigations are required to confirm whether the identified genetic variations are responsible for altered *amh* and *amhy* expression during sex determination.

## Methods

### Tissue collection and HMW DNA extraction

A single male Abbassa individual (Sample ID: A6, Pit tag ID: 00070B13FA) was euthanized by immersion in a solution of clove oil (400 mg/litre) at WorldFish (Egypt). The work was performed under ethical approval of the Universiti Kebangsaan Malaysia Animal Ethics Committee to WorldFIsh, and in accordance with the Guiding Principles of the Animal Care, Welfare and ethics Policy of the World Fish Centre (WorldFish, 2019).

Several tissues were dissected, submerged in absolute ethanol and then flash frozen at − 80 °C. High molecular weight (HMW) DNA was extracted from 7 mg of male gonad tissue using the Circulomics Nanobind Tissue Big DNA Kit. The extraction method was based on the Dounce method described in kit handbook v.1.0. Due to the high DNA content of gonad tissue, the input was reduced from the recommended 25 mg to 7 mg. To minimise viscosity, the DNA was eluted in 300uL EB and left at room temperature for 72 hours. The sample was pipette mixed twice daily using a wide-bore tip. Following this, any remaining jelly-like material was removed using a pipette tip.

### Library Preparation PacBio HiFi

13 µg aliquots of HMW DNA the sample were manually sheared with the Megaruptor 3 instrument (Diagenode, P/N B06010003) according to the Megaruptor 3 operations manual. Each sheared DNA underwent AMPure® PB bead (PacBio®, P/N 100-265-900) purification and concentration before undergoing library preparation using the SMRTbell® Express Template Prep Kit 2.0 (PacBio®, P/N 100-983-900). HiFi libraries were prepared according to the HiFi protocol version 03 (PacBio®, P/N 101-853-100) and libraries were size-fractionated using the SageELF® system (Sage Science®, P/N ELF0001), 0.75% cassette (Sage Science®, P/N ELD7510). The resulting libraries were quantified by fluorescence (Invitrogen Qubit™ 3.0, P/N Q33216) and the size of fractions and libraries was estimated from a smear analysis performed on the FEMTO Pulse® System (Agilent, P/N M5330AA).

The loading calculations for sequencing were performed using the PacBio SMRTlink Binding Calculator v10.1.0.125432 and prepared for sequencing applicable to the library type. Sequencing primers v2 were annealed to the HiFi libraries which were complexed to the sequencing polymerase with the Sequel Binding Kit v2.0 (PacBio, 101-842-900. Sequencing internal control complex 1.0 (PacBio, 101-717-600) was spiked into the final complex preparation at a standard concentration. The libraries were sequenced on the Sequel IIe instrument with Sequel II SMRT®cell 8M cells, chemistry Sequel® II Sequencing Plate 2.0 (PacBio®, 101-820-200) and the Instrument Control Software v10.1. The parameters for sequencing were diffusion loading, 30-hour movies per cell, 2-hour immobilisation time, 4-hour pre-extension time, and 65-70pM on plate loading concentration.

### Omni-C

A total of 20 mg of brain tissue was collected from the same individual (Sample ID: A6, Pit tag ID: 00070B13FA) selected for DNA extraction and sequencing above and stored in ethanol at -80°C. Immediately prior to starting the protocol, it was briefly (30 seconds) immersed in chilled water to remove excess ethanol and blotted dry. The Omni-C library was prepared using the Dovetail® Omni-C® Kit (SKU: 21005) according to the manufacturer’s protocol (https://dovetailgenomics.com/wp-content/uploads/2021/09/Omni-C-Protocol_Non-mammal_v1.0.pdf).

Briefly, the chromatin was fixed with disuccinimidyl glutarate (DSG) and formaldehyde in the nucleus. The cross-linked chromatin was then digested in situ with DNase I (1µl). Following digestion, the cells were lysed with SDS to extract the chromatin fragments and the chromatin fragments were bound to Chromatin Capture Beads. Next, the chromatin ends were repaired and ligated to a biotinylated bridge adapter followed by proximity ligation of adapter-containing ends. After proximity ligation, the crosslinks were reversed, the associated proteins were degraded, and the DNA was purified then converted into a sequencing library (NEBNext^®^ Ultra™ II DNA Library Prep Kit for Illumina^®^ (E7645)) using Illumina-compatible adaptors (NEBNext® Multiplex Oligos for Illumina® (Index Primers Set 1) (E7335)). Biotin-containing fragments were isolated using streptavidin beads prior to PCR amplification.

Test sequencing of the library to establish quality and complexity was carried out using an Illumina MiSeq v2 300 cycle kit (MS-102-2002). As part of a duplex pool it generated 2.06M clusters PF (95%) with 96.65% bases >=Q30 and the data was converted from BCL to fastq format using bcl2fastq-2.20.0. Following data quality control analysis, the same pool was sequenced over two lanes of a NovaSeq 6000 SP flow cell with 300 cycle, v1.5 chemistry kit (20028400) with standard workflow loading, generating 463 million paired-end reads.

### Genome assembly

We generated 31 gigabases (approx. 28x coverage) of Pacific Biosciences HiFi data and 140 gigabases (approx. 127x coverage) of Omni-C data from the Abbassa strain of *Oreochromis niloticus*. We used hifiasm (version 0.16.1^32^) to assemble the HiFi data, using the Omni-C data for phasing. Using the ArimaGenomics mapping pipeline (https://github.com/ArimaGenomics/mapping_pipeline) (commit b2b2457), we used bwa (v0.7.17) to map the R1 and R2 Omni-C reads independently to the hifiasm hic.p.ctg assembly and filtered reads that were chimeric or that were marked as PCR duplicates Salsa2^33^ (version ed76685) was then used with the Arima QCd reads to break contigs in regions identified as misjoins, and join contigs into scaffolds where evidence from paired-end reads suggests should be contiguous (Figure 6).

**Figure 6.**
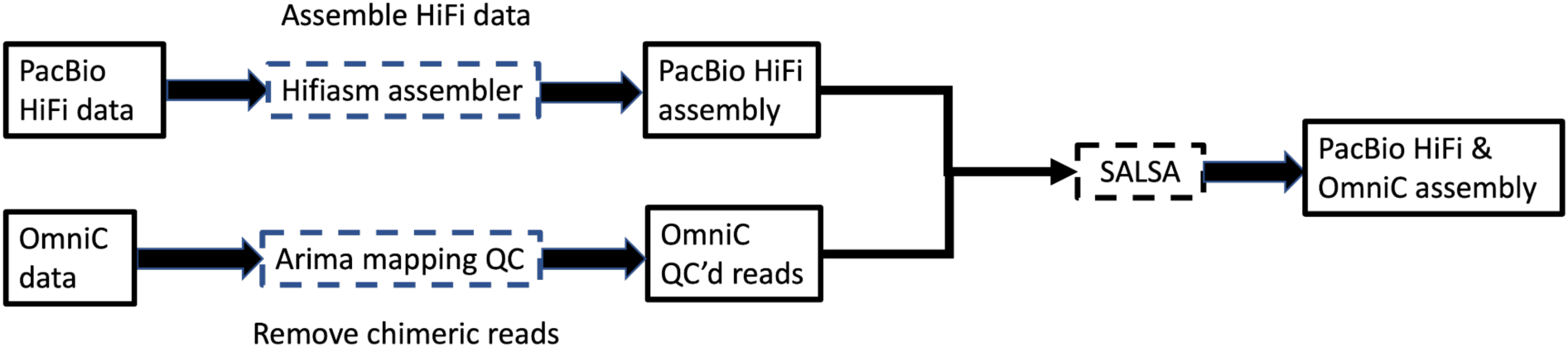
Assembly pipeline for the *O. niloticus* Abbassa genome. Solid squares represent data and dotted lines represent software steps.

### Assembly Quality Control (QC)

We used a number of methods to quality control the assembly.

Assembly stats - we calculated a number of assembly metrics from the reference genome, including assembly size, contig N50, scaffold N50, and number of scaffolds over given lengths.

Genome completeness can also be assessed by counting the number of single-copy orthologs recovered in the assembly that are specific to the clade being sequenced. BUSCO^34^ uses a range of tools, along with an organism-specific ortholog database (orthoDB) to assess the completeness of genomes. We ran BUSCO (v5.0.0) using the actinopterygii_odb10 database on our Abbassa assembly to identify the number of complete and single-copy orthologs, duplicated orthologs, fragmented orthologs, and missing orthologs.

Genome content can be assessed by comparing k-mers (sequences of length ‘k’) in the assembly to those in the reads used to generate it. By comparing the k-mer content of genome assembly to the reads used to assemble it, we can detect both missing and duplicated content in the genome assembly. KAT^35^ (v2.4.1, https://github.com/TGAC/KAT) was used to compare k-mers in the Abbassa genome assembly to those in the Abbassa Omni-C reads. A plot of the k-mer spectra was then generated to visualise the results.

To check for contamination, we used NCBI blastn (v2.8) to compare each scaffold in the Hifiasm assembly to the nt database. The first hit below the default e-value threshold was taken and the taxaid was noted. We then compiled the taxonomic lineages for each taxaid and investigated any scaffold that did not fall at least within the Teleostei (the largest group of ray-finned fishes, of which *Oreochromis* is a member). We found one such scaffold and using NCBI blastn, we extracted all taxaids for each hit below the default e-value and complied the taxanomic lineages for each taxaid. From these, we found that 94.5% of the hits belonged to Teleostei, so retained the scaffold in assembly.

### Comparison with other *Oreochromis* assemblies

We compared our Abbassa assembly with that of the Genetically Improved Farmed Tilapia^10^ (GIFT) assembly and that of the reference *O. niloticus* (Ensembl v110, hereafter UMD) reference sequence ^8^. We used the same QC methods as described above, along with synteny plots to align our assembly against both the GIFT and UMD assemblies using Mummer^36^ (v4.0.0).

### Genome annotation

#### RNA extraction

Four tissues (G: Gills, Go: Gonads, H: Heart, and K: Kidney) were collected from the same individual (Sample ID: A6, Pit tag ID: 00070B13FA) selected for DNA extraction and sequencing above, and stored in RNAlater at -80°C . RNA was extracted from 10-15 mg of tissue input using the Omega EZNA Total RNA Kit I. Homogenization was carried out using a GenoGrinder set to 1000rpm for 5 minutes. The samples were processed in batches of eight, grouped by tissue type so that the different tissue-specific issues arising during extraction could be addressed separately. These issues primarily concerned filter-clogging, which necessitated additional spin time, and loose tissue pellets, which required special care taken when removing the supernatant. To ensure that small RNAs were captured too, the step following homogenization was performed using 100% ethanol rather than 70%. The optional on-column DNase treatment was carried out as specified in the kit’s user manual.

#### PacBio Iso-Seq

The libraries were constructed using 220-270 ng of total RNA. Reverse transcription cDNA synthesis was performed using NEBNext® Single Cell/Low Input cDNA Synthesis & Amplification Module (NEB, E6421) Each cDNA sample was amplified with barcoded primers for a total of 14 cycles. The barcoded cDNA samples were pooled equimolar before SMRTbell library construction. The library pool was prepared according to the guidelines laid out in the Iso-Seq protocol version 02 (PacBio, 101-763-800), using SMRTbell express template prep kit 2.0 (PacBio, 102-088-900). The library pool was quantified using a Qubit Fluorometer 3.0 (Invitrogen) and sized using the Bioanalyzer HS DNA chip (Agilent Technologies, Inc.).

The loading calculations for sequencing were performed using the PacBio SMRTlink Binding Calculator v10.1.0.125432 and prepared for sequencing applicable to the library type.

Sequencing primers v4 were annealed to the Iso-Seq library pool which was complexed to the sequencing polymerase with the Sequel II binding kit V2.1 (PacBio, 101-843-000).

Sequencing internal control complex 1.0 (PacBio, 101-717-600) was spiked into the final complex preparation at a standard concentration. The libraries were sequenced on the Sequel IIe instrument with Sequel II SMRT®cell 8M, chemistry Sequel® II Sequencing Plate 2.0 (PacBio®, 101-820-200) and the Instrument Control Software v10.1. The parameters for sequencing were, diffusion loading, 30-hour movie, 2-hour immobilisation time, 2-hour pre-extension time, 80pM on plate loading concentration. We generated 3 million reads of PacBio IsoSeq with a mean read-length of 2.66 kb.

#### Illumina RNA-Seq

The libraries were constructed using the NEBNext Ultra II RNA Library prep for Illumina kit (NEB#E7760L), NEBNext Poly(A) mRNA Magnetic Isolation Module (NEB#7490) and NEBNext Multiplex Oligos for Illumina® (96 Unique Dual Index Primer Pairs) (E6440S) at a concentration of 10 µM.

A total of 1 µg of RNA was purified to extract mRNA with a Poly(A) mRNA Magnetic Isolation Module. Isolated mRNA was then fragmented for 12 minutes at 94°C, and first strand cDNA was synthesised followed by second strand synthesis. NEBNext Adaptors were ligated to end-repaired, dA-tailed DNA. The ligated products were subjected to a bead-based purification using Beckman Coulter AMPure XP beads (A63882) to remove un-ligated adaptors. Adaptor-ligated DNA was then enriched using 10 cycles of PCR (30 secs at 98°C, 10 cycles of: 10 secs at 98°C _75 secs at 65°C _5 mins at 65°C, final hold at 4°C). The quality of the resulting libraries was determined using Agilent High Sensitivity DNA Kit from Agilent Technologies (5067-4626) and the concentration measured with a High Sensitivity Qubit assay from ThermoFisher (Q32854). The final libraries were equimolar pooled, q-PCR was performed using a Kapa library quant kit from Roche (7960204001), and the pool was sequenced on the NovaSeq on one lane of a SP flow cell with 300 cycle, v1.5 chemistry kit (20028400) using a 2-lane XP kit (20043130) producing a total of 544 million paired-end reads.

#### Annotation

Gene models were annotated via the Robust and Extendable eukaryotic Annotation Toolkit (REAT v0.6.1, https://github.com/EI-CoreBioinformatics/reat). The REAT pipeline enables genome annotation through the combination of multiple sources of evidence including transcriptome (short and long read, REAT transcriptome) as well as protein sequences from related species (REAT homology). Predictions from the transcriptome and homology workflow are then consolidated using Minos (v1.8.0 https://github.com/EI-CoreBioinformatics/minos). After repeat annotation using EIREPEAT (v1.1.0 https://github.com/EI-CoreBioinformatics/eirepeat), the REAT transcriptome workflow was run with RNA-seq (544 million read pairs) and IsoSeq reads (2.5 million full-length, non-concatemer reads) from four samples. Illumina RNA-seq reads were mapped to the genome with HISAT2^37^ (v2.1.0) and high-confidence splice junctions identified by Portcullis^38^ (v1.2.4). The aligned reads were assembled for each tissue with StringTie2^39^ (v2.1.1) and Scallop^40^ (v0.10.2), IsoSeq transcripts were aligned with minimap2^41^ (v2.24). From the combined set of RNA-seq assemblies and IsoSeq alignments a filtered set of non-redundant gene-models were derived using Mikado^42^ (v2.3.4).

The REAT homology workflow was used to generate gene models based on alignment of proteins from nine cichlid species (*O. niloticus*: GCF_001858045.2^20^; *O. aureus*: GCF_013358895.1^17^; *Maylandia zebra zebra mbuna*: GCF_000238955.4^43^; *Pundamilia nyererei*: GCF_000239375.1^44^; *Neolamprologus brichardi*: GCF_000239395.1^44^; *Archocentrus centrarchus*: GCF_007364275.1^45^; *Haplochromis burtoni*: GCF_018398535.1^46^; *Astatotilapia calliptera*: GCF_900246225.1^44^; *Simochromis diagramma*: GCF_900408965.1). These together with the transcriptome derived models were used to train the AUGUSTUS^47^ (v3.4.0) gene predictor, with transcript and protein alignments plus repeat annotation provided as hints in evidence guided gene prediction using the REAT prediction workflow. Four alternative AUGUSTUS gene builds were generated using different evidence inputs or weightings. Gene models from *Oreochromis niloticus* (Nile tilapia) and *Oreochromis aureus* (blue tilapia) were projected onto the *O.niloticus* (abbassa) assembly.

Minos (v1.8.0) was run to generate a consolidated set of gene models from the transcriptome, homology, projected and AUGUSTUS predictions, the pipeline utilizes external metrics to assess how well supported each gene model is by available evidence, based on these and intrinsic characteristics of the gene models a final set of models was selected. For each gene model a confidence and biotype classification were determined based on the type and extent of supporting data.

## Functional annotation

All proteins were annotated using the eifunannot pipeline (v1.4.0) incorporating AHRD^48^ (v.3.3.3, https://github.com/groupschoof/AHRD) (Supplementary Material 1). Sequences were blasted against the reference proteins (combination of Oniloticus.GCF_001858045.2.protein.fa (NCBI Annotation Release 104) and Oaureus.GCF_013358895.1.protein.fa (NCBI Annotation Release 101)) and the UniProt Vertebrata sequences (data download date 15 June 2022), both Swiss-Prot and TrEMBL datasets^49^. Proteins were BLASTed (v2.6.0; blastp) with an e-value of 1e-5. Interproscan^50^ (v5.22.61) results were also provided to AHRD. The standard AHRD example configuration, distributed with the AHRD tool, was utilised with the following modifications in addition to the location of input and output files:

1. we included the GOA mapping from uniprot (ftp://ftp.ebi.ac.uk/pub/databases/GO/goa/UNIPROT/goa_uniprot_all.gaf.gz) as parameter ’gene_ontology_result’,
2. we also included the interpro database (ftp://ftp.ebi.ac.uk/pub/databases/interpro/61.0/interpro.xml.gz) and provided as parameter ’interpro_database’,
3. we changed the parameter ’prefer_reference_with_go_annos’ to ’false’ and did not use the parameter ’gene_ontology_result’
4. The regular expression used to analyse the reference protein fasta header was amended to the custom header format:

fasta_header_regex: "^>(?<accession>(NP_|XP_|YP_)\\d+(\\.\\d+)?)\\s+\\|[^\\|]+\\|\\s+(?<description<[^\\|]+)(\\s*\\|.*)?$

### Repeat annotation of LG3

Using the Mummer delta-filter (v4.0.0), we extracted Abbassa scaffolds that had at least 50kb of sequence which aligned with at least 95% identity to GIFT LG3. This provided us with 13 scaffolds totalling 128.5Mb. We then used RepeatMasker^51^ (v4.0.8) to identify repeat families in these 13 scaffolds, along with LG3 from both GIFT and UMD.

### Analysis of introgression

To quantify genome-wide introgression between different tilapia species and the Abbassa strain, we calculated D and f4 statistics, and used these to calculate *f*-branch to identify specific introgression events.

First, we mapped reads of a single specimen of abbassa, GIFT, wild-type *O. niloticus* from lake Albert, *O. aureus*, *O. mossambicus, O. urolepis* and *M. zebra* against the Abbassa assembly. All reads except the Abbassa (as these were long reads) were trimmed for adapters and poly-G tails using fastP^52^ (v0.20.0), before being mapped against the reference using bwa mem^53^ (v0.7.17). The PacBio Abbassa reads were mapped using minimap2^41^ (v2.24-41122). The average mapping quality per sample per scaffold was calculated. SNPs were then called against the 23 longest scaffolds from the Abbassa assembly using bcftools^54^ (v1.10.2) using bcftools mpileup and bcftools call with multi-allelic mode. SNPs within 3 base-pairs of or overlapping an indel, with total depth less than 100 or over 900, with more than 1 missing sample, genotype quality less than 30 or minor allele count less than 3 were filtered. The average genotype quality per scaffold was calculated.

In order to get a guide phylogenetic tree for *D* and f4 statistics, the SNP set was filtered to only include sites with at least one homozygous alternate and reference allele, and lightly filtered for linkage disequilibrium (r2 > 0.9 over 20kb windows). A phylogenetic tree was then inferred using iqtree^55^ (v2.0) with automated model detection^56^, with ascertainment bias correction, with 1000 rapid bootstraps and 5 independent runs. The resultant tree was rooted with *M. zebra* and used as a guidetree for Dsuite^57^ (v0.4_r43), with Dtrios run with a block jackknife over 1000 SNPs. Fbranch statistics were calculated using the Dtrios Fbranch function, and plotted using the Dsuite dtools python script.

To investigate phylogenetic relationships across the genome, we used a topology weighting analysis, using TWISST (v0.20.0)^58^. SNPs were first phased and imputed using beagle^59^ (v4.1), and then converted to ‘geno’ format, using genomics_general. Phylogenetic trees were then inferred across the 23 longest scaffolds in sliding windows of 50 SNPs (overlap 10 SNPs, minimum good quality SNPS per window 40, minimum good sites per individual 40), using iqtree^60^ (v1.6.12) with automated model detection with ascertainment bias correction, using scripts adapted from genomics_general. TWISST^58^ analysis was then carried out to calculate topology weightings for all except the GIFT and *M. zebra* samples. A loess parameter smoothing (span 0.05 of each scaffold) was applied prior to plotting. Across the genome, the weighting of putative introgression tree subtracted from that of the putative species tree was plotted, to highlight introgressed regions.

### Genetic variation of sex-linked genes could determine male sex in Abbassa and GIFT strains

We used tblastn^61^ to search for Nile tilapia *amh* (XP_013130583.1) and its previously described flanking genes^15,28^ (*oaz1* - XP_003451355.1 and *dot1l -* XP_019207253.1) in the Hifiasm phased assemblies of Abbassa and GIFT^10^. After identifying the gene containing scaffolds, the nucleotide sequence of the *amh* locus was extracted using Bedtools^62^ (v2.30.0) and aligned using MUSCLE^63^ (v2.5.1) to identify genetic variation using SNP-sites .

All X-haplotype and Y-haplotype genes of the *amh* locus were annotated using a combination of FGENESH++^64^, BLAST^61^, and WebScipio^65^.

## Declarations

### Ethics approval and consent to participate

This research used tissue samples obtained in 2018 as part of the WorldFish GIFT tilapia Genetic Improvement Program. All fish were managed and sampled in accordance with the Guiding Principles of the Animal Care, Welfare and ethics Policy of the World Fish Centre (WorldFish, 2004; updated 2019).

The sampling was in accordance with the ARRIVE guidelines (https://arriveguidelines.org) for the reporting of animal experiments.

### Consent for publication

Not applicable

### Availability of data and materials

All data, including reads and reference sequence associated with the Abbassa genome assembly can be found in the European Nucleotide Archive under Project accession PRJEB57170

## Authors’ contributions

WH and JAHB designed the study. GJE generated the Abbassa assembly. AC performed the TWISST and short-read based introgression analysis. TM and GJE performed sex determination analysis. AD extracted HMW DNA and RNA. FF, prepared the Omni-C libraries. TB, VK, and SH prepared Illumina sequencing libraries. NI and TB prepared the PacBio IsoSeq, and PacBio HiFi libraries. CW and KG designed the sequencing strategy and oversaw data production. TT and SA prepared the Abbassa tissues. GK and DS generated the Abbassa annotations. The manuscript was prepared by WH, GJE, JAHB, AC, and TM. The authors read and approved the final manuscript.

## Data availability

All data, including reads and reference sequence associated with the Abbassa genome assembly can be found in the European Nucleotide Archive under Project accession PRJEB57170

## Acknowledgments

The author(s) acknowledge the support of the Biotechnology and Biological Sciences Research Council (BBSRC), part of UK Research and Innovation; this research was funded by the BBSRC Core Strategic Programme Grant BB/CSP1720/1 and its constituent work packages (BBS/E/T/000PR9818 and BBS/E/T/000PR9819), and the Core Capability Grant BB/CCG1720/1 and the National Capability at the Earlham Institute BBS/E/T/000PR9816 (NC1 - Supporting EI’s ISPs and the UK Community with Genomics and Single Cell Analysis), BBS/E/T/000PR9811 (NC4 - Enabling and Advancing Life Scientists in data-driven research through Advanced Genomics and Computational Training), and BBS/E/T/000PR9814 (NC 3 - Development and deployment of versatile digital platforms for ‘omics-based data sharing and analysis). The authors would like to acknowledge the Scientific Computing group, as well as support for the physical HPC infrastructure and data centre delivered via the NBI Research Computing group, the technical team managing the Abbassa tilapia genetic improvement program and WorldFish country director Dr Ahmed Nasr-Allah. This work was undertaken as part of, and funded by, the CGIAR Research Program on Fish Agri-Food Systems (FISH), and partly by the CGIAR Research Initiative on Resilient Aquatic Food Systems for Healthy People and Planet, led by WorldFish. These programs were supported by contributors to the CGIAR Trust Fund. The authors would like to acknowledge the technical team managing the Abbassa tilapia genetic improvement program and WorldFish country director Dr Ahmed Nasr-Allah. We thank the Dovetail Genomics staff members for their input during this project.

