## supplementary material 1 for "High-quality genome assembly for the genetically improved Abbassa Nile tilapia enables the reconstruction of X and Y haplotypes": Functional_annotation.ReadMe.txt

# Functional annotation:

All the proteins were annotated using AHRD v.3.3.3 (Hallab et al., 2014; https://github.com/groupschoof/AHRD/blob/master/README.textile). Sequences were blasted against the reference proteins (combination of Oniloticus.GCF_001858045.2.protein.fa (NCBI Annotation Release 104) and Oaureus.GCF_013358895.1.protein.fa (NCBI Annotation Release 101)) and the UniProt Vertebrata sequences (data download date 15 June 2022), both Swiss-Prot and TrEMBL datasets (The UniProt Consortium, 2014). Proteins were BLASTed (v2.6.0; blastp) with an e-value of 1e-5. We have also provided interproscan (v5.22.61; Jones et al., 2014) results to AHRD. We adapted the standard AHRD example configuration file path test/resources/ahrd_example_input_go_prediction.yml, distributed with the AHRD tool, changing the following apart from the location of input and output files:
1. we included the GOA mapping from uniprot (ftp://ftp.ebi.ac.uk/pub/databases/GO/goa/UNIPROT/goa_uniprot_all.gaf.gz) as parameter 'gene_ontology_result',
2. we also included the interpro database (ftp://ftp.ebi.ac.uk/pub/databases/interpro/61.0/interpro.xml.gz) and provided as parameter 'interpro_database',
3. we changed the parameter 'prefer_reference_with_go_annos' to 'false' and did not use the parameter 'gene_ontology_result'
4. The regular expression used to analyse the reference protein fasta header was amended to the custom header format:

    fasta_header_regex: "^>(?<accession>(NP_|XP_|YP_)\\d+(\\.\\d+)?)\\s+\\|[^\\|]+\\|\\s+(?<description>[^\\|]+)(\\s*\\|.*)?$"

Below are a few example fasta headers from the reference proteins that AHRD parses using the above regular expression:

>NP_001266368.2 | Symbols:  | type III iodothyronine deiodinase  |
>XP_003437545.1 | Symbols:  | glucosamine 6-phosphate N-acetyltransferase  |
>YP_003355022.1 | Symbols:  | NADH dehydrogenase subunit 1 (mitochondrion)  |

We also did a separate blast of the annotated proteins (blastp; -max_target_seqs 1, -evalue 1e-5) versus the reference proteins to identify the best blast hit against it and is provided with the functional annotation.


The header descriptions for the functional annotation tsv file is below:

1 #Transcipt                                     - EI transcript ID
2 #Gene                                          - EI gene ID
3 #Confidence                                    - gene confidence class: High - High Confidence, Low - Low Confidence
4 #Biotype                                       - gene biotype classification
5 #Transcript-Blast-Hit-Coverage-By-Reference    - Transcript coverage by reference protein blast hit
6 #Reference-Blast-Hit                           - Reference protein blast hit identifier
7 #Reference-Blast-Hit-Coverage-By-Transcipt     - Reference protein blast hit coverage by transcript
8 #AHRD-Blast-Hit-Accession                      - The Accession of the Protein the assigned description was taken from, by AHRD
9 #AHRD-Quality-Code                             - From AHRD documentation:
                                                   AHRD’s quality-code consists of a three character string, where each character is either ‘*’ if the respective criteria is met or ‘-’ otherwise. Their meaning is explained in the following table:
                                                   Position Criteria
                                                   1        Bit score of the blast result is >50 and e-value is <e-10
                                                   2        Overlap of the blast result is >60%
                                                   3        Top token score of assigned HRD is >0.5

10 #Human-Readable-Description                   - The assigned Human Readable Descriptions by AHRD
11 #Interpro-ID (Description)                    - The Interproscan ID with description derived by AHRD
12 #Gene-Ontology-Term                           - The Gene Ontology terms derived by AHRD for the transcript
