## supplementary table 1 for "High-quality genome assembly for the genetically improved Abbassa Nile tilapia enables the reconstruction of X and Y haplotypes"

**Supplementary Table 1.** Number of BUSCOs identified as complete, duplicated, fragmented or missing in the Abbassa annotation and Abbassa genome assembly

| **Busco Plots (actinopterygii_odb10)** | **Abbassa annotation** | **Abbassa genome** |
| --- | --- | --- |
| Complete (single copy) | 3566 | 3493 |
| Complete (2 copies) | 56 | 53 |
| Complete (3 copies) | 4 | 6 |
| Complete (4+ copies) | 1 | 2 |
| Complete | 3627 | 3554 |
| Duplicated | 61 | 61 |
| Fragmented | 3 | 12 |
| Missing | 10 | 74 |
| Total | 3640 | 3640 |
