## supplementary figure 4 for "High-quality genome assembly for the genetically improved Abbassa Nile tilapia enables the reconstruction of X and Y haplotypes"

**Supplementary Figure 4** – Alignment of GIFT *amh*, *amhy*, and *amhΔy* 5' UTR and gene promoter region. Nucleotide sequence conservation is shaded and 346 bp Abbassa *amh* 5' UTR sequence shown for reference. Green nucleotides on *amhΔy* sequence indicate the 46 bp 5' UTR. The 165 bp deletion in *amhΔy* gene promoter and 3 bp 'TCT' insertion in *amhy* gene promoter marked in red. The deletions at > 1.6 kb in *amhy* and *amhΔy* are marked in green.

```
                                     ....:10810....:10820....:10830....:10840....:10850....:10860
GIFT_amh_LG23                  9990 AGCTGAGCGGCGTCTGACCGGTTACTGTGGGGTCCTGGGCCGGTTGCAGGGTCCAGCAGA
GIFT_amhΔy_h1tg000178l         9730 AGCTGAGCGGCGTCTGACCGGTTACTGTGGGGTCCTGGGCCGGTTGCAGGGTCCAGCAGA
GIFT_amhy_h1tg000178l          9990 AGCTGAGCGGCGTCTGACCGGTTACTGTGGGGTCCTGGGCCGGTTGCAGGGTCCAGCAGA
GIFT_amh_5'UTR                    1 ------------------------------------------------------------

                                     ....:10870....:10880....:10890....:10900....:10910....:10920
GIFT_amh_LG23                 10050 GTGTCAGCGCCTCGCTGTAAAGAACGAGCAGACCCAACATGTTTGCAGTGTCTGCGCGTG
GIFT_amhΔy_h1tg000178l         9790 GTGTCAGCGCCTCGCTGTAAAGAACGAGCAGACCCAACATGTTTGCAGTGTCTGCGCGTG
GIFT_amhy_h1tg000178l         10050 GTGTCAGCGCCTCGCTGTAAAGAACGAGCAGACCCAACATGTTTGCAGTGTCTGCGCGTG
GIFT_amh_5'UTR                    1 ----------------------------------------GTTTGCAGTGTCTGCGCGTG

                                     ....:10930....:10940....:10950....:10960....:10970....:10980
GIFT_amh_LG23                 10110 CGTTTGTGTGGTCAGATCTCTCACAGGATGGGAGTTACATCCTCAGACCTCCCCTTCTCA
GIFT_amhΔy_h1tg000178l         9850 CGTTTGTGTGGTCAGATCTCTCACAGGATGGGAGTTACATCCTCAGACCTCCCCTTCTCA
GIFT_amhy_h1tg000178l         10110 CGTCTGTGTGGTCAGATCTCTCACAGGATGGGAGTTACATCCTCAGACCTCCCCTTCTCA
GIFT_amh_5'UTR                   20 CGTCTGTGTGGTCAGATCTCTCACAGGATGGGAGTTACATCCTCAGACCTCCCCTTCTCA

                                     ....:10990....:11000....:11010....:11020....:11030....:11040
GIFT_amh_LG23                 10170 CCACCTGGGGTCCCCTTTCTGCCAAAATAGCAGCACTGTGTGTCCTTGATGACTGAGATT
GIFT_amhΔy_h1tg000178l         9910 CCACCCGGGGTCCCCTTTCTGCCAAAATAGCAGCACTGTGTGTCCTTGATGACTGAGATT
GIFT_amhy_h1tg000178l         10170 CCACCCGGGGTCCTCTTTCTGCCAAAATAGCAGCACTGTGTGTCCTTGATGACTGAGATT
GIFT_amh_5'UTR                   80 CCACCTGGGGTCCCCTTTCTGCCAAAATAGCAGCACTGTGTGTCCTTGATGACTGAGATT

                                     ....:11050....:11060....:11070....:11080....:11090....:11100
GIFT_amh_LG23                 10230 GTCAGTATTTGAGGTATTTTAACCTGCTTGTGGAGAACATTCTAAAATCAGACAGCAAAC
GIFT_amhΔy_h1tg000178l         9970 GTCAGTATTTGAGGTATTTTAACCTGCTTGTGGAGAACATTCTAAAATCAGACAGCAAAC
GIFT_amhy_h1tg000178l         10230 GTCAGTATTTGAGGTATTTTAACCTGCTTGTGGAGAACATTCTAAAATCAGACAGCAAAC
GIFT_amh_5'UTR                  140 GTCAGTATTTGAGGTATTTTAACCTGCTTGTGGAGAACATTCTAAAATCAGACAGCAAAC

                                     ....:11110....:11120....:11130....:11140....:11150....:11160
GIFT_amh_LG23                 10290 GGGACACGGAGGTAAACAGAAGACGACTTTGGACACACTGAACATCCTTATCTAGCAGAC
GIFT_amhΔy_h1tg000178l        10030 GCGACACGGAGGTAAACAGAAGACGACTTTGGACACACTGAACATCCTTATCTAGCAGAC
GIFT_amhy_h1tg000178l         10290 GCGACACGGAGGTAAACAGAAGACGACTTTGGACACACTGAACATCCTTATCTAGCAGAC
GIFT_amh_5'UTR                  200 GGGACACGGAGGTAAACAGAAGACGACTTTGGACACACTGAACATCCTTATCTAGCAGAC

                                     ....:11170....:11180....:11190....:11200....:11210....:11220
GIFT_amh_LG23                 10350 ACAAACAGGTCCCGGAAAGAAAGTTTCTCACGAACCTTTCATAGAATACACCAGGCTGAA
GIFT_amhΔy_h1tg000178l        10090 ACAAACAGGTCCCAGAAAGAAAGTTTCTCACGAACCTTTCATAGAATACATCAGGCTGAA
GIFT_amhy_h1tg000178l         10350 ACAAACAGGTCTCAGAAAGAAAGTTTCTCACAAACCTTTCATAGAATACACCAGGCTGAA
GIFT_amh_5'UTR                  260 ACAAACAGGTCCCGGAAAGAAAGTTTCTCACGAACCTTTCATAGAATACACCAGGCTGAA

                                     ....:11230....:11240....:11250....:11260....:11270....:11280
GIFT_amh_LG23                 10410 AGATGAAGAGTGCTGTTTAATGTTTCGTGGCTGCAGAGCTAATATGCACGCTGCTTAACT
GIFT_amhΔy_h1tg000178l        10150 AGATGAAGAGTGCCGTTTAAAGTTTCGTGGCTGCAGAGCTAATATGCACGCTGCTTAACT
GIFT_amhy_h1tg000178l         10410 AGATGAAGAGTGCTGTTTAATGTTTCGTGGCTGCAGAGCTAATATGCACGCTGCTTAACT
GIFT_amh_5'UTR                  320 AGATGAAGAGTGCTGTTTAATGTTTC---------------------------------------

                                     ....:11290....:11300....:11310....:11320....:11330....:11340
GIFT_amh_LG23                 10470 TAACCCTCTCGAGGCAGGCGTTGCCGATTTGCAACAGTTAAAAACTAACAACCTGATTAC
GIFT_amhΔy_h1tg000178l        10210 TAACCCTCTCGAGGCAGGCGTTGCCGATTTGCAACAGTTAAAAACTAACAACCTGATTAC
GIFT_amhy_h1tg000178l         10470 TAACCCTCTCGAGGCAGGCGTTGCCGATTTGCAACAGTTAAAAACTAACAACCTGATTAC
GIFT_amh_5'UTR                   29 ------------------------------------------------------------

                                     ....:11350....:11360....:11370....:11380....:11390....:11400
GIFT_amh_LG23                 10530 CCTACATACATATTTTATGAGTC-TTTTTTTTTTTTGATAATTTTCCCCAAATATCAGAT
GIFT_amhΔy_h1tg000178l        10270 CCTACATACATATTTTATGAGTCTTTTTTTTTTTTTGATAATTTTCCCCAAATATCAGAT
GIFT_amhy_h1tg000178l         10530 CCTACATACATATTTTATGAGTCTTTTTTTTTTTTTGATAATTTTCCCCAAATATCAGAT
GIFT_amh_5'UTR                   29 ------------------------------------------------------------

                                     ....:11410....:11420....:11430....:11440....:11450....:11460
GIFT_amh_LG23                 10589 TTTTCCTGTAACTAAAACTTAATTTGAGCCTGAGAGGGTTAAACAAAGAGGACAGTAAAA
GIFT_amhΔy_h1tg000178l        10330 TTTTCCTGTAACTAAAACTTAATTTGAGCCTGAGAGGGTTAAACAAAGAGGACAGTAGAA
GIFT_amhy_h1tg000178l         10590 TTTTCCTGTAACTAAAACTTAATTTGAGCCTGAGAGGGTTAAACAAAGAGGACAGTAGAA
GIFT_amh_5'UTR                   29 ------------------------------------------------------------

                                     ....:11470....:11480....:11490....:11500....:11510....:11520
GIFT_amh_LG23                 10649 CTGGAGGTTTGCTTCCATCACGTCGACAACAAAGCTCAAAACGATTTTAAAGAAACACTT
GIFT_amhΔy_h1tg000178l        10390 CTGGAGGTTTGCTTCCATCACGTCGACAACAAAGCTCAAAACGATTTTAAAGAAACACTT
GIFT_amhy_h1tg000178l         10650 CTGGAGGTTTGCTTCCATCATGTCGACAACAAAGCTCAAAACGATTTTAAAGAAACACTT
GIFT_amh_5'UTR                   29 ------------------------------------------------------------

                                     ....:11530....:11540....:11550....:11560....:11570....:11580
```

```
GIFT_amh_LG23            10709 TGCATTTTGGTAAATCTGCTTACTTGTTGCTTTGGACTGATGAC---TCCTCTTAGAGAG
GIFT_amhΔy_h1tg000178l   10450 TGCATTT-----------------------------------------------------
GIFT_amhy_h1tg000178l    10710 TGCATTTTGGTAAATCTGCTTACTTGTTGCTTTGGACTGATGACTCTTCCTCTTAGAGAG
GIFT_amh_5'UTR              29 ------------------------------------------------------------

                               ....:11590....:11600....:11610....:11620....:11630....:11640
GIFT_amh_LG23            10766 ACTGCTGGCAGAAAGCCTTGAAGGGAAACTTCAGCCACATCCACTGTTTTTCATCTTTCT
GIFT_amhΔy_h1tg000178l   10457 ------------------------------------------------------------
GIFT_amhy_h1tg000178l    10770 ACTGCTGGCAGAAAGCCTTGAAGGGAAACTTCAGCCACATCCACTGTTTTTCATCTTTCT
GIFT_amh_5'UTR              29 ------------------------------------------------------------

                               ....:11650....:11660....:11670....:11680....:11690....:11700
GIFT_amh_LG23            10826 GTTTCTCTAAGCGGGGGATGTCCAACATCAGGCCCAGGGGCTAGAATCACCCAGCAATGA
GIFT_amhΔy_h1tg000178l   10457 ---------------------------------------------------CAGCAATGA
GIFT_amhy_h1tg000178l    10830 GTTTCTCTAAGCGGGGGATGTCCAACATCAGGCCCAGGGGCTAGAATCACCCAGCAATGA
GIFT_amh_5'UTR              29 ------------------------------------------------------------

                               ....:11710....:11720....:11730....:11740....:11750....:11760
GIFT_amh_LG23            10886 CTCCAACCCAGCTCACTTGAAGGATGGCATAAAT--------------------------
GIFT_amhΔy_h1tg000178l   10466 CTCCAACCCAGCTCACTTGAAGGATGGCATAAAT--------------------------
GIFT_amhy_h1tg000178l    10890 CTCCAACCCAGCTCACTTGAAGGATGGCAAAAATTTGCCAGCACCAAAACACCCCTTTCA
GIFT_amh_5'UTR              29 ------------------------------------------------------------

                               ....:11770....:11780....:11790....:11800....:11810....:11820
GIFT_amh_LG23            10920 ------------------TTTGGACTTTTAACTGTATTTTCTTAAATTTTATGGCTTTTC
GIFT_amhΔy_h1tg000178l   10500 ------------------TTTGGACTTTTAACTGTATTTTCTTAAATTTTAAGGCTTTTC
GIFT_amhy_h1tg000178l    10950 CATATATATGTATATATATTTATATACTTATATATATTTATATAGACTGTATATAGTTAC
GIFT_amh_5'UTR              29 ------------------------------------------------------------

                               ....:11830....:11840....:11850....:11860....:11870....:11880
GIFT_amh_LG23            10962 CTGCTAATAAAGAACTCTGCCACATGTTCATGCTACACCAAAGTGATTAACAATTACATG
GIFT_amhΔy_h1tg000178l   10542 CTGCTAATAAAGAACTCTGCCACATGTTCATGCTACACCAAAGTGATTAACAATTACATG
GIFT_amhy_h1tg000178l    11010 TTATT--TTACATACCTTTCTGTTTATGACGGAGATGTACAATTAAAGAAAACTTATGTA
GIFT_amh_5'UTR              29 ------------------------------------------------------------

                               ....:11890....:11900....:11910....:11920....:11930....:11940
GIFT_amh_LG23            11022 ACAAAAAAGTTCCTGTTTTTT-TCACCATC-----------TGCTCCAGAAATTAGT---
GIFT_amhΔy_h1tg000178l   10602 ACAAAAAAGTTCCTGTTTTTTTTCACCATC-----------TGCTCCAGAAATTAGT---
GIFT_amhy_h1tg000178l    11068 CAAAAACAGTGTGTCTTGTTCACCAACAGCGGGCCGATGTATGTTTAAAACATTAACAGG
GIFT_amh_5'UTR              29 ------------------------------------------------------------

                               ....:11950....:11960....:11970....:11980....:11990....:12000
GIFT_amh_LG23            11067 -----------------------------------------------TTTTTTCTGTAAT
GIFT_amhΔy_h1tg000178l   10648 -----------------------------------------------TTTTTTCTGTAAT
GIFT_amhy_h1tg000178l    11128 GGCCCCCGTGGCCGTCGCTCGGGCCCCGAGCCGGCTGCACGAGAGCACTTTCATCCAAAG
GIFT_amh_5'UTR              29 ------------------------------------------------------------

                               ....:12010....:12020....:12030....:12040....:12050....:12060
GIFT_amh_LG23            11080 ATTACACATTTATTTATTTATAATGGAAAATTTCTTTGATTTCAAGATTCAAAGTGTTTA
GIFT_amhΔy_h1tg000178l   10661 ATTACACATTTATTTATTTATGATGGAAAATTTCTTTGATTTCAAGATTCAAAGTGTTTA
GIFT_amhy_h1tg000178l    11188 GTGGCGTCTACAGTTGTACACGATG------------CATTCAAAGAACGAGACAATAGC
GIFT_amh_5'UTR              29 ------------------------------------------------------------

                               ....:12070....:12080....:12090....:12100....:12110....:12120
GIFT_amh_LG23            11140 TTGTCATGTGTCCAAAGAAAAAAGGC-----------------ATTTCTCTGTGCAATGA
GIFT_amhΔy_h1tg000178l   10721 TTGTCATGTGTCCAAAGAAAAAAGGC-----------------ATTTCTCTGTGCAGTGA
GIFT_amhy_h1tg000178l    11236 ATATTAAATATGAGAAAGGAAATGCCAAAAAACGGCTTTACAAATGTCCCTGTGTACAGC
GIFT_amh_5'UTR              29 ------------------------------------------------------------

                               ....:12130....:12140....:12150....:12160....:12170....:12180
GIFT_amh_LG23            11183 AACTTTTGCTTTGCT--------------------------------GTCCACCCACAGA
GIFT_amhΔy_h1tg000178l   10764 AACTTTTGCTTTGCT--------------------------------GTCCACCCACAGA
GIFT_amhy_h1tg000178l    11296 ACCAAGTACTTGACTTGTTTTTGATGTTTTCATTACAGAATAAATAAGATCAGGCATAGT
GIFT_amh_5'UTR              29 ------------------------------------------------------------

                               ....:12190....:12200....:12210....:12220....:12230....:12240
GIFT_amh_LG23            11211 TGCCGGTTAATA--------TTTACAGTAGAAATAGAACAGATAAATACAAAT-------
GIFT_amhΔy_h1tg000178l   10792 TGCCGGTTAATA--------TTTACAATAGAAATAGAACAGATAAATACAAAT-------
GIFT_amhy_h1tg000178l    11356 CCAAACTTAGCACCATTGTTTTCAGACTGTGAGTCAAGTACACACACACTAATGCATTCT
GIFT_amh_5'UTR              29 ------------------------------------------------------------

                               ....:12250....:12260....:12270....:12280....:12290....:12300
GIFT_amh_LG23            11256 --------AGCACAAATAAA----TAAAGACAGAAAAGGAGACAAATTATTGAAGTGTGA
```

```
GIFT_amhΔy_h1tg000178l  10837 ---------AGCACAAATAAA----TAAAGACAGAAAAGGAGACAAATTATTGAAGTGTGA
GIFT_amhy_h1tg000178l   11416 ACAATGTCAACACAATTCAGGATTTAAAAAAAAAAAAAAGGCAAATGGAAAGACCG-AC
GIFT_amh_5'UTR             29 ------------------------------------------------------------

                              ....:12310....:12320....:12330....:12340....:12350....:12360
GIFT_amh_LG23           11304 AACATAGATGTGTGCAAAAGAGCTATAGCTTAATATGCTGGCTTAATATGCAGGATGACC
GIFT_amhΔy_h1tg000178l  10885 AACAGAGATGTGTGCAAAAGAGCTATAGCTTAATATGCTGGCTTAATATGCAGGATGACC
GIFT_amhy_h1tg000178l   11475 AGCATAAATATAATCATCTAAATTATCCACAACTATACAGTCAGAGAATTAATCACATTC
GIFT_amh_5'UTR             29 ------------------------------------------------------------

                              ....:12370....:12380....:12390....:12400....:12410....:12420
GIFT_amh_LG23           11364 TGATGTGCAAAAAATTATTATTAATGTTCAAAAATAGGTCAGCTTGTTTGAGTCTGATGG
GIFT_amhΔy_h1tg000178l  10945 TGATGTGCAAAAAATTATTATTGATGTTCAAAAATAGGTCAGCTTGTTTGAGTCTGATGG
GIFT_amhy_h1tg000178l   11535 ATA-----GTACAAACATCGTTTACAAAAACAAATTACACCGTTTTTTAAAAAGCAAACA
GIFT_amh_5'UTR             29 ------------------------------------------------------------

                              ....:12430....:12440....:12450....:12460....:12470....:12480
GIFT_amh_LG23           11424 CAGTGGGGAAGAAGGCGTTGTTGAGTCTGGATGTTCTGGATTTCACACTTCTAAACCTCC
GIFT_amhΔy_h1tg000178l  11005 CAGTGGGGAAGAAGGCGTTGTTGAGTCTGGATGTTCTGGATTTCACACTTCTAAACCTCC
GIFT_amhy_h1tg000178l   11590 AAGAGGTGTGAAACAAGTCCAGTAGTTTG-----TTGTGGTTATGGTCAACTATGTTTTC
GIFT_amh_5'UTR             29 ------------------------------------------------------------

                              ....:12490....:12500....:12510....:12520....:12530....:12540
GIFT_amh_LG23           11484 GCCCCGAGGGCAGAAGTGTGAACAGTCCGTGTTGGGGATGTGTGGGGTCTTTGAGGATGG
GIFT_amhΔy_h1tg000178l  11065 GCCCCGAGGGCAGAAGTGTGAACAGTCCATGTTGGGGATGTGTGGGGTCTTTGAGGATGG
GIFT_amhy_h1tg000178l   11645 CCAGTGTGTGAGGATTTGCATGCATGACTGTGTGTGTGTGTGTGTGTGTGTTTAAACTAA
GIFT_amh_5'UTR             29 ------------------------------------------------------------

                              ....:12550....:12560....:12570....:12580....:12590....:12600
GIFT_amh_LG23           11544 AGGCAG----------------------CTCTCCTTTGGA--------------------
GIFT_amhΔy_h1tg000178l  11125 AGGCGG----------------------CTCTCCTCTGGA--------------------
GIFT_amhy_h1tg000178l   11705 ACGCAGTAAAATTTCACATATCAGTCCACTTTCATTTAAACCTGCATTTCCTAACAGCAC
GIFT_amh_5'UTR             29 ------------------------------------------------------------

                              ....:12610....:12620....:12630....:12640....:12650....:12660
GIFT_amh_LG23           11562 ----------------------CTCTGCGATGGTAGATGCTGTGCAGAGAGGGCAGCGGA
GIFT_amhΔy_h1tg000178l  11143 ----------------------CTCTGCGATGGTAGATGCTGTGCAGAGAGGGCAGCGGA
GIFT_amhy_h1tg000178l   11765 GGATAAATATGAATGCATGGATCTCTGCCGTTATGCTTAGCTTACAAAGAG---------
GIFT_amh_5'UTR             29 ------------------------------------------------------------

                              ....:12670....:12680....:12690....:12700....:12710....:12720
GIFT_amh_LG23           11600 GTCCTGATTATCTTCCCTGCAGTTGTTATCACTC------------------------TC
GIFT_amhΔy_h1tg000178l  11181 GTCCTGATTATCTTCCCTGCAGTTGTTATCACTC------------------------TC
GIFT_amhy_h1tg000178l   11816 -----GAATATTTCTGCTGCCGTTTGTGTGACTTCACAAACCGACGGGAAAGAAACGGTT
GIFT_amh_5'UTR             29 ------------------------------------------------------------

                              ....:12730....:12740....:12750....:12760....:12770....:12780
GIFT_amh_LG23           11636 TGCAGGTGATTGCAGTCCATGGCAGTAGTGCTGCCGTATCATGCAGTGATGCAGCTGGTC
GIFT_amhΔy_h1tg000178l  11217 TGCAGGTGATTGCGGTCCATGGCAGTAGTGCTGCCATATCATGCAGTGATGCAGCTGGTC
GIFT_amhy_h1tg000178l   11871 AATAAATACACGCAAACTCACTGAACTCTGTTGTTCTTGGACATGTTGATTCAGCTGTTC
GIFT_amh_5'UTR             29 ------------------------------------------------------------

                              ....:12790....:12800....:12810....:12820....:12830....:12840
GIFT_amh_LG23           11696 AGTGTGCTCTCCACAATGCAGCTGTAGAAC-------CTGCTGAGGATCATACCAAATTT
GIFT_amhΔy_h1tg000178l  11277 AGTGTGCTCTCCACAATGCAGCTGTAGAAC-------CTGCTGAGGATCATACCAAATTT
GIFT_amhy_h1tg000178l   11931 GCTGAAC-CTTAACAGAGCAGCCACAGTGACGGGTGGCTGAAGAGCACCATTTCATTTTT
GIFT_amh_5'UTR             29 ------------------------------------------------------------

                              ....:12850....:12860....:12870....:12880....:12890....:12900
GIFT_amh_LG23           11749 CCTCAGCCTCCTCAGGAAATGC-AGCCATTCCTGAGCCTTCTTGAC-----CAGCTGTGT
GIFT_amhΔy_h1tg000178l  11330 CCTCAGCCTCCTCAGGAAATGC-AGCCGTTCCTGAGCCTTCTTGAC-----CAGCTGTGT
GIFT_amhy_h1tg000178l   11990 TTTAAAACAGATGTTAATGTACAAGACGCTGCATTGCATTATTGGCACCCACAGGCATGT
GIFT_amh_5'UTR             29 ------------------------------------------------------------

                              ....:12910....:12920....:12930....:12940....:12950....:12960
GIFT_amh_LG23           11803 GGT--GTTCAGTGTCCAGGTGATGTCCACAGAAATGTAGACGCCCAGGTATCTAAAGCTG
GIFT_amhΔy_h1tg000178l  11384 GGT--GTTCAGTGTCCAGGTGATGTCCACAGAAATGTAGACGCCCAGGTATCTAAAGCTG
GIFT_amhy_h1tg000178l   12050 TATTAATTTATCCTCAAAATAAGGCCAAAATCAACATCCGCAGATGTTTTCCTCATTCCT
GIFT_amh_5'UTR             29 ------------------------------------------------------------

                              ....:12970....:12980....:12990....:13000....:13010....:13020
GIFT_amh_LG23           11861 CTCACCCTCTCCACTTCAAGGCCCCGGATAAACCGCGGCTGGTGAGGCCTCCTCTTCTTC
GIFT_amhΔy_h1tg000178l  11442 CTCACCCTCTCCACTTCAAGGCCCCGGATAAACCGCGGCTGGTGAGGCCTCCTCTTCTTC
```

```
GIFT_amhy_h1tg000178l   12110 CTCATCCTTT-----------------------------GGGAGATTTGTCTGATTATTG
GIFT_amh_5'UTR             29 ------------------------------------------------------------

                              ....:13030....:13040....:13050....:13060....:13070....:13080
GIFT_amh_LG23           11921 TTCGTGTCCACTATCATCTCCTTTGTCTTGTCAGTGTTGAGGGTGAGGTTGTTGTCCTCA
GIFT_amhΔy_h1tg000178l  11502 TTCGTGTCCACTATCATCT-----------------------------------------
GIFT_amhy_h1tg000178l   12141 CTAACATTAAATATC---------------------------------------------
GIFT_amh_5'UTR             29 ------------------------------------------------------------

                              ....:13090....:13100....:13110....:13120....:13130....:13140
GIFT_amh_LG23           11981 CACCATGACACCAGACCGGCCACCTCTCTCCTGTAAGCCGCTTCGTCCCCTCCAGTGATG
GIFT_amhΔy_h1tg000178l  11521 ------------------------------------------------------------
GIFT_amhy_h1tg000178l   12156 ------------------------------------------------------------
GIFT_amh_5'UTR             29 ------------------------------------------------------------

                              ....:13150....:13160....:13170....:13180....:13190....:13200
GIFT_amh_LG23           12041 CGACGGATCACTGCAGTGCCATCTGAGAACTTCAAAATGATGTTATCTTTGTTGGAGGTG
GIFT_amhΔy_h1tg000178l  11521 ------------------------------------------------------------
GIFT_amhy_h1tg000178l   12156 ------------------------------------------------------------
GIFT_amh_5'UTR             29 ------------------------------------------------------------

                              ....:13210....:13220....:13230....:13240....:13250....:13260
GIFT_amh_LG23           12101 ACACAGTCGTGAGCGTACAGGGTGTAGAGGATGGGACTGAGGACTTTTGACATTACATTA
GIFT_amhΔy_h1tg000178l  11521 ------------------------------------------------------------
GIFT_amhy_h1tg000178l   12156 ------------------------------------------------------------
GIFT_amh_5'UTR             29 ------------------------------------------------------------

                              ....:13270....:13280....:13290....:13300....:13310....:13320
GIFT_amh_LG23           12161 TATATGGAAAAACTGAGACGTAGAGTTGAGGTTGCACTTTCTTATATTAAGATTCACATG
GIFT_amhΔy_h1tg000178l  11521 ------------------------------------------------------------
GIFT_amhy_h1tg000178l   12156 ------------------------------------------------------------
GIFT_amh_5'UTR             29 ------------------------------------------------------------

                              ....:13330....:13340....:13350....:13360....:13370....:13380
GIFT_amh_LG23           12221 TAGTTAAGCTTCTTTACTCTGACAGAGCGGGGCGTTTCACTGGTCAGGCACACTTAAGAC
GIFT_amhΔy_h1tg000178l  11521 ------------------------------------------------------------
GIFT_amhy_h1tg000178l   12156 ------------------------------------------------------------
GIFT_amh_5'UTR             29 ------------------------------------------------------------

                              ....:13390....:13400....:13410....:13420....:13430....:13440
GIFT_amh_LG23           12281 CAAAGTGGGCTGAATGTGGCCCACGATCCACAATCAGTTGGACATCCCTGCTTCAGGCTA
GIFT_amhΔy_h1tg000178l  11521 ------------------------------------------------------------
GIFT_amhy_h1tg000178l   12156 ------------------------------------------------------------
GIFT_amh_5'UTR             29 ------------------------------------------------------------

                              ....:13450....:13460....:13470....:13480....:13490....:13500
GIFT_amh_LG23           12341 ACCATCTCCTGCCCCCGGCTCCCTGTGTAAATGTACGATCACTAGTGACACTACTAAACA
GIFT_amhΔy_h1tg000178l  11521 ------------------------------------------------------------
GIFT_amhy_h1tg000178l   12156 ---------------------------------------------------TGACATTAG
GIFT_amh_5'UTR             29 ------------------------------------------------------------

                              ....:13510....:13520....:13530....:13540....:13550....:13560
GIFT_amh_LG23           12401 CTGTTAAAAGGCTTTTATTCTACTCTAAATAAAAGTCTGATCTGTTTTTATAAGACATAT
GIFT_amhΔy_h1tg000178l  11521 -----------CTTTTATTCTACTCTAAATAAAAGTCTGATCTGTTTTTATAAGACATAT
GIFT_amhy_h1tg000178l   12165 CAGAAAAACAGAAACAACTTAACTTTAAGGAAATAGCTACAAAATATTTA----ATATTT
GIFT_amh_5'UTR             29 ------------------------------------------------------------

                              ....:13570....:13580....:13590....:13600....:13610....:13620
GIFT_amh_LG23           12461 TATGAG-CATATTAAAGAGTAGCGGACCAAGGACAGATCCCTGAGGAACTCCCTCCTGAC
GIFT_amhΔy_h1tg000178l  11570 TATGAG-CATATTAAAGAATAGCGGACCAAGGACAGATCCCTGAGGAA----CTCCTGAC
GIFT_amhy_h1tg000178l   12221 TTTGAGACAGAAGGAGTCAGAGCAAGCCAAA---------CAGACAATCTAATTTATTAT
GIFT_amh_5'UTR             29 ------------------------------------------------------------

                              ....:13630....:13640....:13650....:13660....:13670....:13680
GIFT_amh_LG23           12520 TCAATTTTGATTTGATCAACTAAAACATGAAACCTTCAGTACTTCACCAGT---------
GIFT_amhΔy_h1tg000178l  11625 TCAATTTTGATTTGATCAACTAAAACATGAAACCTTCAGTACTTCACCAGT---------
GIFT_amhy_h1tg000178l   12272 TTGGCTTTGAATTAAGCACCAACAACTCA----TTGCAGTTTGTCACAAATGTAAATATG
GIFT_amh_5'UTR             29 ------------------------------------------------------------

                              ....:13690....:13700....:13710....:13720....:13730....:13740
GIFT_amh_LG23           12571 ---------TAAACCAACTGAACCATGAAGTCTTTGAATGAAAAAGCCAAAATGTGCAAT
GIFT_amhΔy_h1tg000178l  11676 ---------TAAACCAACTGAACCATGAAGTCTTTGAATGAAAAAGCCAAGATGTGCAAT
GIFT_amhy_h1tg000178l   12328 AAAACACAGTAAATACAGTAAATCATGAACATGGCAGGGGTGCCAACTAACACAAGCAAG
```

```
GIFT_amh_5'UTR                 29 ------------------------------------------------------------

                                  ....:13750....:13760....:13770....:13780....:13790....:13800
GIFT_amh_LG23                  12622 GAATTCAAAGCCAGTGTTTGCAAATGCAGTTTTTGAATCGTTATTTTCAGAGTCCAATTT
GIFT_amhΔy_h1tg000178l         11727 GAATTCAAAGCCAGTGTTTGCAAATGCAGTTTTTGAATCGTTATTTTCAGAGTCCAATTT
GIFT_amhy_h1tg000178l          12388 G------------GCTTGTACATCTTCTGCTATGTGAGCTTAATATCCAGTGCGAAGTT-
GIFT_amh_5'UTR                 29 ------------------------------------------------------------

                                  ....:13810....:13820....:13830....:13840....:13850....:13860
GIFT_amh_LG23                  12682 TAAAAAACTGAAAAAATTAACTATTACAGCTCATAAAAACAT-TACAGCAGCACCC----
GIFT_amhΔy_h1tg000178l         11787 TAAAAAACTGAAGAAATTAACTATTACAGCTCATAAAAACAT-TACAGCAGCACCCAGAG
GIFT_amhy_h1tg000178l          12435 ----AGACCAAAAAAGTTCCTCATTTGTTACGCAACCAAATATACTGCAACAAATACAG
GIFT_amh_5'UTR                 29 ------------------------------------------------------------

                                  ....:13870....:13880....:13890....:13900....:13910....:13920
GIFT_amh_LG23                  12737 --------------------------------TAACTATTGCAAACCAGGAAATAATCTG
GIFT_amhΔy_h1tg000178l         11846 TCATTTTAGCATTTCTAAAATTATAAGAAAAGTAACTATTGCAAACCAGGAAATAATCTG
GIFT_amhy_h1tg000178l          12491 ----------------AATTTCACCGGGGGAGGAAATAACGCATGCACACAAAAGCAAAG
GIFT_amh_5'UTR                 29 ------------------------------------------------------------

                                  ....:13930....:13940....:13950....:13960....:13970....:13980
GIFT_amh_LG23                  12765 GTTTATAAGGACTG---------CAGTCTTTTAACACTTTTTATAT--------------
GIFT_amhΔy_h1tg000178l         11906 GTTTATAAGGACTG---------CAGTCTTTTAACACTTTTTATAT--------------
GIFT_amhy_h1tg000178l          12535 GTGCTTGACGTCTGAGCAGAGTTCAATGAACTGGCTCTGTGTGTGTGTCTGAGTGTGTGT
GIFT_amh_5'UTR                 29 ------------------------------------------------------------

                                  ....:13990....:14000....:14010....:14020....:14030....:14040
GIFT_amh_LG23                  12802 -----------TAATGTTTTTAAGTACAAAATATACTGCTATTTGTTTTATTTTGCTTAA
GIFT_amhΔy_h1tg000178l         11943 -----------TAATGTTTTTAAGTACAAAATATACTGCAATTTGTTTTATTTTGCTTAA
GIFT_amhy_h1tg000178l          12595 GACTCGTCAAGTGAAACTTTGGGGGGGGAAGTAGATCCTGGTTTATCCTG--TCGTTAAA
GIFT_amh_5'UTR                 29 ------------------------------------------------------------

                                  ....:14050....:14060....:14070....:14080....:14090....:14100
GIFT_amh_LG23                  12851 TTTTCATTCAGTTAATATCTTTTTCTAATTTTTAGAGAAGCTTTGTGG------------
GIFT_amhΔy_h1tg000178l         11992 TTTTCATTCAGTTAATATCTTTTTCTAATTTTTAGAGAAGCTTTGTGG------------
GIFT_amhy_h1tg000178l          12653 TGATAATTC------TTTTTTTTTCTGATTTTGCTGAATCCCAGTTGCTATAATTGGCA
GIFT_amh_5'UTR                 29 ------------------------------------------------------------

                                  ....:14110....:14120....:14130....:14140....:14150....:14160
GIFT_amh_LG23                  12899 ------AAAAGAAACTGTTCGGGAGCAAAAATATTTCACAGTTCCAATCCG--------CT
GIFT_amhΔy_h1tg000178l         12040 ------AAAAGAAACTGTTCGGGAGCAAAAATATTTCACAGTTCCAATCCG--------CT
GIFT_amhy_h1tg000178l          12707 CCAGAACAAAGAAACTACATATGAACAGATTAGATAGCACCTTAAATTCAAACACAATAT
GIFT_amh_5'UTR                 29 ------------------------------------------------------------

                                  ....:14170....:14180....:14190....:14200....:14210....:14220
GIFT_amh_LG23                  12945 TTAGTTTGTTTCATAACTGAAATGTCATTAATTTATTTTCATCTATGTCAGATGAGAAAT
GIFT_amhΔy_h1tg000178l         12086 TTAGTTTGTTTCATAACTGAAATGTCATTAATTTATTTTCATCTATGTCAGATGAGAAAT
GIFT_amhy_h1tg000178l          12767 TTAAGTTCACTCAGAGCTGGACAACAGCCAGACTGTAAAACACTACTCCACACACAAACA
GIFT_amh_5'UTR                 29 ------------------------------------------------------------

                                  ....:14230....:14240....:14250....:14260....:14270....:14280
GIFT_amh_LG23                  13005 GAATCAGATTGTTCATCAACTGT------------------GTGGTCTGCCTGCAT-TT
GIFT_amhΔy_h1tg000178l         12146 GAATCAGATTGTTCATCAACTGT------------------GTGGTCTGCCTGCAT-TT
GIFT_amhy_h1tg000178l          12827 TGACCAG--TACCTATGAATTCTGCTCAGGGTGACCATAGTGGTGGTTAGTGTGTGTGTT
GIFT_amh_5'UTR                 29 ------------------------------------------------------------

                                  ....:14290....:14300....:14310....:14320....:14330....:14340
GIFT_amh_LG23                  13045 TGGGTGAACTTACCATTAAAGGTCATGGGAAGAGAATGGGTTATCTTAATGAACCCACAC
GIFT_amhΔy_h1tg000178l         12186 TGGGTGAACTTACCATTAAAGGTCATGGGAAGAGAATGGGTTATCTTAATGAACCCACAC
GIFT_amhy_h1tg000178l          12885 TGTGTGTGTGTGTGTGTGTGTGTGTGTGTGTGTGTGTGTGTGTGTGTGAGAGAGAGACAA
GIFT_amh_5'UTR                 29 ------------------------------------------------------------

                                  ....:14350....:14360....:14370....:14380....:14390....:14400
GIFT_amh_LG23                  13105 TGGAGATCTGGAAGCATTCACGCTGGAAGACGAGCAGGAAACATGTTGTCAGAGGGAGAC
GIFT_amhΔy_h1tg000178l         12246 TGGAGATCTGGAAGCATTCACGCTGGAAGACGAGCAGGAAACATGTTGTCAGAGGGAGAC
GIFT_amhy_h1tg000178l          12945 AGATCCCAGTACAGAAGCCGGTCTTGGAGGCGATTGGGTAGTGCAGAGTTTGAAGCACCA
GIFT_amh_5'UTR                 29 ------------------------------------------------------------

                                  ....:14410....:14420....:14430....:14440....:14450....:14460
GIFT_amh_LG23                  13165 CTGAGTGGCTTCGAGGAGCTGTTACTCCCCTCAGCATTCTCACAGCTGCAGGCTTGTTGT
GIFT_amhΔy_h1tg000178l         12306 CTGAGCGGCTTCGAGGAGCTGTTACTCCCCTCAGCATTCTCACAGCTGCAGGCTTGTTGT
GIFT_amhy_h1tg000178l          13005 GTGAATGAGGAGAGTGCCTTGGTCCACACCTGGACATTTGCATAGTT---TATTTAATGT
GIFT_amh_5'UTR                 29 ------------------------------------------------------------
```

```
                                     ....:14470....:14480....:14490....:14500....:14510....:14520
GIFT_amh_LG23                  13225 GCAGACAGTC---------------AAGGTCTCGGAGCCATTTGTGGACGTTTTCCCCCG
GIFT_amhΔy_h1tg000178l         12366 GCAGACAGTC---------------AAGGTCTCGGAGCCATTTGTGGACGTTTTCCCCCG
GIFT_amhy_h1tg000178l          13062 TGAGGTAGTTTTCATCAATTACCTGAAACATTCAAAGACAGACGAGGACATC--------
GIFT_amh_5'UTR                    29 ------------------------------------------------------------

                                     ....:14530....:14540....:14550....:14560....:14570....:14580
GIFT_amh_LG23                  13270 GTGATAAAGCCAAGGTTCACCATACCTTATAAATC---ATCAGTGAGGAATGGCAGGAAA
GIFT_amhΔy_h1tg000178l         12411 GTGATAAAGCCAAGGTTCACCATACCTTATAAATC---ATCAGTGAGGAATGGCAGGAAA
GIFT_amhy_h1tg000178l          13114 ATGAGTGATTCTAAGGCTTCCAGAAATAACACAGCAAAAACACAGACAGCTGATTGAGGA
GIFT_amh_5'UTR                    29 ------------------------------------------------------------

                                     ....:14590....:14600....:14610....:14620....:14630....:14640
GIFT_amh_LG23                  13327 TGTTCATTGTATCAAGGCGGTGTTTGAAG--------------------CACACCAGCCT
GIFT_amhΔy_h1tg000178l         12468 TGTTCATTGTATCAAGGCGGTGTTTGAAG--------------------CACACCAGCCT
GIFT_amhy_h1tg000178l          13174 TCTAAACCCTAACAAGTTCAAAATCAAAGTCTTTACTTTTAATCTAACACCTACTTCTCT
GIFT_amh_5'UTR                    29 ------------------------------------------------------------

                                     ....:14650....:14660....:14670....:14680....:14690....:14700
GIFT_amh_LG23                  13367 GGATGTTTTCAGTCACA------------------CTTGTGGAACCAGGAATTGCACTTT
GIFT_amhΔy_h1tg000178l         12508 GGATGTTTTCAGTCACA------------------CTTGTGGAACCAGGAATTGCACTTT
GIFT_amhy_h1tg000178l          13234 TACCTTGTTCAATCAGACAAACAGCTTGCAGAAGCACATGGAAAACAGGCATCCTGTCTC
GIFT_amh_5'UTR                    29 ------------------------------------------------------------

                                     ....:14710....:14720....:14730....:14740....:14750....:14760
GIFT_amh_LG23                  13410 GATGAAAAAGTA-------------ATCAACATCTACACTCCTGT-------------TT
GIFT_amhΔy_h1tg000178l         12551 GATGAAAAAGTA-------------ATCAACATCTACACTCCTGT-------------TT
GIFT_amhy_h1tg000178l          13294 TCTCACTCCCTACCAGCCTCTCTCGCTCACCCTCTATCCACCCTCAGATTGTTCTCCGCC
GIFT_amh_5'UTR                    29 ------------------------------------------------------------

                                     ....:14770....:14780....:14790....:14800....:14810....:14820
GIFT_amh_LG23                  13444 CTCCCCATCATGTATTCACCTCTCTACAGTAGCACAC----TTAAGTTTTCCAGGTTGGA
GIFT_amhΔy_h1tg000178l         12585 CTCCCCATCATGTATTCACCTCTCTACAGTAGCACAC----TTAAGTTTTCCAGGTTGGA
GIFT_amhy_h1tg000178l          13354 CTTCCCCTCTCCTCTCCCCTTCCCCTCAGCCCCAAACCCTATCTAGGTCGGGAGCTGGGG
GIFT_amh_5'UTR                    29 ------------------------------------------------------------

                                     ....:14830....:14840....:14850....:14860....:14870....:14880
GIFT_amh_LG23                  13500 AAAAGGACAAAAACACAGAAGACATACATTTCTGGGGAATCTCATCTGTCCATGTTGTAA
GIFT_amhΔy_h1tg000178l         12641 AAAAGGACAAAAACACAGAAGACATACATTTCTGGGGAATCTCATCTGTCCATGTTGTAA
GIFT_amhy_h1tg000178l          13414 AGCTGGGAGGTGGTGTAGGTACCATGCATGCTCTGAGTCCTCCACATGCTCCCA------
GIFT_amh_5'UTR                    29 ------------------------------------------------------------

                                     ....:14890....:14900....:14910....:14920....:14930....:14940
GIFT_amh_LG23                  13560 GGTTGTATCCTCATTACC-----CCGAAGCCAAAGGCGGTAACCGGGGAACGCCACAGGC
GIFT_amhΔy_h1tg000178l         12701 GGTTGTATCCTCATTACC-----CCGAAGCCAAAGGCGGTAACCAGGGAACGCCACAGGC
GIFT_amhy_h1tg000178l          13468 ----GCACCACCAGAACTGCTGGTCGGTGGCGGAGGGGGCAGACGGGGGGAG--------
GIFT_amh_5'UTR                    29 ------------------------------------------------------------

                                     ....:14950....:14960....:14970....:14980....:14990....:15000
GIFT_amh_LG23                  13615 CTGGCTTCAGGTGGAAAGCTTCAGTGCACGGACCTTTTGGGAGGCTTTCTCTGAGGCATC
GIFT_amhΔy_h1tg000178l         12756 CTGGCTTCAGGTGGAAAGCTTCAGTGCACGGGCCTTTTGGGAGGCTTTCTCTGAGGCATC
GIFT_amhy_h1tg000178l          13516 ---------------------TAGTGCACTGCAAATGGACGAGA-------GGAGGAGGC
GIFT_amh_5'UTR                    29 ------------------------------------------------------------

                                     ....:15010....:15020....:15030....:15040....:15050....:15060
GIFT_amh_LG23                  13675 ACAGGCATTCGACACAGAATAAGGTAACACGTCTGT--------GAACTGAAGCAGCAGA
GIFT_amhΔy_h1tg000178l         12816 ACAGGCATTCGACACAGAATAAGGTAACACGTCTGT--------GAACTGAAGCAGCAGA
GIFT_amhy_h1tg000178l          13548 AGAGGAGGCTGAGAGAGAGGAAGAGGAGAGGGATGCAGCAGCAGAGACAGAAGCAGCAGA
GIFT_amh_5'UTR                    29 ------------------------------------------------------------

                                     ....:15070....:15080....:15090....:15100....:15110....:15120
GIFT_amh_LG23                  13727 AGCTGTTTGTTTTTACAGACCTCACTGGACCATTAATCCTCAAGTTAGTGACTGAAGCAG
GIFT_amhΔy_h1tg000178l         12868 AGCTGTTTGTTTTTACAGACCTCACTGGACCATTAATCCTCAAGTTAGTGACTGAAGCAG
GIFT_amhy_h1tg000178l          13608 GGTAGAGGATG-----AGATGGAGAGGGATGATTGTGCTCTCTGGTGGTGGAGAGAGGAG
GIFT_amh_5'UTR                    29 ------------------------------------------------------------

                                     ....:15130....:15140....:15150....:15160....:15170....:15180
GIFT_amh_LG23                  13787 CTCTCTTATGCAGGACTACATTATGTTTATTATAACGTTCTCCATTTCTTTTTAAAGCTG
GIFT_amhΔy_h1tg000178l         12928 CTCTCTTATGCAGGACTACAT--TGTTTATTATAACGTTCTCCATTTCTTTTTAAAGCTG
GIFT_amhy_h1tg000178l          13663 -------AGGGACAGACGCGAGGAGT-----------------------------GGAGG
GIFT_amh_5'UTR                    29 ------------------------------------------------------------
```

```
                                   ....:15190....:15200....:15210....:15220....:15230....:15240
GIFT_amh_LG23              13847 ACATATTTTTTAAAACAATGTCATTAATTGTGGTTTTTAATCATCCATCAATTTCATATC
GIFT_amhΔy_h1tg000178l     12986 ACATATTTTTTAAAACAATGTCATTAATTGTGGTTTTTAATCATCCATCAATTTCATATC
GIFT_amhy_h1tg000178l      13687 GAGTCTGATCGAGAAGACGCTGGGGAGTCAGAGTGCGTGGATGTCGATGAGGAGGGGACG
GIFT_amh_5'UTR                29 ------------------------------------------------------------

                                   ....:15250....:15260....:15270....:15280....:15290....:15300
GIFT_amh_LG23              13907 CATCTCTCTGATTCAGGCTTGCAGGAGGGCAGGACCACAGTGAACAGGCTGTTAGTCTTT
GIFT_amhΔy_h1tg000178l     13046 CATCTCTCTGATTCAGGCTTGCAGGAGGGCAGGACCACAGTGAACAGGCTGTTAGTCTTT
GIFT_amhy_h1tg000178l      13747 C--------TATGCAGGGGTGGAGGAGGGGACAGGGAGAGCGTGGGAGGAGAGGGGCT--
GIFT_amh_5'UTR                29 ------------------------------------------------------------

                                   ....:15310....:15320....:15330....:15340....:15350....:15360
GIFT_amh_LG23              13967 CTCAGGGCTGACACACACACACACACACACACACACACACACACCCATCACTAATGATGC
GIFT_amhΔy_h1tg000178l     13106 CTCAGGGCTG------------ACACACACACACACACACACACCCATCACTAATGATGC
GIFT_amhy_h1tg000178l      13797 --------------------------------ATGCAGGACCGACTGCGTGCGGCTGTGC
GIFT_amh_5'UTR                29 ------------------------------------------------------------

                                   ....:15370....:15380....:15390....:15400....:15410....:15420
GIFT_amh_LG23              14027 TGACTTCTCAAAAACTCGAGTATCTTATACAGAGAAGCATTTTTTAATCAATTTATTTTA
GIFT_amhΔy_h1tg000178l     13154 TGACTTCTCAAAAACTCGAGTATCTTATACAGAGACGCATTTTTTAATCAATTTATTTTA
GIFT_amhy_h1tg000178l      13825 TGGGCCTGGGATAACGAGAGGGGCATAGGCGAGGACACG---------------------
GIFT_amh_5'UTR                29 ------------------------------------------------------------

                                   ....:15430....:15440....:15450....:15460....:15470....:15480
GIFT_amh_LG23              14087 CATCGCAGGCCACATGCAGCCCACTTTGAACTTTTGTGAACCGGACCAGTGAAGCTTCCC
GIFT_amhΔy_h1tg000178l     13214 CATCGCAGGCCACATG-----AACTTTGAACTTTTGTGAACCGGACCAGAGAAGCTTCCC
GIFT_amhy_h1tg000178l      13864 -GGCGCCAGGCGCGTG---------------------GAGGCTGGCTGGGGAGTGGTGTG
GIFT_amh_5'UTR                29 ------------------------------------------------------------

                                   ....:15490....:15500....:15510....:15520....:15530....:15540
GIFT_amh_LG23              14147 TTTCTGTCTCTGTGAAGAGGT---------------TTACCTACACATTTGACCTCAGAG
GIFT_amhΔy_h1tg000178l     13269 TTTCTGTCTCTGTGAAGAGGT---------------TTACCTACACATTTGACCTCAGAG
GIFT_amhy_h1tg000178l      13902 TTGCTGGGTTTGGGAGGAGGTAGAGGAAGAGGAGGAGGAGAGACTGAGCAGAGAGTTGAG
GIFT_amh_5'UTR                29 ------------------------------------------------------------

                                   ....:15550....:15560....:15570....:15580....:15590....:15600
GIFT_amh_LG23              14192 ACGTGTGGAAACAGGAAGTGCAGTTTCAACCACATATCTGTTTTTCTACTCAATTAAGAA
GIFT_amhΔy_h1tg000178l     13314 ACGTGTGGAAACAGGAAGTGCAGTTTCAACCACATATCTGTTTTTCTACTCAATTAAGAA
GIFT_amhy_h1tg000178l      13962 GCCTCCTGAGGTGGGGAGCGTTGCCCCAAAC-------------------TGATGGGAGG
GIFT_amh_5'UTR                29 ------------------------------------------------------------

                                   ....:15610....:15620....:15630....:15640....:15650....:15660
GIFT_amh_LG23              14252 GTGCTGCTGTAATGAACTGAAGAAATCTGTCAGCACAAC---------------------
GIFT_amhΔy_h1tg000178l     13374 GTGCTGCTGTAATGAACTGAAGAAATCTGTCAGCACAAC---------------------
GIFT_amhy_h1tg000178l      14003 ACACTGAGGGAACCACGTGGTGAAATATACCTGCAAATCATACAGCAGAAATGGAGGGAG
GIFT_amh_5'UTR                29 ------------------------------------------------------------

                                   ....:15670....:15680....:15690....:15700....:15710....:15720
GIFT_amh_LG23              14291 -------------------------------AGATACTGTGGACCAGCTGTAAAAAACAA
GIFT_amhΔy_h1tg000178l     13413 -------------------------------AGATACTGTGGACCAGCTGTAAAAAACAA
GIFT_amhy_h1tg000178l      14063 GAGGAGGAGGAGGAGGAGGGCGAGAGAGTGGAGACAAGAGAGAGCAGTGGCAAAAAGAGC
GIFT_amh_5'UTR                29 ------------------------------------------------------------

                                   ....:15730....:15740....:15750....:15760....:15770....:15780
GIFT_amh_LG23              14320 ACTGCTATAGTAGATTAAAAAAA-ATTTCTTTAAGGGTTTATATATTTGTTGGTTACTT
GIFT_amhΔy_h1tg000178l     13442 ACTGCTATAGTAGATTAAAAAAAAATTTCTTTAAGGGTTTATATATTTGTTGGTTACTT
GIFT_amhy_h1tg000178l      14123 AGGGGTGGGGGACAGAGAGAGAAG---------CAAGGGCTGA-----------------
GIFT_amh_5'UTR                29 ------------------------------------------------------------

                                   ....:15790....:15800....:15810....:15820....:15830....:15840
GIFT_amh_LG23              14379 TGTCTTTTCATTAGGAAAAACTATCAAACTTGTGG-AAGTTCACATTTTGCCGTCGAGTG
GIFT_amhΔy_h1tg000178l     13502 TGTCTTTTCATTAGGAAAAACTATCAAACTTGTGG-AAGTTCACATTTTGCCGTCGAGCG
GIFT_amhy_h1tg000178l      14157 -GCCAGGTTGTGCAAAGCGCCAAGCAAGTCTGAGACAAGCACACAGAAAGCCAAACAGAGT
GIFT_amh_5'UTR                29 ------------------------------------------------------------

                                   ....:15850....:15860....:15870....:15880....:15890....:15900
GIFT_amh_LG23              14438 GGCCAGATTGGAGTCTTTGCCGGGCCGACTTGGCATTACAAGGTAAATGCCGCCTACCTG
GIFT_amhΔy_h1tg000178l     13561 GGCCAGATTGGAGTCTTTGCCGGGCCGACTTGGCATTACAAGGTAAATGCCGCCTACCTG
GIFT_amhy_h1tg000178l      14216 CAGTCCACTGGAAATGTAACGGAGCCAATTCAGCGCAGCCAGTGTAACGCT---------
GIFT_amh_5'UTR                29 ------------------------------------------------------------

                                   ....:15910....:15920....:15930....:15940....:15950....:15960
```

```
GIFT_amh_LG23            14498 TGAAGTCTACTAACTGAGGATTTAGGTAGCGGGGCTCCAAACATGGTTCAGGTATGAAGA
GIFT_amhΔy_h1tg000178l   13621 TGAAGTCT----ACTGAGGATTTAGGTAGCGGGGCTCCAAACATGGTTCAGGTATGAAGA
GIFT_amhy_h1tg000178l    14267 ----------------------------GCAGGAAGTGAAATGACTCTGATGCACTAACA
GIFT_amh_5'UTR              29 ------------------------------------------------------------

                               ....:15970....:15980....:15990....:16000....:16010....:16020
GIFT_amh_LG23            14558 TATCAGCATGACACCCTCTTCCGTCACGGCAAACCAAATGGACTCAGACAAAGTTCACTC
GIFT_amhΔy_h1tg000178l   13677 TATCAGCATGACACCCTCTTCCGTCACGGCAAACCAAATGGACTCAGACAAAGTTCACTC
GIFT_amhy_h1tg000178l    14299 CACCCGGGGAAAACGCACACGCAGCAGCGTGACCTGCATGAAATCAAAC---GGTCACTC
GIFT_amh_5'UTR              29 ------------------------------------------------------------

                               ....:16030....:16040....:16050....:16060....:16070....:16080
GIFT_amh_LG23            14618 CAAACGTAAACTCAAT-ATTATAAGAAACATTTTATTAATTATCATAAATCGCAGGGGAG
GIFT_amhΔy_h1tg000178l   13737 CAAACGTAAACTCAAT-ATTATAAGAAACATTTTATTAATTATCATAAATCGCAGGGGAG
GIFT_amhy_h1tg000178l    14356 GCGCTTTGTTATTAATGATTTCAATGAAAGTTTTGTGAAAAAT-----------------
GIFT_amh_5'UTR              29 ------------------------------------------------------------

                               ....:16090....:16100....:16110....:16120....:16130....:16140
GIFT_amh_LG23            14677 AAAAAAAAACATTAATTCCATTTTTTCATTATAACATCTTAGGATAAAAGCACCAACATC
GIFT_amhΔy_h1tg000178l   13796 AAAAAAAAACATTAATTCCATTTTTTCATTATAACATCTTAGGATAAAAGCACCAACATC
GIFT_amhy_h1tg000178l    14399 ---------------------------GTCACATTTTCCGAGGGTGTCTCCACCTGCATT
GIFT_amh_5'UTR              29 ------------------------------------------------------------

                               ....:16150....:16160....:16170....:16180....:16190....:16200
GIFT_amh_LG23            14737 AAACAGTAACAATGTTTCCATGAAAAACCGAGGGCGTTTTGGAAAGTAGGTTTGCCATTA
GIFT_amhΔy_h1tg000178l   13856 AAACAGTAACAATGTTTCCATGAAAAACCGAGGGCGTTTTGGAAAGTAGGTTTGCCATTA
GIFT_amhy_h1tg000178l    14432 ACCTAAACAGCCAGTCCACGTCAAAGTATGGCAGCACAGCGAGTCACAGTTTTGC-----
GIFT_amh_5'UTR              29 ------------------------------------------------------------

                               ....:16210....:16220....:16230....:16240....:16250....:16260
GIFT_amh_LG23            14797 TTGCACCTCAAACCTGCCCTGGCTTCACCAATCAGAGAGCACTATACACATCCTGTAGGT
GIFT_amhΔy_h1tg000178l   13916 TTGCACCTCAAACCTGCCCTGGCTTCACCAATCAGAGAGCACTATACACATCCTGTAGGT
GIFT_amhy_h1tg000178l    14487 -----CTGCTCACCTGTCCTTGCTGCCC--------------------------------
GIFT_amh_5'UTR              29 ------------------------------------------------------------

                               ....:16270....:16280....:16290....:16300....:16310....:16320
GIFT_amh_LG23            14857 CACGCGTGTCCTTGAATGTTACCCCCAATTCAAAGAGGGATTCAAACAAGACACCGAACC
GIFT_amhΔy_h1tg000178l   13976 CACGCGTGTCCTTGAATGTTACCCCCAATTCAAAGAGGGATTCAAACAAGACACCGAACC
GIFT_amhy_h1tg000178l    14510 ------------------TCGGGCAAAGCTCCAGGAGGGATTTCAAC---GTACAGGTTT
GIFT_amh_5'UTR              29 ------------------------------------------------------------

                               ....:16330....:16340....:16350....:16360....:16370....:16380
GIFT_amh_LG23            14917 TCCTCCTTAGAGACGCAAATCTAGACAGAGGCATTTGTGAGATGAGGAAGATGCTGAAA-
GIFT_amhΔy_h1tg000178l   14036 TCCTCCTTAGAGACGCAAATCTAGACAGAGGCATTTGTGAGATGAGGAAGATGCTGAAA-
GIFT_amhy_h1tg000178l    14549 CCTCCTTTCAGCGCCGCCCTCCAGACACAGTGCTTTCCTTCAGAGCGCCGGGGCTCAAAG
GIFT_amh_5'UTR              29 ------------------------------------------------------------

                               ....:16390....:16400....:16410....:16420....:16430....:16440
GIFT_amh_LG23            14976 -----------------GTCGTTGCCATTTCACAGCTTCTCGAGATCAGATCGCTCATAA
GIFT_amhΔy_h1tg000178l   14095 -----------------GTCGTTGCCATTTCACAGCTTCTCGAGATCAGATCGCTCATAA
GIFT_amhy_h1tg000178l    14609 AGTAAGAAGCGCACTCTGCTATTGGCAGGGGAAGGCTCCTCAAGAACCAGCACCTGAGCA
GIFT_amh_5'UTR              29 ------------------------------------------------------------

                               ....:16450....:16460....:16470....:16480....:16490....:16500
GIFT_amh_LG23            15019 AGCTCGGACTGAT------------TGTCGGGGGTGGGGGTGTACGTGCCGGCAGGACTG
GIFT_amhΔy_h1tg000178l   14138 AGCTCGGACTGAT------------TGTCGGGGGTGGGGGTGTACGTGCCGGCAGGACTG
GIFT_amhy_h1tg000178l    14669 TCCTGGGAAAGTGTAGTTTAGGTTATGTTAAAAATGAAACTGCACATACCAGCAAGGACA
GIFT_amh_5'UTR              29 ------------------------------------------------------------

                               ....:16510....:16520....:16530....:16540....:16550....:16560
GIFT_amh_LG23            15067 TGTCTGAGGAGATCTGATGAGGGCAGCTGATACATTTTACCTTGAAATACACTGACTGAA
GIFT_amhΔy_h1tg000178l   14186 TGTCTGAGGAGATCTGATGAGGGCAGCTGATACATTTTACCTTGAAATACACTGACTGAA
GIFT_amhy_h1tg000178l    14729 ATTCTG----TATTTCTTTTTAAGCTTTCATCACTTACAGAATAGAACAGCTTAGCTGAA
GIFT_amh_5'UTR              29 ------------------------------------------------------------

                               ....:16570....:16580....:16590....:16600....:16610....:16620
GIFT_amh_LG23            15127 GGGTGGC--------------------------CCTGGTCCTGAGTTGGGTCGTAGAGGG
GIFT_amhΔy_h1tg000178l   14246 GGGTGGC--------------------------CCTGGTCCTGAGTTGGGTCGTAGAGGG
GIFT_amhy_h1tg000178l    14785 AGATTGCGATCCCGTTGGTCATTCCCTCGCCCACCTGGGATCTCCAGGGGTGGGAGAGGG
GIFT_amh_5'UTR              29 ------------------------------------------------------------

                               ....:16630....:16640....:16650....:16660....:16670....:16680
GIFT_amh_LG23            15161 ACTTTGT--------CATGCCACCTGAAGGTAAATTAACGTAACA--TGCAGCAGATTAG
```

```
GIFT_amhΔy_h1tg000178l  14280 ACTTTGT--------CATGCCACCTGAAGGTAAATTAACGTAACA--TGCAGCAGATTAG
GIFT_amhy_h1tg000178l   14845 GCATCAGGAGCACCACAGAGGCCCCGGACACGGGTTACTGCAACAGCGGGAGCACCTCAG
GIFT_amh_5'UTR             29 ------------------------------------------------------------

                              ....:16690....:16700....:16710....:16720....:16730....:16740
GIFT_amh_LG23           15211 GAAACTTGTTCTTTCAAGCAGTGTGGGGAGTTCCTGAGTTAAGACCTTTGGCAGGATTTC
GIFT_amhΔy_h1tg000178l  14330 GAAACTTGTTCTTTCAAGCAGTGTGGGGAGTTCCTGAGTTAAGACCTTTGGCAGGATTTC
GIFT_amhy_h1tg000178l   14905 AGGACTGGAGATACTGGAAAGCGAAACAAACAAGTGGTTTATGTCACACCACCACATTAC
GIFT_amh_5'UTR             29 ------------------------------------------------------------

                              ....:16750....:16760....:16770....:16780....:16790....:16800
GIFT_amh_LG23           15271 GAGTAAACAGCCTTGACTGCAAA-----------------GCTCTGAAAGATTATCTACG
GIFT_amhΔy_h1tg000178l  14390 GAGTAAACAGCCTTGACTGCAAA-----------------GCTCTGAAAGATTATCTACG
GIFT_amhy_h1tg000178l   14965 TGATGCCTCACTAAAACTGTTAACAATTTTATGTCAGCTGTCAGTCAAATATTGACCACC
GIFT_amh_5'UTR             29 ------------------------------------------------------------

                              ....:16810....:16820....:16830....:16840....:16850....:16860
GIFT_amh_LG23           15314 AATAAT--GTACCAGGATGCAGTATGATGCAGAACTGCACA-------------------
GIFT_amhΔy_h1tg000178l  14433 AATAAT--GTACCAGGATGCAGTATGATGCAGAACTGCACA-------------------
GIFT_amhy_h1tg000178l   15025 AACTATGAGAAACTGTTTGTTTTTTGTCTCACAAAGACACATCGAGGCTACGGCTTCTGC
GIFT_amh_5'UTR             29 ------------------------------------------------------------

                              ....:16870....:16880....:16890....:16900....:16910....:16920
GIFT_amh_LG23           15353 -----------------GCTTTAAAGCCAAATCACA----CCCCAGAAATACTTTTAAAA
GIFT_amhΔy_h1tg000178l  14472 -----------------GCTTTAAAGCCAAATCACA----CCCCAGAAATACTTTTAAAA
GIFT_amhy_h1tg000178l   15085 ACACTTCTGCTGCGGGCACTTTAAGCTTGTTCCACAGTTCTCACAGGAAACCTGCCAGA-
GIFT_amh_5'UTR             29 ------------------------------------------------------------

                              ....:16930....:16940....:16950....:16960....:16970....:16980
GIFT_amh_LG23           15392 ACTATAAAACTGCCTAAAATCTGTATGTAATGAATGGATACCTCCCACCTTCC------
GIFT_amhΔy_h1tg000178l  14511 ACTATAAAACTGCCTAAAATCTGTATGTAATGAATGGAATACCTCCCACCTTCC------
GIFT_amhy_h1tg000178l   15144 -----AGAACAGGCTATTTTTTTCCTCTAGCCGCTAGCCTACTTTAGCTCTTCACTCACC
GIFT_amh_5'UTR             29 ------------------------------------------------------------

                              ....:16990....:17000....:17010....:17020....:17030....:17040
GIFT_amh_LG23           15446 ---TCAAAGTGCTCTGACAGGCACGTACACCC------------CCGCCTCCGCCCGCCA
GIFT_amhΔy_h1tg000178l  14565 ---TCAAAGTGCTCTGACAGGCACGTACACCC------------CCGCCTCCGCCCGCCG
GIFT_amhy_h1tg000178l   15199 GCGTCACTGCAGTTTAAGAGACTGGGACAAACAATACTGATCAGCCGTTCTAATTCAGCG
GIFT_amh_5'UTR             29 ------------------------------------------------------------

                              ....:17050....:17060....:17070....:17080....:17090....:17100
GIFT_amh_LG23           15491 GTGTCCCTCCAGATTAAAGATTCAAAGGTTAGACTGGGCTGTGAGAAAAGCCACCAGGTG
GIFT_amhΔy_h1tg000178l  14610 GTGTCCCTCCAGATTAAAGATTCAAAGGTTAGACTGGGCTGTGAGAAAAGCCACCAGGTG
GIFT_amhy_h1tg000178l   15259 CTGTCACAGCACTCCTCAGATTTAG---------CTTGCTGTGACTGCATACATTTA--C
GIFT_amh_5'UTR             29 ------------------------------------------------------------

                              ....:17110....:17120....:17130....:17140....:17150....:17160
GIFT_amh_LG23           15551 AGCTTGTGTGTAAGATTCTTAAACCATAAAACCCTGCAGCACAACAGGAGGAGAGTAGTA
GIFT_amhΔy_h1tg000178l  14670 AGCTTGTGTGTAAGATTCTTAAACCATAAAACCCTGCAGCACAACAGGAGGAGAGTAGTA
GIFT_amhy_h1tg000178l   15308 AGCTTTGAGCGAAGGCCTGTTCACTGGGATTCTATTAGGAACAAAAATAACAGCGCCGCA
GIFT_amh_5'UTR             29 ------------------------------------------------------------

                              ....:17170....:17180....:17190....:17200....:17210....:17220
GIFT_amh_LG23           15611 TTCATCAATAGCTGCGATTTCCTTGTTTTAAGTGAAGGAGAGCACCGTGCTGTCTGACTA
GIFT_amhΔy_h1tg000178l  14730 TTCATCAATAGCTGCGATTTCCTTGTTTTAAGTGAAGGAGAGCACCGTGCTGTCTGACTA
GIFT_amhy_h1tg000178l   15368 AACACCATCACACACAGGGATGCTCTCTTACGCAAATGCCACCAATGCGCTACATGTTGA
GIFT_amh_5'UTR             29 ------------------------------------------------------------

                              ....:17230....:17240....:17250....:17260....:17270....:17280
GIFT_amh_LG23           15671 GTAGTA---------------------------------------TTAAAAACTGACAG
GIFT_amhΔy_h1tg000178l  14790 GTAGTA---------------------------------------TTAAAAACTGACAG
GIFT_amhy_h1tg000178l   15428 GCAGAGGTGTGGAAACCAGTGTCATTCGCAGTGTGTAATATCCAACTTATGATCTCACAG
GIFT_amh_5'UTR             29 ------------------------------------------------------------

                              ....:17290....:17300....:17310....:17320....:17330....:17340
GIFT_amh_LG23           15691 AAATCTGAAGACACAATCAAAAGCATAACATGTAACTGCAATC-GTATCAAACACCAAGA
GIFT_amhΔy_h1tg000178l  14810 AAATCTGAAGACACAATCAAAAGCATAACATGTAACTGCAATC-GTATCAAACACCAAGA
GIFT_amhy_h1tg000178l   15488 AGGATTGGGGATTTTAACCAA--------ATGCAGTTACTCCCTAAATCCGGTAATCAGA
GIFT_amh_5'UTR             29 ------------------------------------------------------------

                              ....:17350....:17360....:17370....:17380....:17390....:17400
GIFT_amh_LG23           15750 ATAAGAAAATCAAAACACCATCGACAATAATTACAAAATACCT------CTCAGTGCTGG
GIFT_amhΔy_h1tg000178l  14869 ATAAGAAAATCAAAACGCCATCGACAATAATTACAAAATACCT------CTCAGTGCTGG
```

```
GIFT_amhy_h1tg000178l   15540 TTAGGAA----AGGAAGCCACTTGCCATCCTTGCAGAACCCCTTAACGCCTCCTTTCTGC
GIFT_amh_5'UTR             29 ------------------------------------------------------------

                              ....:17410....:17420....:17430....:17440....:17450....:17460
GIFT_amh_LG23           15804 TAACAAATGGACTAGTCTAATTCTAAAATGACTTTTTCTCTTCACTTGC----------A
GIFT_amhΔy_h1tg000178l  14923 TAACAAATGGACTAGTCTAATTCTAAAATGACTTTTTCTCTTCACTTGC----------A
GIFT_amhy_h1tg000178l   15596 GATTTAAAGAGAACACAAAGTTCTCAGTCAGTTTCTGCCTTTAGCATTCTTTGAGACTGA
GIFT_amh_5'UTR             29 ------------------------------------------------------------

                              ....:17470....:17480....:17490....:17500....:17510....:17520
GIFT_amh_LG23           15854 CACGAGTGCACGGGACAGGAGTTTTATTTTTCTATAAAGCTTTGATTATTACACTAATAT
GIFT_amhΔy_h1tg000178l  14973 CACGAGTGCACGGGACAGGAGTTTTATTTTTCTATAAAGCTTTGATTATTACACTAATAT
GIFT_amhy_h1tg000178l   15656 CATGTGAACAGTAGGCAG------TGCTGCGCAATGACGCCCTGACGGTCATCTTAGTA-
GIFT_amh_5'UTR             29 ------------------------------------------------------------

                              ....:17530....:17540....:17550....:17560....:17570....:17580
GIFT_amh_LG23           15914 TCCCTTAGATATGCGCTGGCAAGTGAAAAATTAAAAAA--ACCAAACAAAAACACACACG
GIFT_amhΔy_h1tg000178l  15033 TCCCTTAGATATGCGCTGGCAAGTGAAAAATTAAAAAAAAACCAAACAAAAACACACACG
GIFT_amhy_h1tg000178l   15709 -----AAAACAGCTGCTGTGAATAGTGTTTGTAGGAGACTCTGCTGTAAATGAACATCAT
GIFT_amh_5'UTR             29 ------------------------------------------------------------

                              ....:17590....:17600....:17610....:17620....:17630....:17640
GIFT_amh_LG23           15972 TACAAACTGAGCTTTAGTGTAATCCCACTATCAGAGGAAAAAAGGAAGCCTGTCTGATCT
GIFT_amhΔy_h1tg000178l  15093 TACAAACTGAGCTTTAGTGTAATCCCACTATCAGAGGAAAAAAGGAAGCCTGTCTGATCT
GIFT_amhy_h1tg000178l   15764 TGTAAAGGACATCTGCAATAAATCTCATGCATCCATCTAGCAGAGAATCATCTTTAATCC
GIFT_amh_5'UTR             29 ------------------------------------------------------------

                              ....:17650....:17660....:17670....:17680....:17690....:17700
GIFT_amh_LG23           16032 CACTCACACACACACACACACACACACACACACACACACACACACACACACACTAAGACAAG
GIFT_amhΔy_h1tg000178l  15153 CACTCACACACACACACACACACACAC-------------------------TAAGACAAG
GIFT_amhy_h1tg000178l   15824 CATGTTCTAGCCTGCACTGTTAGACACTTCAATGTAGTATGACAT-----TTGCAATGTG
GIFT_amh_5'UTR             29 ------------------------------------------------------------

                              ....:17710....:17720....:17730....:17740....:17750....:17760
GIFT_amh_LG23           16092 TGTTTCACATCAGCTCACGCGGCGAGGTACGGCCTCGTCAGTAAGACGTCTACAGGCGGT
GIFT_amhΔy_h1tg000178l  15189 TGTTTCACATCAGCTCACGCGGCGAGGTACGGCCTCGTCAGTAAGACGTCTACAGGCGGT
GIFT_amhy_h1tg000178l   15879 GGTTTTATTTTAGCTCACCTAACACGTTCTGGCTT--TCAACAAAAGAAAAATTAGCAGG
GIFT_amh_5'UTR             29 ------------------------------------------------------------

                              ....:17770....:17780....:17790....:17800....:17810....:17820
GIFT_amh_LG23           16152 GTGTGTAAAGAGGCTGACGCGCAGTTCTGTATGCACCGGTGAAGATTTTGATTGTAAGGT
GIFT_amhΔy_h1tg000178l  15249 GTGTGTAAAGAGGCTGACGCGCAGTTCTGTATGCACCGGTGAAGATTTTGATTGTAAGGT
GIFT_amhy_h1tg000178l   15937 TTAAAAA------------------------------TGTAACAGTCTTTAATGTGGGGT
GIFT_amh_5'UTR             29 ------------------------------------------------------------

                              ....:17830....:17840....:17850....:17860....:17870....:17880
GIFT_amh_LG23           16212 CTGAACTGCCCTCGCTTGGAAACAGAAGCACCATTTTTCCTGCAGAATCTGGGTTGGTAA
GIFT_amhΔy_h1tg000178l  15309 CTGAACTGCCCTCGCTTGGAAACAGAAGCACCATTTTTCCTGCAGAATCTGGGTTGGTAA
GIFT_amhy_h1tg000178l   15967 TAAATTTACACAAGTTTAAATAC-----------------------ATTTGATTTTATCA
GIFT_amh_5'UTR             29 ------------------------------------------------------------

                              ....:17890....:17900....:17910....:17920....:17930....:17940
GIFT_amh_LG23           16272 AAAAATTAAACACCAAAAACAAACAAAAAAACCAACAATAAATCAAAGAAGTGAAAATAC
GIFT_amhΔy_h1tg000178l  15369 AAAAATTAAACACCAAAAACAAACAAAAAAAACAACAATAAATCAAAGAAGTGAAAATAC
GIFT_amhy_h1tg000178l   16004 GGTGGTCAAAAATGAAAAGAATGAAAGAAAAGGCCTAGCCTTCTCTTAAGGTAACA---
GIFT_amh_5'UTR             29 ------------------------------------------------------------

                              ....:17950....:17960....:17970....:17980....:17990....:18000
GIFT_amh_LG23           16332 CTGAAACAAGTGATAACAGTTCTAGTACTGGAATCTGGTGGCGTTTGAGCCGCCGCACCT
GIFT_amhΔy_h1tg000178l  15429 CTGAAACAAGTGATAACAGTTCTAGTACTGGAATCTGGTGGCGTTTGAGCCGCCGCACCT
GIFT_amhy_h1tg000178l   16061 --------------GAGACTTTTTACCTTGGGAAGGGGGAGGCTATTAGCTGCTATA---
GIFT_amh_5'UTR             29 ------------------------------------------------------------

                              ....:18010....:18020....:18030....:18040....:18050....:18060
GIFT_amh_LG23           16392 TTGTGTAATAAACACACAGCCTCTGAGTTTGTGGCTGCGGGGGAT---------------
GIFT_amhΔy_h1tg000178l  15489 TTGTGTAATAAACACACAGCCTCTGAGTTTGTGGCTGCGGGGGATGTGTGTGTGTGTGTG
GIFT_amhy_h1tg000178l   16104 -------------------------------------------ATAGATGATGTTATTTT
GIFT_amh_5'UTR             29 ------------------------------------------------------------

                              ....:18070....:18080....:18090....:18100....:18110....:18120
GIFT_amh_LG23           16437 -GTGTGTGTGTGTGTGTTTTCAGGTGCTGATGTGCAGCCATGTTAAGGTCCTCTGGTGGC
GIFT_amhΔy_h1tg000178l  15549 TGTGTGTGTGTGTGTGTTTTCAGGTGCTGATGTGCAGCCATGTTAAGGTCCTCTGGTGGC
GIFT_amhy_h1tg000178l   16121 AATGCATGCTTTATTATTGGAAGGTGTGGATGTTAAACCAGCCTTATCACACATGATAGC
```

```
GIFT_amh_5'UTR                 29 ------------------------------------------------------------

                                  ....:18130....:18140....:18150....:18160....:18170....:18180
GIFT_amh_LG23               16496 AATGCTTTGCTCTGCTCAGCTTCAAGCCTCCTGTTACAGGGTGAATTTAAGGGTCACAAT
GIFT_amhΔy_h1tg000178l      15609 AATGCTTTGCTCTGCTCAGCTTCAAGCCTCCTGTTACAGGGTGAATTTAAGGGTCACAAT
GIFT_amhy_h1tg000178l       16181 TATGTGTCAGCATATTTA---TAAAGGCATCTTTTCAAGCTTTGGCTTAAAGA-------
GIFT_amh_5'UTR                 29 ------------------------------------------------------------

                                  ....:18190....:18200....:18210....:18220....:18230....:18240
GIFT_amh_LG23               16556 GCCAGCACCAAAACACCCCTTTCACATATATATGTATATATATTTATATACTTATATATA
GIFT_amhΔy_h1tg000178l      15669 GCCAGCACCAAAACACCCCTTTCACATATATATGTATATATATTTATATACTTATATATA
GIFT_amhy_h1tg000178l       16231 ---------GAAACACTATTATATAATGTACATATATAAATATCTAACCAT-----GACA
GIFT_amh_5'UTR                 29 ------------------------------------------------------------

                                  ....:18250....:18260....:18270....:18280....:18290....:18300
GIFT_amh_LG23               16616 TTTATATAGACTGTATATAGTTACTTATTTTATATACCTTTCTGTTTATGATGGAGATGT
GIFT_amhΔy_h1tg000178l      15729 TTTATATAGACTGTATATAGTTACTTATTTTATATACCTTTCTGTTTATGATGGAGATGT
GIFT_amhy_h1tg000178l       16277 ATTACATAACAATTACATAGCCGAAACAGCTGAACACTCTTGCAC--------------T
GIFT_amh_5'UTR                 29 ------------------------------------------------------------

                                  ....:18310....:18320....:18330....:18340....:18350....:18360
GIFT_amh_LG23               16676 ACAATTAAAGAAAACTTATTTACAAAAACAGTGTGTCTTGTTCACCAACAGCGGGCCGAT
GIFT_amhΔy_h1tg000178l      15789 ACAATTAAAGAAAACTTATGTACAAAAACAGTGTGTCTTGTTCACCAACAGCGGGCCGAT
GIFT_amhy_h1tg000178l       16323 GCCACTGCAAAGAGATTAGGGGCTGCAGAGCGGCGT-------------AACAAGTTAGT
GIFT_amh_5'UTR                 29 ------------------------------------------------------------

                                  ....:18370....:18380....:18390....:18400....:18410....:18420
GIFT_amh_LG23               16736 GTATGTTTAAAACAATAACAGGGGCCCCCGTGGCCGTCGCTCGGGCCCCGAGCCGGCTGC
GIFT_amhΔy_h1tg000178l      15849 GTATGTTTAAAACATTAACAGGGGCCCCCGTGGCCGTCGCTCGGGCCCCGAGCCGGCTGC
GIFT_amhy_h1tg000178l       16370 ATTAGTTAACAGTAATAACCTTGGC-----------------------------------
GIFT_amh_5'UTR                 29 ------------------------------------------------------------

                                  ....:18430....:18440....:18450....:18460....:18470....:18480
GIFT_amh_LG23               16796 ACGAGAGCACTTTCATCCAAAGGTGGCGTCTACAGTTGTACACGATGCATTCAAAGAACG
GIFT_amhΔy_h1tg000178l      15909 ACGAGAGCACTTTCATCCAAAGGTGGCGTCTACAGTTGTACACGATGCATTCAAAGAACG
GIFT_amhy_h1tg000178l       16395 -------------TATCACAGGTTAGTGTATACTTCTTTGGGTGATGGATGCCGAACTTT
GIFT_amh_5'UTR                 29 ------------------------------------------------------------

                                  ....:18490....:18500....:18510....:18520....:18530....:18540
GIFT_amh_LG23               16856 AGACAATAGCATATTAAATATGAGAAAGGAAATGCCAAAAAACGGCTTTACAAATGTCCC
GIFT_amhΔy_h1tg000178l      15969 AGACAATAGCATATTAAATATGAGAAAGGAAATGCCAAAAAACGGCTTTACAAATGTCCC
GIFT_amhy_h1tg000178l       16442 TCACAGCTATCTTTGA--------------------GAGTAACAGAGTCATTATCGTATT
GIFT_amh_5'UTR                 29 ------------------------------------------------------------

                                  ....:18550....:18560....:18570....:18580....:18590....:18600
GIFT_amh_LG23               16916 TGTGTACAGCACCAAGTACTTGACTTGTTTTTGATGTTTTCATTACAGAATAAATAAGA-
GIFT_amhΔy_h1tg000178l      16029 TGTGTACAGCACCAAGTACTTGACTTGTTTTTGATGTTTTCATTACAGAATAAATAAGA-
GIFT_amhy_h1tg000178l       16482 TTTGG--------TACTGCAAGATGTGTGCTTAATAATTTTATAATATAGCAATAATAAT
GIFT_amh_5'UTR                 29 ------------------------------------------------------------

                                  ....:18610....:18620....:18630....:18640....:18650....:18660
GIFT_amh_LG23               16975 --TCAGGCATAGTCCAAACTTAGCACCATTGTTTTCAGACTGT------GAGTCAAGTAC
GIFT_amhΔy_h1tg000178l      16088 --TCAGGCATAGTCCAAACTTAGCACCATTGTTTTCAGACTGT------GAGTCAAGTAC
GIFT_amhy_h1tg000178l       16534 TTTCATAAATGTTCCAAAGAGCGCAGAAATGTCTACAGCCGGTTTCTCCAAGTCTAGGAA
GIFT_amh_5'UTR                 29 ------------------------------------------------------------

                                  ....:18670....:18680....:18690....:18700....:18710....:18720
GIFT_amh_LG23               17027 ACACACACTAATGCATTCTACAATGTCAACACAATTCAGGATTTAAAAAAAAAAAAAAAA
GIFT_amhΔy_h1tg000178l      16140 ACACACACTAATGCATTCTACAATGTCAACACAATTCAGGATTTAAAAAAAAAAAAAAAA
GIFT_amhy_h1tg000178l       16594 ATGCACACTAAACTGCTCAAGCTGGACAAACAGCTCTACAGCTCAGAGGACCAAAAATAA
GIFT_amh_5'UTR                 29 ------------------------------------------------------------

                                  ....:18730....:18740....:18750....:18760....:18770....:18780
GIFT_amh_LG23               17087 AAA---------------------------------------------------------
GIFT_amhΔy_h1tg000178l      16200 AAAAAAAAAAGGCAAATGGAAAGACCGACAGCATAAATATAATCATCTAAATTATCCACA
GIFT_amhy_h1tg000178l       16654 ACTGGACTTTG-----------GATCGTCTCTTTAAATATCACATGCTCAAATATT----
GIFT_amh_5'UTR                 29 ------------------------------------------------------------

                                  ....:18790....:18800....:18810....:18820....:18830....:18840
GIFT_amh_LG23               17090 ------------------------------------------------------------
GIFT_amhΔy_h1tg000178l      16260 ACTATACAGTCAGAGAATGAATCACATTCACAGTACAAACATTGTTTACAAAAACAAATT
GIFT_amhy_h1tg000178l       16699 --------------------------------------------------ATGATATATG
GIFT_amh_5'UTR                 29 ------------------------------------------------------------
```

```
                               ....:18850....:18860....:18870....:18880....:18890....:18900
GIFT_amh_LG23            17090 ------------------------------------------------------------
GIFT_amhΔy_h1tg000178l   16320 ACACCGTTTTTTAAAAAGCAAACAAAGAGGTGTGAAACAAGTCCAGTAGTTTGTTGTGGT
GIFT_amhy_h1tg000178l    16709 AGAATTTTTATTATGCAACACTATAGGATTTCCACAGTATTTAGGGCTGGTTGATATGGA
GIFT_amh_5'UTR              29 ------------------------------------------------------------

                               ....:18910....:18920....:18930....:18940....:18950....:18960
GIFT_amh_LG23            17090 ------------------------------------------------------------
GIFT_amhΔy_h1tg000178l   16380 TATGGTCAACTATGTTTTCCCAGTGTGTGAGGATTTGCATGCATGACTGTGTGTGTGTAT
GIFT_amhy_h1tg000178l    16769 ATCACACGTGTCTTGTGTCCCATTGTCTG-------------------------------
GIFT_amh_5'UTR              29 ------------------------------------------------------------

                               ....:18970....:18980....:18990....:19000....:19010....:19020
GIFT_amh_LG23            17090 ------------------------------------------------------------
GIFT_amhΔy_h1tg000178l   16440 GTGTGTTTAAACTAAACGCAGTAAATTTTCACATATCAGTCCACTTTCATTTAAACCTGC
GIFT_amhy_h1tg000178l    16798 -----------------------------------CCTCCAGGTTTCCATATGGGACGGC
GIFT_amh_5'UTR              29 ------------------------------------------------------------

                               ....:19030....:19040....:19050....:19060....:19070....:19080
GIFT_amh_LG23            17090 ------------------------------------------------------------
GIFT_amhΔy_h1tg000178l   16500 ATTTCCTAACAGCACGGATAAATATGAATGCATGGATCTCTGCTGTTATGCTTAGCTTAC
GIFT_amhy_h1tg000178l    16823 TTCACCCGGTGGTGGGGATCTGAAGAAGT-------------------CGGAGCGATAAG
GIFT_amh_5'UTR              29 ------------------------------------------------------------

                               ....:19090....:19100....:19110....:19120....:19130....:19140
GIFT_amh_LG23            17090 ------------------------------------------------------------
GIFT_amhΔy_h1tg000178l   16560 AAAGAGGAATATTTCTGCTGCCGTTTGTGTGACTTCACAAACCGACGGGAAAGAAACGGT
GIFT_amhy_h1tg000178l    16864 AAGGAGGAAGAAGCCAGCTGCTAATAAAGCAATCCTGAAACCCAAAATCAAAGCATAAAA
GIFT_amh_5'UTR              29 ------------------------------------------------------------

                               ....:19150....:19160....:19170....:19180....:19190....:19200
GIFT_amh_LG23            17090 ------------------------------------------------------------
GIFT_amhΔy_h1tg000178l   16620 TAATAAATACACGCAAACTCACTGAACTCTGTTGTTCTTGGACATGTGGATTCAGCTGTT
GIFT_amhy_h1tg000178l    16924 CATATACTGTAGCTTATGTAGCTGAATGGCTACGAACACAGAACTTTAAGTACACTTTGC
GIFT_amh_5'UTR              29 ------------------------------------------------------------

                               ....:19210....:19220....:19230....:19240....:19250....:19260
GIFT_amh_LG23            17090 ------------------------------------------------------------
GIFT_amhΔy_h1tg000178l   16680 CGCTGAACCTTAACAGAGCAGCCACAGTGACGGGTGGCTGAAGAGCACCATTTCATTTTT
GIFT_amhy_h1tg000178l    16984 ATTTAATTATTACAAGTTTG-------------------AGAAGGCTATATTTCATCCTT
GIFT_amh_5'UTR              29 ------------------------------------------------------------

                               ....:19270....:19280....:19290....:19300....:19310....:19320
GIFT_amh_LG23            17090 ------------------------------------------------------------
GIFT_amhΔy_h1tg000178l   16740 TTTAAAACAGATGTTAATGTACAAGACGCTGCATTGCATTATTGGCACCCACAGGCATGT
GIFT_amhy_h1tg000178l    17025 TTAA---------------------------CAGTTTAACTTTAACACTCAGAAACATTA
GIFT_amh_5'UTR              29 ------------------------------------------------------------

                               ....:19330....:19340....:19350..
GIFT_amh_LG23            17090 --------------------------------
GIFT_amhΔy_h1tg000178l   16800 TATTAATTTAACTTATCATCAAAATAAAGC--
GIFT_amhy_h1tg000178l    17058 TTCCAATTAGAAATATTTTCAAAATGACAAAA
GIFT_amh_5'UTR              29 --------------------------------
```
