## supplementary figure 3 for "High-quality genome assembly for the genetically improved Abbassa Nile tilapia enables the reconstruction of X and Y haplotypes"

**Supplementary Figure 3** – Alignment of Abbassa *amh, amhy,* and *amhΔy* 5' UTR and gene promoter region. Nucleotide sequence conservation is shaded and 346 bp Abbassa *amh* 5' UTR sequence shown for reference. Green nucleotides on *amhΔy* sequence indicate the 46 bp 5' UTR. The 165 bp deletion in *amhΔy* gene promoter and 3 bp 'TCT' insertion in *amhy* gene promoter marked in red. The deletions at > 1.6 kb in *amhy* and *amhΔy* are marked in green.

```
                                   ....:10810....:10820....:10830....:10840....:10850....:10860
Abbassa_amh_h2tg000119l       9936 AATAACCACATGCATGCATTTTTCCATTCAAAGTTAACGAGTTATTAATTAATCAGCTGA
Abbassa_amhΔy_h1tg000212l     9676 AATAACCACATGCATGCATTTTTCCATTCAAAGTTAACGAGTTATTAATTAATCAGCTGA
Abbassa_amh_5’UTR                1 ------------------------------------------------------------
Abbassa_amhy_h1tg000212l      9936 AATAACCACATGCATGCATTTTTCCATTCAAAGTTAACGAGTTATTAATTAATCAGCTGA

                                   ....:10870....:10880....:10890....:10900....:10910....:10920
Abbassa_amh_h2tg000119l       9996 GCGGCGTCTGACCGGTTACTGTGGGGTCCTGGGCCGGTTGCAGGGTCCAGCAGAGTGTCA
Abbassa_amhΔy_h1tg000212l     9736 GCGGCGTCTGACCGGTTACTGTGGGGTCCTGGGCCGGTTGCAGGGTCCAGCAGAGTGTCA
Abbassa_amh_5'UTR                1 ------------------------------------------------------------
Abbassa_amhy_h1tg000212l      9996 GCGGCGTCTGACCGGTTACTGTGGGGTCCTGGGCCGGTTGCAGGGTCCAGCAGAGTGTCA

                                   ....:10930....:10940....:10950....:10960....:10970....:10980
Abbassa_amh_h2tg000119l      10056 GCGCCTCGCTGTAAAGAACGAGCAGACCCAACATGTTTGCAGTGTCTGCGTGTGCGTTTG
Abbassa_amhΔy_h1tg000212l     9796 GCGCCTCGCTGTAAAGAACGAGCAGACCCAACATGTTTGCAGTGTCTGCGCGTGCGTTTG
Abbassa_amh_5'UTR                1 ----------------------------------GTTTGCAGTGTCTGCGCGTGCGTCTG
Abbassa_amhy_h1tg000212l     10056 GCGCCTCGCTGTAAAGAACGAGCAGACCCAACATGTTTGCAGTGTCTGCGCGTGCGTCTG

                                   ....:10990....:11000....:11010....:11020....:11030....:11040
Abbassa_amh_h2tg000119l      10116 TGTGGTCAGATCTCTCACAGGATGGGAGTTACATCCTCAGACCTCCCCTTCTCACCACCT
Abbassa_amhΔy_h1tg000212l     9856 TGTGGTCAGATCTCTCACAGGATGGGAGTTACATCCTCAGACCTCCCCTTCTCACCACCC
Abbassa_amh_5'UTR               26 TGTGGTCAGATCTCTCACAGGATGGGAGTTACATCCTCAGACCTCCCCTTCTCACCACCT
Abbassa_amhy_h1tg000212l     10116 TGTGGTCAGATCTCTCACAGGATGGGAGTTACATCCTCAGACCTCCCCTTCTCACCACCC

                                   ....:11050....:11060....:11070....:11080....:11090....:11100
Abbassa_amh_h2tg000119l      10176 GGGGTCCCCTTTCTGCCAAAATAGCAGCACTGTGTGTCCTTGATGACTGAGATTGTCAGT
Abbassa_amhΔy_h1tg000212l     9916 GGGGTCCCCTTTCTGCCAAAATAGCAGCACTGTGTGTCCTTGATGACTGAGATTGTCAGT
Abbassa_amh_5’UTR               86 GGGGTCCCCTTTCTGCCAAAATAGCAGCACTGTGTGTCCTTGATGACTGAGATTGTCAGT
Abbassa_amhy_h1tg000212l     10176 GGGGTCCTCTTTCTGCCAAAATAGCAGCACTGTGTGTCCTTGATGACTGAGATTGTCAGT

                                   ....:11110....:11120....:11130....:11140....:11150....:11160
Abbassa_amh_h2tg000119l      10236 ATTTGAGGTATTTTAACCTGCTTGTGGAGAACATTCTAAAATCAGACAGCAAACGGGACA
Abbassa_amhΔy_h1tg000212l     9976 ATTTGAGGTATTTTAACCTGCTTGTGGAGAACATTCTAAAATCAGACAGCAAACGCGACA
Abbassa_amh_5’UTR              146 ATTTGAGGTATTTTAACCTGCTTGTGGAGAACATTCTAAAATCAGACAGCAAACGGGACA
Abbassa_amhy_h1tg000212l     10236 ATTTGAGGTATTTTAACCTGCTTGTGGAGAACATTCTAAAATCAGACAGCAAACGCGACA

                                   ....:11170....:11180....:11190....:11200....:11210....:11220
Abbassa_amh_h2tg000119l      10296 CGGAGGTAAACAGAAGACGACTTTGGACACACTGAACATCCTTATCTAGCAGACACAAAC
Abbassa_amhΔy_h1tg000212l    10036 CGGAGGTAAACAGAAGACGACTTTGGACACACTGAACATCCTTATCTAGCAGACACAAAC
Abbassa_amh_5’UTR              206 CGGAGGTAAACAGAAGACGACTTTGGACACACTGAACATCCTTATCTAGCAGACACAAAC
Abbassa_amhy_h1tg000212l     10296 CGGAGGTAAACAGAAGACGACTTTGGACACACTGAACATCCTTATCTAGCAGACACAAAC

                                   ....:11230....:11240....:11250....:11260....:11270....:11280
Abbassa_amh_h2tg000119l      10356 AGGTCCCGGAAAGAAAGTTTCTCACGAACCTTTCATAGAATACACCAGGCTGAAAGATGA
Abbassa_amhΔy_h1tg000212l    10096 AGGTCCCAGAAAGAAAGTTTCTCACGAACCTTTCATAGAATACATCAGGCTGAAAGATGA
Abbassa_amh_5’UTR              266 AGGTCCCGGAAAGAAAGTTTCTCACGAACCTTTCATAGAATACACCAGGCTGAAAGATGA
Abbassa_amhy_h1tg000212l     10356 AGGTCTCAGAAAGAAAGTTTCTCACAAACCTTTCATAGAATACACCAGGCTGAAAGATGA

                                   ....:11290....:11300....:11310....:11320....:11330....:11340
Abbassa_amh_h2tg000119l      10416 AGAGTGCTGTTTAATGTTTCGTGGCTGCAGAGCTAATATGCACGCTGCTTAACTTAACCC
Abbassa_amhΔy_h1tg000212l    10156 AGAGTGCCGTTTAAAGTTTCGTGGCTGCAGAGCTAATATGCACGCTGCTTAACTTAACCC
Abbassa_amh_5’UTR              326 AGAGTGCTGTTTAATGTTTC----------------------------------------
Abbassa_amhy_h1tg000212l     10416 AGAGTGCTGTTTAATGTTTCGTGGCTGCAGAGCTAATATGCACGCTGCTTAACTTAACCC

                                   ....:11350....:11360....:11370....:11380....:11390....:11400
Abbassa_amh_h2tg000119l      10476 TCTCGAGGCAGGCGTTGCCGATTTGCAACAGTTAAAAACTAACAACCTGATTACCCTACA
Abbassa_amhΔy_h1tg000212l    10216 TCTCGAGGCAGGCGTTGCCGATTTGCAACAGTTAAAAACTAACAACCTGATTACCCTACA
Abbassa_amh_5’UTR               29 ------------------------------------------------------------
Abbassa_amhy_h1tg000212l     10476 TCTCGAGGCAGGCGTTGCCGATTTGCAACAGTTAAAAACTAACAACCTGATTACCCTACA

                                   ....:11410....:11420....:11430....:11440....:11450....:11460
Abbassa_amh_h2tg000119l      10536 TACATATTTTATGAGTCTTTTTTTTTTTTTTGATAATTTCCCCAAATATCAGATTTTTCC
Abbassa_amhΔy_h1tg000212l    10276 TACATATTTTATGAGTCTTTTTTTTTTTTTTGATAATTTCCCCAAATATCAGATTTTTCC
Abbassa_amh_5’UTR               29 ------------------------------------------------------------
Abbassa_amhy_h1tg000212l     10536 TACATATTTTATGAGTCTTTTTTTTTTTTTTGATAATTTCCCCAAATATCAGATTTTTCC

                                   ....:11470....:11480....:11490....:11500....:11510....:11520
Abbassa_amh_h2tg000119l      10596 TGTAACTAAAACTTAATTTGAGCCTGAGAGGGTTAAACAAAGAGGACAGTAAAACTGGAG
Abbassa_amhΔy_h1tg000212l    10336 TGTAACTAAAACTTAATTTGAGCCTGAGAGGGTTAAACAAAGAGGACAGTAGAACTGGAG
Abbassa_amh_5’UTR               29 ------------------------------------------------------------
Abbassa_amhy_h1tg000212l     10596 TGTAACTAAAACTTAATTTGAGCCTGAGAGGGTTAAACAAAGAGGACAGTAGAACTGGAG

                                   ....:11530....:11540....:11550....:11560....:11570....:11580
```

```
Abbassa_amh_h2tg000119l     10656 GTTTGCTTCCATCACGTCGACAACAAAGCTCAAAACGATTTTAAAGAAACACTTTGCATT
Abbassa_amhΔy_h1tg000212l   10396 GTTTGCTTCCATCACGTCGACAACAAAGCTCAAAACGATTTTAAAGAAACACTTTGCATT
Abbassa_amh_5'UTR              29 ------------------------------------------------------------
Abbassa_amhy_h1tg000212l    10656 GTTTGCTTCCATCATGTCGACAACAAAGCTCAAAACGATTTTAAAGAAACACTTTGCATT

                                  ....:11590....:11600....:11610....:11620....:11630....:11640
Abbassa_amh_h2tg000119l     10716 TTGGTAAATCTGCTTACTTGTTGCTTTGGACTGATGAC---TCCTCTTAGAGAGACTGCT
Abbassa_amhΔy_h1tg000212l   10456 T-----------------------------------------------------------
Abbassa_amh_5'UTR              29 ------------------------------------------------------------
Abbassa_amhy_h1tg000212l    10716 TTGGTAAATCTGCTTACTTGTTGCTTTGGACTGATGACTCTTCCTCTTAGAGAGACTGCT

                                  ....:11650....:11660....:11670....:11680....:11690....:11700
Abbassa_amh_h2tg000119l     10773 GGCAGAAAGCCTTGAAGGGAAACTTCAGCCACATCCACTGTTTTTCATCTTTCTGTTTCT
Abbassa_amhΔy_h1tg000212l   10457 ------------------------------------------------------------
Abbassa_amh_5'UTR              29 ------------------------------------------------------------
Abbassa_amhy_h1tg000212l    10776 GGCAGAAAGCCTTGAAGGGAAACTTCAGCCACATCCACTGTTTTTCATCTTTCTGTTTCT

                                  ....:11710....:11720....:11730....:11740....:11750....:11760
Abbassa_amh_h2tg000119l     10833 CTAAGCGGGGGATGTCCAACATCAGGCCCAGGGGCTAGAATCACCCAGCAATGACTCCAA
Abbassa_amhΔy_h1tg000212l   10457 ---------------------------------------------CAGCAATGACTCCAA
Abbassa_amh_5'UTR              29 ------------------------------------------------------------
Abbassa_amhy_h1tg000212l    10836 CTAAGCGGGGGATGTCCAACATCAGGCCCAGGGGCTAGAATCACCCAGCAATGACTCCAA

                                  ....:11770....:11780....:11790....:11800....:11810....:11820
Abbassa_amh_h2tg000119l     10893 CCCAGCTCACTTGAAGGATGGCATAAATTTTGGA--------------CTTTTAACTGTA
Abbassa_amhΔy_h1tg000212l   10472 CCCAGCTCACTTGAAGGATGGCATAAATTTTGGA--------------CTTTTAACTGTA
Abbassa_amh_5'UTR              29 ------------------------------------------------------------
Abbassa_amhy_h1tg000212l    10896 CCCAGCTCACTTGAAGGATGGCAAAAATTTGCCAGCACCAAAACACCCCTTTCACATATA

                                  ....:11830....:11840....:11850....:11860....:11870....:11880
Abbassa_amh_h2tg000119l     10939 TTTTCTTA---AATTTTATGGCTTTTCCTGCTAATAAAGAAACTCTGCCACATGTTCATGC
Abbassa_amhΔy_h1tg000212l   10518 TTTTCTTA---AATTTTAAGGCTTTTCCTGCTAATAAAGAAACTCTGCCACATGTTCATGC
Abbassa_amh_5'UTR              29 ------------------------------------------------------------
Abbassa_amhy_h1tg000212l    10956 TATGTATATATATTTATATACTTATATATATTTATATAGACTGTATATAGTTACTTATTT

                                  ....:11890....:11900....:11910....:11920....:11930....:11940
Abbassa_amh_h2tg000119l     10996 TACACC---------------AAAGTGATTAACAATTA---------CATGACAAAAAAGT
Abbassa_amhΔy_h1tg000212l   10575 TACACC---------------AAAGTGATTAACAATTA---------CATGACAAAAAAGT
Abbassa_amh_5'UTR              29 ------------------------------------------------------------
Abbassa_amhy_h1tg000212l    11016 TACATACCTTTCTGTTTATGACGGAGATGTACAATTAAAGAAAACTTATGTACAAAAACA

                                  ....:11950....:11960....:11970....:11980....:11990....:12000
Abbassa_amh_h2tg000119l     11033 TCCTG-TTTTTTTCACCATCTGCTCCAGAAATTAGTTTTTTTCTGTAATATTAC------
Abbassa_amhΔy_h1tg000212l   10612 TCCTGTTTTTTTTCACCATCTGCTCCAGAAATTAGTTTTTTTCTGTAATATTAC------
Abbassa_amh_5'UTR              29 ------------------------------------------------------------
Abbassa_amhy_h1tg000212l    11076 GTGTG-TCTTGTTCACCAACAGC-----GGGCCGATGTATGTTTAAAACATTAACAGGGG

                                  ....:12010....:12020....:12030....:12040....:12050....:12060
Abbassa_amh_h2tg000119l     11086 --------------------ACATTTATTTATTTATAATGGAAAATTTCTTTGATTTCAA
Abbassa_amhΔy_h1tg000212l   10666 --------------------ACATTTATTTATTTATGATGGAAAATTTCTTTGATTTCAA
Abbassa_amh_5'UTR              29 ------------------------------------------------------------
Abbassa_amhy_h1tg000212l    11130 CCCCCGTGGCCGTCGCTCGGGCCCCGAGCCGGCTGCACGAGAGCA---CTTTCATCCAAA

                                  ....:12070....:12080....:12090....:12100....:12110....:12120
Abbassa_amh_h2tg000119l     11126 GATTCAAAGTGTTTATTGTC-------ATGTGTCCAAAGA-----AAAAAGGCATTTC--
Abbassa_amhΔy_h1tg000212l   10706 GATTCAAAGTGTTTATTGTC-------ATGTGTCCAAAGA-----AAAAAGGCATTTC--
Abbassa_amh_5'UTR              29 ------------------------------------------------------------
Abbassa_amhy_h1tg000212l    11187 GGT----GGCGTCTACAGTTGTACACGATGCATTCAAAGAACGAGACAATAGCATATTAA

                                  ....:12130....:12140....:12150....:12160....:12170....:12180
Abbassa_amh_h2tg000119l     11172 -TCTGTGCAATGAAACTT--------TTGCTTTGC-----------TGTCCACCCACAGAT
Abbassa_amhΔy_h1tg000212l   10752 -TCTGTGCAGTGAAACTT--------TTGCTTTGC-----------TGTCCACCCACAGAT
Abbassa_amh_5'UTR              29 ------------------------------------------------------------
Abbassa_amhy_h1tg000212l    11243 ATATGAGAAAGGAAATGCCAAAAAACGGCTTTACAAATGTCCCTGTGTACAGCACCAAGT

                                  ....:12190....:12200....:12210....:12220....:12230....:12240
Abbassa_amh_h2tg000119l     11213 GCCGG--------TTAATATTTACAGTAGAAATAGAACAGATAAATACAAATA-------
Abbassa_amhΔy_h1tg000212l   10793 GCCGG--------TTAATATTTACAATAGAAATAGAACAGATAAATACAAATA-------
Abbassa_amh_5'UTR              29 ------------------------------------------------------------
Abbassa_amhy_h1tg000212l    11303 ACTTGACTTGTTTTTGATGTTTTCATT----ACAGAATAATAAGATCAGGCATAGTCCA

                                  ....:12250....:12260....:12270....:12280....:12290....:12300
Abbassa_amh_h2tg000119l     11258 -------------------------------------GCACAAATAAATAAAGACAGAA
```

```
Abbassa_amhΔy_h1tg000212l  10838 --------------------------------------GCACAAATAAATAAAGACAGAA
Abbassa_amh_5'UTR             29 ------------------------------------------------------------
Abbassa_amhy_h1tg000212l   11359 AACTTAGCACCATTGTTTTCAGACTGTGAGTCAAGTACACACACACTAATGCATTCTACA

                                 ....:12310....:12320....:12330....:12340....:12350....:12360
Abbassa_amh_h2tg000119l    11280 AAGGAGACAAATTATTGAAGTGTGAAACATAGATGTGTGCA-----AAAGAGCTATAGC-
Abbassa_amhΔy_h1tg000212l  10860 AAGGAGACAAATTATTGAAGTGTGAAACAGAGATGTGTGCA-----AAAGAGCTATAGC-
Abbassa_amh_5'UTR             29 ------------------------------------------------------------
Abbassa_amhy_h1tg000212l   11419 ATGTCAACACAATTCAGGATTTAAAAAAAAAAAAAAAGGCAAATGGAAAGACCGACAGCA

                                 ....:12370....:12380....:12390....:12400....:12410....:12420
Abbassa_amh_h2tg000119l    11334 TTAATATGCTGGCTTAA--TATGCAGGATGACCTGATGTGCAAAAAATTATT---ATTAA
Abbassa_amhΔy_h1tg000212l  10914 TTAATATGCTGGCTTAA--TATGCAGGATGACCTGATGTGCAAAAAATTATT---ATTGA
Abbassa_amh_5'UTR             29 ------------------------------------------------------------
Abbassa_amhy_h1tg000212l   11479 TAAATATAATCATCTAAATTATCCACAACTATACAGT---CAGAGAATTAATCACATTCA

                                 ....:12430....:12440....:12450....:12460....:12470....:12480
Abbassa_amh_h2tg000119l    11389 TGTT--------------CAAAAATAGGTCAGCTTGTTTGAGTCTGATGGCA--------
Abbassa_amhΔy_h1tg000212l  10969 TGTT--------------CAAAAATAGGTCAGCTTGTTTGAGTCTGATGGCA--------
Abbassa_amh_5'UTR             29 ------------------------------------------------------------
Abbassa_amhy_h1tg000212l   11536 TAGTACAAACATCGTTTACAAAAACAAATTACACCGTTT--TTTAAAAAGCAAACAAAGA

                                 ....:12490....:12500....:12510....:12520....:12530....:12540
Abbassa_amh_h2tg000119l    11427 -GTGGGGAAGAAGGC--GTTGTTGAGTCTGGATGTTCTGGATTTCACACTTCTAAACCTC
Abbassa_amhΔy_h1tg000212l  11007 -GTGGGGAAGAAGGC--GTTGTTGAGTCTGGATGTTCTGGATTTCACACTTCTAAACCTC
Abbassa_amh_5'UTR             29 ------------------------------------------------------------
Abbassa_amhy_h1tg000212l   11594 GGTGTGAAACAAGTCCAGTAGTTTGTTGTGGTTATGGTCAA----------CTATGTTTT

                                 ....:12550....:12560....:12570....:12580....:12590....:12600
Abbassa_amh_h2tg000119l    11484 CGCCCCGAGGGCAGAAGTGTGAACAGTCCGTGTTGGGGATGTGTGGGGTCTTTGAGGATG
Abbassa_amhΔy_h1tg000212l  11064 CGCCCCGAGGGCAGAAGTGTGAACAGTCCATGTTGGGGATGTGTGGGGTCTTTGAGGATG
Abbassa_amh_5'UTR             29 ------------------------------------------------------------
Abbassa_amhy_h1tg000212l   11644 CCCAGTGTGTGAGGATTTGCATGCATGACTGTGTGTGTGTGTGTGTGTGTGTTTAAACTA

                                 ....:12610....:12620....:12630....:12640....:12650....:12660
Abbassa_amh_h2tg000119l    11544 GAGGCAG----------------------CTCTCCTTTGGACTCTGCGATGGTAGATGCT
Abbassa_amhΔy_h1tg000212l  11124 GAGGCGG----------------------CTCTCCTCTGGACTCTGCGATGGTAGATGCT
Abbassa_amh_5'UTR             29 ------------------------------------------------------------
Abbassa_amhy_h1tg000212l   11704 AACGCAGTAAAATTTCACATATCAGTCCACTTTCATTTAAAC-CTGCATTTCCTAACA--

                                 ....:12670....:12680....:12690....:12700....:12710....:12720
Abbassa_amh_h2tg000119l    11582 GTGCAGAGAGGGCAGCGGAGTCCTGATTATCTTCCCTGCAGTTGTTATCACTCTCTGCAG
Abbassa_amhΔy_h1tg000212l  11162 GTGCAGAGAGGGCAGCGGAGTCCTGATTATCTTCCCTGCAGTTGTTATCACTCTCTGCAG
Abbassa_amh_5'UTR             29 ------------------------------------------------------------
Abbassa_amhy_h1tg000212l   11761 --GCACGGATAAATATGAATGCATGG-----ATCTCTGCCGTTATGCTTAGCTTACAAAG

                                 ....:12730....:12740....:12750....:12760....:12770....:12780
Abbassa_amh_h2tg000119l    11642 GTGATTGCAGTCCATGGCAGTAGTGCTGCCGTATCATGCAGTGATGCAGCTGGTCAGTGT
Abbassa_amhΔy_h1tg000212l  11222 GTGATTGCGGTCCATGGCAGTAGTGCTGCCATATCATGCAGTGATGCAGCTGGTCAGTGT
Abbassa_amh_5'UTR             29 ------------------------------------------------------------
Abbassa_amhy_h1tg000212l   11814 AGGAAT------------ATTTCTGCTGCCGTTT----------------------GTGT

                                 ....:12790....:12800....:12810....:12820....:12830....:12840
Abbassa_amh_h2tg000119l    11702 GCTCTCCACAATGCAGCTGT--AGAACCTGCTGAGGATCATACCAAATTTCCTCAGCCTC
Abbassa_amhΔy_h1tg000212l  11282 GCTCTCCACAATGCAGCTGT--AGAACCTGCTGAGGATCATACCAAATTTCCTCAGCCTC
Abbassa_amh_5'UTR             29 ------------------------------------------------------------
Abbassa_amhy_h1tg000212l   11840 GA-CTTCACAAACCGACGGGAAAGAAACGGTTAATAAATACAC------------GCAAA

                                 ....:12850....:12860....:12870....:12880....:12890....:12900
Abbassa_amh_h2tg000119l    11760 CTCAGGAAATGCAGCCATTCCTGAGCCTTCTTGACCAGCTGTGTGGTGTTCAGTGTCCAG
Abbassa_amhΔy_h1tg000212l  11340 CTCAGGAAATGCAGCCGTTCCTGAGCCTTCTTGACCAGCTGTGTGGTGTTCAGTGTCCAG
Abbassa_amh_5'UTR             29 ------------------------------------------------------------
Abbassa_amhy_h1tg000212l   11887 CTCACTGAACTCTGTTGTTCTTGGACATGTTGATTCAGCTGTTCGCTGAACCTTAACAGA

                                 ....:12910....:12920....:12930....:12940....:12950....:12960
Abbassa_amh_h2tg000119l    11820 GTGATGTCCACAGAAATGTAGACGCCCAGGTATCTAAAGCTGCTCACCCTCTCCACTTCA
Abbassa_amhΔy_h1tg000212l  11400 GTGATGTCCACAGAAATGTAGACGCCCAGGTATCTAAAGCTGCTCACCCTCTCCACTTCA
Abbassa_amh_5'UTR             29 ------------------------------------------------------------
Abbassa_amhy_h1tg000212l   11947 GCAG---CCACAGTGA----------CGGGTGGCTGAAGAGCACCATTTCATTTTTTTTA

                                 ....:12970....:12980....:12990....:13000....:13010....:13020
Abbassa_amh_h2tg000119l    11880 AGGCCCCGGATAAACCGCGGCTGGTGAGGCCTCCTCTTCTTCTTCGTGTCCACTATCATC
Abbassa_amhΔy_h1tg000212l  11460 AGGCCCCGGATAAACCGCGGCTGGTGAGGCCTCCTCTTCTTCTTCGTGTCCACTATCATC
```

```
Abbassa_amh_5'UTR             29 ------------------------------------------------------------
Abbassa_amhy_h1tg000212l   11994 AAACAGATGTTAATGTACAA-----GACGCTGCATTGCATTATTGGCACCCACAGGCATG

                                 ....:13030....:13040....:13050....:13060....:13070....:13080
Abbassa_amh_h2tg000119l    11940 TCCTTTGTCT-TGTCAGTGTTGAGGGTGAGGTTGTTGTCCTCACACCATGACACCAGACC
Abbassa_amhΔy_h1tg000212l  11520 TCTTTT------------------------------------------------------
Abbassa_amh_5'UTR             29 ------------------------------------------------------------
Abbassa_amhy_h1tg000212l   12049 TTATTAATTTATCCTCAAAATAAGGCCAAAATCAACATCCGCAGATGTTTTCCTCATTCC

                                 ....:13090....:13100....:13110....:13120....:13130....:13140
Abbassa_amh_h2tg000119l    11999 GGCCACCTCTCTCCTGTAAGCCGCTTCGTCCCCTCCAGTGATGCGACGGATCACTGCAGT
Abbassa_amhΔy_h1tg000212l  11526 ------------------------------------------------------------
Abbassa_amh_5'UTR             29 ------------------------------------------------------------
Abbassa_amhy_h1tg000212l   12109 TCTCATCCTT--------------------------------------------------

                                 ....:13150....:13160....:13170....:13180....:13190....:13200
Abbassa_amh_h2tg000119l    12059 GCCATCTGAGAACTTCAAAATGATGTTATCTTTGTTGGAGGTGACACAGTCGTGAGCGTA
Abbassa_amhΔy_h1tg000212l  11526 ------------------------------------------------------------
Abbassa_amh_5'UTR             29 ------------------------------------------------------------
Abbassa_amhy_h1tg000212l   12119 ------------------------------------------------------------

                                 ....:13210....:13220....:13230....:13240....:13250....:13260
Abbassa_amh_h2tg000119l    12119 CAGGGTGTAGAGGATGGGACTGAGGACTTTTGACATTACAT------TATATATGGAAAA
Abbassa_amhΔy_h1tg000212l  11526 ------------------------------------------------------------
Abbassa_amh_5'UTR             29 ------------------------------------------------------------
Abbassa_amhy_h1tg000212l   12119 -------TGGGAGATTTGTCTGATTATTGCTAACATTAAATATCTGACATTAGCAGAAAA

                                 ....:13270....:13280....:13290....:13300....:13310....:13320
Abbassa_amh_h2tg000119l    12173 ACTGAGACGTAGAGTTGAGGTTGCACTTTCTTATATTAAGATTCACATGTAGTTAAGCTT
Abbassa_amhΔy_h1tg000212l  11526 ------------------------------------------------------------
Abbassa_amh_5'UTR             29 ------------------------------------------------------------
Abbassa_amhy_h1tg000212l   12172 ACAGAAACA---------------------------------------------------

                                 ....:13330....:13340....:13350....:13360....:13370....:13380
Abbassa_amh_h2tg000119l    12233 CTTTACTCTGACAGAGCGGGGCGTTTCACTGGTCAGGCACACTTAAGACCAAAGTGGGCT
Abbassa_amhΔy_h1tg000212l  11526 ------------------------------------------------------------
Abbassa_amh_5'UTR             29 ------------------------------------------------------------
Abbassa_amhy_h1tg000212l   12181 ------------------------------------------------------------

                                 ....:13390....:13400....:13410....:13420....:13430....:13440
Abbassa_amh_h2tg000119l    12293 GAATGTGGCCCACGATCCACAATCAGTTGGACATCCCTGCTTCAGGCTAACCATCTCCTG
Abbassa_amhΔy_h1tg000212l  11526 ------------------------------------------------------------
Abbassa_amh_5'UTR             29 ------------------------------------------------------------
Abbassa_amhy_h1tg000212l   12181 ------------------------------------------------------------

                                 ....:13450....:13460....:13470....:13480....:13490....:13500
Abbassa_amh_h2tg000119l    12353 CCCCCGGCTCCGTGTGTAAATGTACGATCACTAGTGACACTACTAAACACTGTTAAAAGG
Abbassa_amhΔy_h1tg000212l  11526 ------------------------------------------------------------
Abbassa_amh_5'UTR             29 ------------------------------------------------------------
Abbassa_amhy_h1tg000212l   12181 ------------------------------------------------------------

                                 ....:13510....:13520....:13530....:13540....:13550....:13560
Abbassa_amh_h2tg000119l    12413 CTTTTATTCTACTCTAAATAAAAGTCTGATCTGTTTTTATAAGACATATTATGAG-CATA
Abbassa_amhΔy_h1tg000212l  11526 -----ATTCTACTCTAAATAAAAGTCTGATCTGTTTTTATAAGACATATTATGAG-CATA
Abbassa_amh_5'UTR             29 ------------------------------------------------------------
Abbassa_amhy_h1tg000212l   12181 -----ACTTAACTTTAAGGAAATAGCTACAAAATATTTA----ATATTTTTTGAGACAGA

                                 ....:13570....:13580....:13590....:13600....:13610....:13620
Abbassa_amh_h2tg000119l    12472 TTAAAGAGTAGCGGACCAAGGACAGATCCCTGAGGAACTCCCTCCTGACTCAATTTTGAT
Abbassa_amhΔy_h1tg000212l  11580 TTAAAGAATAGCGGACCAAGGACAGATCCCTGAGGAAC----TCCTGACTCAATTTTGAT
Abbassa_amh_5'UTR             29 ------------------------------------------------------------
Abbassa_amhy_h1tg000212l   12232 AGGAGTCAGAGCAAGCCAAA---------CAGACAATCTAATTTATTATTTGGCTTTGAA

                                 ....:13630....:13640....:13650....:13660....:13670....:13680
Abbassa_amh_h2tg000119l    12532 TTGATCAACTAAAACATGAAACCTTCAGTACTTCACCAGTTAAACCAACTGAAC------
Abbassa_amhΔy_h1tg000212l  11636 TTGATCAACTAAAACATGAAACCTTCAGTACTTCACCAGTTAAACCAACTGAAC------
Abbassa_amh_5'UTR             29 ------------------------------------------------------------
Abbassa_amhy_h1tg000212l   12283 TTAAGCACCAACAACTCA----TTGCAGTTTGTCACAAATGTAAATATGAAAACACAGTA

                                 ....:13690....:13700....:13710....:13720....:13730....:13740
Abbassa_amh_h2tg000119l    12586 ----CATGAAGTCTTTGAATGAAAAAGCCAAAATGTGCAATGAATTCAAAGCCAGTGTTT
Abbassa_amhΔy_h1tg000212l  11690 ----CATGAAGTCTTTGAATGAAAAAGCCAAGATGTGCAATGAATTCAAAGCCAGTGTTT
Abbassa_amh_5'UTR             29 ------------------------------------------------------------
```

```
Abbassa_amhy_h1tg000212l    12339 AATACAGTAAATC-----ATGAACATGGCAGGGGTGCCAACTAACAC-AAGCAAGGGCTT

                                  ....:13750....:13760....:13770....:13780....:13790....:13800
Abbassa_amh_h2tg000119l     12642 GCAAA--TGCAGTTTTTGAATCGTTATTTTCAGAGTCCAATTTTAAAAAACTGAAAAAAT
Abbassa_amhΔy_h1tg000212l   11746 GCAAA--TGCAGTTTTTGAATCGTTATTTTCAGAGTCCAATTTTAAAAAACTGAAGAAAT
Abbassa_amh_5′UTR              29 ------------------------------------------------------------
Abbassa_amhy_h1tg000212l    12393 GTACATCTTCTGCTATGTGAGCTTAATATCCAGTGCGAAGTT-----AGACCAAAAAAGT

                                  ....:13810....:13820....:13830....:13840....:13850....:13860
Abbassa_amh_h2tg000119l     12700 TAACTATTACAGCTCATAAAAACAT-TACAGCAGCACCC---------------------
Abbassa_amhΔy_h1tg000212l   11804 TAACTATTACAGCTCATAAAAACAT-TACAGCAGCACCCAGAGTCATTTTAGCATTTCTA
Abbassa_amh_5′UTR              29 ------------------------------------------------------------
Abbassa_amhy_h1tg000212l    12448 TCCTCATTTTGTTACGCAACCAAATATACTGCAACAAATACAGAATTTC-----------

                                  ....:13870....:13880....:13890....:13900....:13910....:13920
Abbassa_amh_h2tg000119l     12738 ---------------TAACTATTGCAAACCAG---GAAATAATCTGGTTTATAAG--GAC
Abbassa_amhΔy_h1tg000212l   11863 AAATTATAAGAAAAGTAACTATTGCAAACCAG---GAAATAATCTGGTTTATAAG--GAC
Abbassa_amh_5′UTR              29 ------------------------------------------------------------
Abbassa_amhy_h1tg000212l    12497 -ACCGGGGGAGGAAATAACGCATGCACACAAAAGCAAAGGTGCTTGACGTCTGAGCAGAG

                                  ....:13930....:13940....:13950....:13960....:13970....:13980
Abbassa_amh_h2tg000119l     12778 TGCAGTCTTTTAACACTTTTTATAT--------------------------TAATGTTTTT
Abbassa_amhΔy_h1tg000212l   11918 TGCAGTCTTTTAACACTTTTTATAT--------------------------TAATGTTTTT
Abbassa_amh_5′UTR              29 ------------------------------------------------------------
Abbassa_amhy_h1tg000212l    12556 TTCAATGAACTGGCTCTGTGTGTGTGTCTGAGTGTGTGTGACTCGTCAAGTGAAACTTTG

                                  ....:13990....:14000....:14010....:14020....:14030....:14040
Abbassa_amh_h2tg000119l     12813 AAGTACAAAATATACTGCTATTTGTTTTATTTTGCTTAATTTTCATTCAGTTAATATCTT
Abbassa_amhΔy_h1tg000212l   11953 AAGTACAAAATATACTGCAATTTGTTTTATTTTGCTTAATTTTCATTCAGTTAATATCTT
Abbassa_amh_5′UTR              29 ------------------------------------------------------------
Abbassa_amhy_h1tg000212l    12616 GGGGGGGAAGTAGATCCTGGTTTATCCTG--TCGTTAAATGATAATTC------TTTTTT

                                  ....:14050....:14060....:14070....:14080....:14090....:14100
Abbassa_amh_h2tg000119l     12873 TTTCTAATTTTTAGAGAAGCTTTGTGG------------------AAAAGAAACTGTTCG
Abbassa_amhΔy_h1tg000212l   12013 TTTCTAATTTTTAGAGAAGCTTTGTGG------------------AAAAGAAACTGTTCG
Abbassa_amh_5′UTR              29 ------------------------------------------------------------
Abbassa_amhy_h1tg000212l    12668 TTTCCTGATTTTGCTGAATCCCAGTTGCTATAATTGGCACCAGAACAAAGAAACTACATA

                                  ....:14110....:14120....:14130....:14140....:14150....:14160
Abbassa_amh_h2tg000119l     12915 GGAGCAAAATATTTCACAGTTCCAATCCG--------CTTTAGTTTGTTTCATAACTGAA
Abbassa_amhΔy_h1tg000212l   12055 GGAGCAAAATATTTCACAGTTCCAATCCG--------CTTTAGTTTGTTTCATAACTGAA
Abbassa_amh_5′UTR              29 ------------------------------------------------------------
Abbassa_amhy_h1tg000212l    12728 TGAACAGATTAGATAGCACCTTAAATTCAAACACAATATTTAAGTTCACTCAGAGCTGGA

                                  ....:14170....:14180....:14190....:14200....:14210....:14220
Abbassa_amh_h2tg000119l     12967 ---------------ATGTCATTAATTTATTTTCATCTATGTCAGATGAGAAATGAA---
Abbassa_amhΔy_h1tg000212l   12107 ---------------ATGTCATTAATTTATTTTCATCTATGTCAGATGAGAAATGAA---
Abbassa_amh_5′UTR              29 ------------------------------------------------------------
Abbassa_amhy_h1tg000212l    12788 CAACAGCCAGACTGTAAAACACTACTCCACACACAAACATGACCAGT-ACCTATGAATTC

                                  ....:14230....:14240....:14250....:14260....:14270....:14280
Abbassa_amh_h2tg000119l     13009 ---TCAGATTGTTCATCA--------ACTGTGTG-GTCTGCCTGCAT-----------TT
Abbassa_amhΔy_h1tg000212l   12149 ---TCAGATTGTTCATCA--------ACTGTGTG-GTCTGCCTGCAT-----------TT
Abbassa_amh_5′UTR              29 ------------------------------------------------------------
Abbassa_amhy_h1tg000212l    12847 TGCTCAGGGTGACCATAGTGGTGGTTAGTGTGTGTGTTTGTGTGTGTGTGTGTGTGTGTG

                                  ....:14290....:14300....:14310....:14320....:14330....:14340
Abbassa_amh_h2tg000119l     13046 TGGGTGAACTTACCATTAAAGGTCATGGGAAGAGAATGGGTTATCTTAATGAACCCACAC
Abbassa_amhΔy_h1tg000212l   12186 TGGGTGAACTTACCATTAAAGGTCATGGGAAGAGAATGGGTTATCTTAATGAACCCACAC
Abbassa_amh_5′UTR              29 ------------------------------------------------------------
Abbassa_amhy_h1tg000212l    12907 TGTGTGTGTGTGTGTGTGTGTGTGTGAGAGAGAGA-----------CAAAGATCCCA---

                                  ....:14350....:14360....:14370....:14380....:14390....:14400
Abbassa_amh_h2tg000119l     13106 TGGAGATCTGGAAGCATTCACGCTGGAAGACGAGCAGGAAACATGTTGTCAGAGGGAGAC
Abbassa_amhΔy_h1tg000212l   12246 TGGAGATCTGGAAGCATTCACGCTGGAAGACGAGCAGGAAACATGTTGTCAGAGGGAGAC
Abbassa_amh_5′UTR              29 ------------------------------------------------------------
Abbassa_amhy_h1tg000212l    12953 -----GTACAGAAGC---CGGTCTTGGAGGCGATTGGGTAGTGCAGAGTTTGAAGCACCA
```

```
                                    ....:14410....:14420....:14430....:14440....:14450....:14460
Abbassa_amh_h2tg000119l       13166 CTGAGTGGCTTCGAGGAGCTGTTACTCCCCTCAGCATTCTCACAGCTGCAGGCTTGTTGT
Abbassa_amhΔy_h1tg000212l     12306 CTGAGCGGCTTCGAGGAGCTGTTACTCCCCTCAGCATTCTCACAGCTGCAGGCTTGTTGT
Abbassa_amh_5'UTR                29 ------------------------------------------------------------
Abbassa_amhy_h1tg000212l      13005 GTGAATGAGGAGAGTGCCTTGGTCCACACCTGGACATTTGCATAGTT---TATTTAATGT

                                    ....:14470....:14480....:14490....:14500....:14510....:14520
Abbassa_amh_h2tg000119l       13226 GCAGACAGTC---------------AAGGTCTCGGAGCCATTTGTGGACGTTTTCCCCCG
Abbassa_amhΔy_h1tg000212l     12366 GCAGACAGTC---------------AAGGTCTCGGAGCCATTTGTGGACGTTTTTCCCCG
Abbassa_amh_5'UTR                29 ------------------------------------------------------------
Abbassa_amhy_h1tg000212l      13062 TGAGGTAGTTTTCATCAATTACCTGAAACATTCAAAGACAGACGAGGACATC--------

                                    ....:14530....:14540....:14550....:14560....:14570....:14580
Abbassa_amh_h2tg000119l       13271 GTGATAAAGCCAAGGTTCACCATACCTTATAAATC---ATCAGTGAGGAATGGCAGGAAA
Abbassa_amhΔy_h1tg000212l     12411 GTGATAAAGCCAAGGTTCACCATACCTTATAAATC---ATCAGTGAGGAATGGCAGGAAA
Abbassa_amh_5'UTR                29 ------------------------------------------------------------
Abbassa_amhy_h1tg000212l      13114 ATGAGTGATTCTAAGGCTTCCAGAAATAACACAGCAAAAACACAGACAGCTGATTGAGGA

                                    ....:14590....:14600....:14610....:14620....:14630....:14640
Abbassa_amh_h2tg000119l       13328 TGTTCATTGTATCAAGGCGGTGTTTGAAGC----------------ACACCAGC----CT
Abbassa_amhΔy_h1tg000212l     12468 TGTTCATTGTATCAAGGCGGTGTTTGAAGC----------------ACACCAGC----CT
Abbassa_amh_5'UTR                29 ------------------------------------------------------------
Abbassa_amhy_h1tg000212l      13174 TCTAAACCCTAACAAGTTCAAAATCAAAGTCTTTACTTTTAATCTAACACCTACTTCTCT

                                    ....:14650....:14660....:14670....:14680....:14690....:14700
Abbassa_amh_h2tg000119l       13368 GGATGTTTTCAGTCACA-----------------CTTGTGGGAACCAGGAATTGCACTTT
Abbassa_amhΔy_h1tg000212l     12508 GGATGTTTTCAGTCACA-----------------CTTGTGGGAACCAGGAATTGCACTTT
Abbassa_amh_5'UTR                29 ------------------------------------------------------------
Abbassa_amhy_h1tg000212l      13234 TACCTTGTTCAATCAGACAAACAGCTTGCAGAAGCACATGGAAAACAGGCATCCTGTCTC

                                    ....:14710....:14720....:14730....:14740....:14750....:14760
Abbassa_amh_h2tg000119l       13411 GATGAAAAAGTA-------------ATCAACATCTACACTCC-------TGTT------T
Abbassa_amhΔy_h1tg000212l     12551 GATGAAAAAGTA-------------ATCAACATCTACACTCC-------TGTT------T
Abbassa_amh_5'UTR                29 ------------------------------------------------------------
Abbassa_amhy_h1tg000212l      13294 TCTCACTCCCTACCAGCCTCTCTCGCTCACCCTCTATCCACCCTCAGATTGTTCTCCGCC

                                    ....:14770....:14780....:14790....:14800....:14810....:14820
Abbassa_amh_h2tg000119l       13445 CTCCCCATCATGTATTCACCTCTCTACAGTAGCACACTTAAGTTTTCCAGGTTGGAAAAA
Abbassa_amhΔy_h1tg000212l     12585 CTCCCCATCATGTATTCACCTCTCTACAGTAGCACACTTAAGTTTTCCAGGTTGGAAAAA
Abbassa_amh_5'UTR                29 ------------------------------------------------------------
Abbassa_amhy_h1tg000212l      13354 CTTCCCCTCTCCTCTCCCCTTCCCCTCAGCCCCAAACC----CTATCTAGGTCGGGAGCT

                                    ....:14830....:14840....:14850....:14860....:14870....:14880
Abbassa_amh_h2tg000119l       13505 GGACAAAAACACAGAAGACATACATTTCTGGGGAATCTCAT-----CTGTCCATGTTGTA
Abbassa_amhΔy_h1tg000212l     12645 GGACAAAAACACAGAAGACATACATTTCTGGGGAATCTCAT-----CTGTCCATGTTGTA
Abbassa_amh_5'UTR                29 ------------------------------------------------------------
Abbassa_amhy_h1tg000212l      13410 GGGGAG---------------------CTGGGAGGTGGTGTAGGTACCATGCATGCTCTG

                                    ....:14890....:14900....:14910....:14920....:14930....:14940
Abbassa_amh_h2tg000119l       13560 AGGTTGTATCCTCATTACCCCGAAGCCAAAGG------CGGTAACCGGGGAACGCCACAG
Abbassa_amhΔy_h1tg000212l     12700 AGGTTGTATCCTCATTACCCCGAAGCCAAAGG------CGGTAACCAGGGAACGCCACAG
Abbassa_amh_5'UTR                29 ------------------------------------------------------------
Abbassa_amhy_h1tg000212l      13449 AGTCC---TCCACATGCTCCCAGCACCACCAGAACTGCTGGTCGGTGGCGGAGGGGGCAG

                                    ....:14950....:14960....:14970....:14980....:14990....:15000
Abbassa_amh_h2tg000119l       13614 GCCTGGCTTCAGGTGGAAAGCTTCAGTGC--ACGGACCTTTTGGGAGGCTTTCTCTGAGG
Abbassa_amhΔy_h1tg000212l     12754 GCCTGGCTTCAGGTGGAAAGCTTCAGTGC--ACGGGCCTTTTGGGAGGCTTTCTCTGAGG
Abbassa_amh_5'UTR                29 ------------------------------------------------------------
Abbassa_amhy_h1tg000212l      13506 A--------CGGGGGGAGTAGTGCACTGCAAATGGAC-----GAGAG---------GAGG
```

```
                                  ....:15010....:15020....:15030....:15040....:15050....:15060
Abbassa_amh_h2tg000119l     13672 CATCACAGGCATTCGACACAGAATAAGGTAACACGTCTGT--------GAACTGAAGCAG
Abbassa_amhΔy_h1tg000212l   12812 CATCACAGGCATTCGACACAGAATAAGGTAACACGTCTGT--------GAACTGAAGCAG
Abbassa_amh_5’UTR              29 ------------------------------------------------------------
Abbassa_amhy_h1tg000212l    13544 AGGCAGAGGAGGCTGAGAGAGAGGAAGAGGAGAGGGATGCAGCAGCAGAGACAGAAGCAG

                                  ....:15070....:15080....:15090....:15100....:15110....:15120
Abbassa_amh_h2tg000119l     13724 CAGAAGCTGTTTGTTTTTACAGACCTCACTGGACCATTA-ATCCTCAAGTTAGTGACTGA
Abbassa_amhΔy_h1tg000212l   12864 CAGAAGCTGTTTGTTTTTACAGACCTCACTGGACCATTA-ATCCTCAAGTTAGTGACTGA
Abbassa_amh_5’UTR              29 ------------------------------------------------------------
Abbassa_amhy_h1tg000212l    13604 CAGAGGTAGAGGATG-----AGATGGAGAGGGATGATTGTGCTCTCTGGTGGTGGAGAGA

                                  ....:15130....:15140....:15150....:15160....:15170....:15180
Abbassa_amh_h2tg000119l     13783 AGCAGCTCTCTTATGC-AGGACTACATTATGTTTATTATAACGTTCTCCATTTCTTTTTA
Abbassa_amhΔy_h1tg000212l   12923 AGCAGCTCTCTTATGC-AGGACTACATT--GTTTATTATAACGTTCTCCATTTCTTTTTA
Abbassa_amh_5’UTR              29 ------------------------------------------------------------
Abbassa_amhy_h1tg000212l    13659 GGAGAGGGACAGACGCGAGGAGTGGAGGGAGTCTGATCGAG-------------------

                                  ....:15190....:15200....:15210....:15220....:15230....:15240
Abbassa_amh_h2tg000119l     13842 AAGCTGACATATTTTTTAAAACAAT-----GTCATTAATTGTGGTTTTTAATCATCCATC
Abbassa_amhΔy_h1tg000212l   12980 AAGCTGACATATTTTTTAAAACAAT-----GTCATTAATTGTGGTTTTTAATCATCCATC
Abbassa_amh_5’UTR              29 ------------------------------------------------------------
Abbassa_amhy_h1tg000212l    13700 -----------------AAGACGCTGGGGAGTCAGAGTGCGTGGATGTCGATGAGGAGGG

                                  ....:15250....:15260....:15270....:15280....:15290....:15300
Abbassa_amh_h2tg000119l     13897 AATTTCATATCCATCTCTCTGATTCAGGCTTGCAGGAGGGCA------------------
Abbassa_amhΔy_h1tg000212l   13035 AATTTCATATCCATCTCTCTGATTCAGGCTTGCAGGAGGGCA------------------
Abbassa_amh_5’UTR              29 ------------------------------------------------------------
Abbassa_amhy_h1tg000212l    13743 GACGCTATG---------------CAGGGGTGGAGGAGGGGACAGGGAGAGCGTGGGAGG

                                  ....:15310....:15320....:15330....:15340....:15350....:15360
Abbassa_amh_h2tg000119l     13939 ----GGACCACAGTGAACAGGCT--GTTAGTCTTTCTCAGGGCTGACACACACA------
Abbassa_amhΔy_h1tg000212l   13077 ----GGACCACAGTGAACAGGCT--GTTAGTCTTTCTCAGGGCTGA--------------
Abbassa_amh_5’UTR              29 ------------------------------------------------------------
Abbassa_amhy_h1tg000212l    13788 AGAGGGGCTATGCAGGACCGACTGCGTGCGGCTGTGCTGGGCCTGGGATAACGAGAGGGG

                                  ....:15370....:15380....:15390....:15400....:15410....:15420
Abbassa_amh_h2tg000119l     13987 CACACACACACACACACACACCCATCAC---------------------TAATGATGCTG
Abbassa_amhΔy_h1tg000212l   13117 CACACACACACACACACACACCCATCAC---------------------TAATGATGCTG
Abbassa_amh_5’UTR              29 ------------------------------------------------------------
Abbassa_amhy_h1tg000212l    13848 CATAGGCGAGGACACGGGCGCCAGGCGCGTGGAGGCTGGCTGGGGAGTGGTGTGTTGCTG

                                  ....:15430....:15440....:15450....:15460....:15470....:15480
Abbassa_amh_h2tg000119l     14026 ACTTCTCAAAAACTCGAGTATCTTATACAGAGAAGCATTTTTTAATCAATTTATTTTACA
Abbassa_amhΔy_h1tg000212l   13156 ACTTCTCAAAAACTCGAGTATCTTATACAGAGACGCATTTTTTAATCAATTTATTTTACA
Abbassa_amh_5’UTR              29 ------------------------------------------------------------
Abbassa_amhy_h1tg000212l    13908 GGTTTGGGAGGA----GGTAGAGGAAGAGGAGGAGGAGAGACTGAGCAGAGAGTTG----

                                  ....:15490....:15500....:15510....:15520....:15530....:15540
Abbassa_amh_h2tg000119l     14086 TCGCAGGCCACATGCAGC--------CCACTTTGAACTTTTGTGAACCGGACCAGTGAAG
Abbassa_amhΔy_h1tg000212l   13216 TCGCAGGCCACATGA-------------ACTTTGAACTTTTGTGAACCGGACCAGAGAAG
Abbassa_amh_5’UTR              29 ------------------------------------------------------------
Abbassa_amhy_h1tg000212l    13960 ----AGGCCTCCTGAGGTGGGGAGCGTTGCCCCAAACTGATGGGAGGACACTGAGGGAA-

                                  ....:15550....:15560....:15570....:15580....:15590....:15600
Abbassa_amh_h2tg000119l     14138 CTTCCCTTTCTGTCTCTGTGAAGAGGTTTACCTACACATTTGACCTCAGAGACGT-----
Abbassa_amhΔy_h1tg000212l   13263 CTTCCCTTTCTGTCTCTGTGAAGAGGTTTACCTACACATTTGACCTCAGAGACGT-----
Abbassa_amh_5’UTR              29 ------------------------------------------------------------
Abbassa_amhy_h1tg000212l    14015 -------------CCACGTGGTGAAATATACCTGCAAATCATACAGCAGAAATGGAGGGA
```

```
                               ....:15610....:15620....:15630....:15640....:15650....:15660
Abbassa_amh_h2tg000119l   14193 ----------------------------GTGGAAACAGGA-AGTGCAGTTTCAACCA---
Abbassa_amhΔy_h1tg000212l 13318 ----------------------------GTGGAAACAGGA-AGTGCAGTTTCAACCA---
Abbassa_amh_5'UTR            29 ------------------------------------------------------------
Abbassa_amhy_h1tg000212l  14062 GGAGGAGGAGGAGGAGGAGGGCGAGAGAGTGGAGACAAGAGAGAGCAGTGGCAAAAAGAG

                               ....:15670....:15680....:15690....:15700....:15710....:15720
Abbassa_amh_h2tg000119l   14221 ----------------------------------------------------------CA
Abbassa_amhΔy_h1tg000212l 13346 ----------------------------------------------------------CA
Abbassa_amh_5'UTR            29 ------------------------------------------------------------
Abbassa_amhy_h1tg000212l  14122 CAGGGGTGGGGGACAGAGAGAGAAGCAAGGGCTGAGCCAGGTTGTGCAAAGCGCCAAGCA

                               ....:15730....:15740....:15750....:15760....:15770....:15780
Abbassa_amh_h2tg000119l   14223 TATCT---------------------------GTTTTTCTACT----------------C
Abbassa_amhΔy_h1tg000212l 13348 TATCT---------------------------GTTTTTCTACT----------------C
Abbassa_amh_5'UTR            29 ------------------------------------------------------------
Abbassa_amhy_h1tg000212l  14182 AGTCTGAGACAAGCACACAGAAAGCAAACAGAGTCAGTCCACTGGAAATGTAACGGAGCC

                               ....:15790....:15800....:15810....:15820....:15830....:15840
Abbassa_amh_h2tg000119l   14240 AATTAAGAAGTGCTGCTGTAATGAACTGAAGAAA----------TCTGTCAGCACAACAG
Abbassa_amhΔy_h1tg000212l 13365 AATTAAGAAGTGCTGCTGTAATGAACTGAAGAAA----------TCTGTCAGCACAACAG
Abbassa_amh_5'UTR            29 ------------------------------------------------------------
Abbassa_amhy_h1tg000212l  14242 AATTCAGCGCAGCCAGTGTAACG--CTGCAGGAAGTGAAATGACTCTGATGCACTAACAC

                               ....:15850....:15860....:15870....:15880....:15890....:15900
Abbassa_amh_h2tg000119l   14290 ATACTGTGGAC---------CAGCTGTA----------AAAAACAAACTGCTA-------
Abbassa_amhΔy_h1tg000212l 13415 ATACTGTGGAC---------CAGCTGTA----------AAAAACAAACTGCTA-------
Abbassa_amh_5'UTR            29 ------------------------------------------------------------
Abbassa_amhy_h1tg000212l  14300 ACCCGGGGAAAACGCACACGCAGCAGCGTGACCTGCATGAAATCAAACGGTCACTCGCGC

                               ....:15910....:15920....:15930....:15940....:15950....:15960
Abbassa_amh_h2tg000119l   14324 --------TAGTAGATTAAAAAAAA-ATTTCTTTAAGGGTTTATATATTTGTTGGTTACT
Abbassa_amhΔy_h1tg000212l 13449 --------TAGTAGATTAAAAAAAAAATTTCTTTAAGGGTTTATATATTTGTTGGTTACT
Abbassa_amh_5'UTR            29 ------------------------------------------------------------
Abbassa_amhy_h1tg000212l  14360 TTTGTTATTAATGATTTCAATGAAAGTTTGTGAAAATGTCACATTTTCCGAGG-----

                               ....:15970....:15980....:15990....:16000....:16010....:16020
Abbassa_amh_h2tg000119l   14375 TTGTCTTTTCATTAGGAAAAACTATCAAACTTGTGGAAGTTCACATTTTGCCGTCGAGTG
Abbassa_amhΔy_h1tg000212l 13501 TTGTCTTTTCATTAGGAAAAACTATCAAACTTGTGGAAGTTCACATTTTGCCGTCGAGCG
Abbassa_amh_5'UTR            29 ------------------------------------------------------------
Abbassa_amhy_h1tg000212l  14415 --GTGTCTCCACCTGCATTACCTAAACAGCCAGTCCACGTCAAAGTATGGCAGCACAGCG

                               ....:16030....:16040....:16050....:16060....:16070....:16080
Abbassa_amh_h2tg000119l   14435 GGCCAGATTGGAGTCTTTGCCGGGCCGACTTGGCATTACAAGGTAAATGCCGCCTACCTG
Abbassa_amhΔy_h1tg000212l 13561 GGCCAGATTGGAGTCTTTGCCGGGCCGACTTGGCATTACAAGGTAAATGCCGCCTACCTG
Abbassa_amh_5'UTR            29 ------------------------------------------------------------
Abbassa_amhy_h1tg000212l  14473 AGTCACAGT------TTTGCCTGCTCACCT------------GTCCTTGCTGCCCTCGGG

                               ....:16090....:16100....:16110....:16120....:16130....:16140
Abbassa_amh_h2tg000119l   14495 TGAAGTCTACTAACTGAGGATTTAGGTAGCGGGCTCCAAACATGGTTCAGGTATGAAGA
Abbassa_amhΔy_h1tg000212l 13621 TGAAGTCT----ACTGAGGATTTAGGTAGCGGGCTCCAAACATGGTTCAGGTATGAAGA
Abbassa_amh_5'UTR            29 ------------------------------------------------------------
Abbassa_amhy_h1tg000212l  14515 CAAAGCTC------------------CAGGAGGGATTTCAAC---GTACAGGTTTCCTCC

                               ....:16150....:16160....:16170....:16180....:16190....:16200
Abbassa_amh_h2tg000119l   14555 TATCAGCATGACACCCTCTTCCGTCACGG----------CAAACCAAATGGACTCAGACA
Abbassa_amhΔy_h1tg000212l 13677 TATCAGCATGACACCCTCTTCCGTCACGG----------CAAACCAAATGGACTCAGACA
Abbassa_amh_5'UTR            29 ------------------------------------------------------------
Abbassa_amhy_h1tg000212l  14554 TTTCAGCGC----CGCCCTCCAGACACAGTGCTTTCCTTCAGAGCGCCGGGGCTCAAAGA
```

```
                                 ....:16210....:16220....:16230....:16240....:16250....:16260
Abbassa_amh_h2tg000119l     14605 -----AAGTTCACTCC---------------------------AAACGTAAACTCAATAT
Abbassa_amhΔy_h1tg000212l   13727 -----AAGTTCACTCC---------------------------AAACGTAAACTCAATAT
Abbassa_amh_5'UTR              29 ------------------------------------------------------------
Abbassa_amhy_h1tg000212l    14610 GTAAGAAGCGCACTCTGCTATTGGCAGGGGAAGGCTCCTCAAGAACCAGCACCTGAGCAT

                                 ....:16270....:16280....:16290....:16300....:16310....:16320
Abbassa_amh_h2tg000119l     14633 TATAAGAAA---CATTTTATTAATTATCATAAATCGCAGGGGAG-AAAAAAAAACATTAA
Abbassa_amhΔy_h1tg000212l   13755 TATAAGAAA---CATTTTATTAATTATCATAAATCGCAGGGGAGAAAAAAAAAACATTAA
Abbassa_amh_5'UTR              29 ------------------------------------------------------------
Abbassa_amhy_h1tg000212l    14670 CCTGGGAAAGTGTAGTTTAGGTTATGTTAAAAATGAAACTGCACATACCAGCAAGGACAA

                                 ....:16330....:16340....:16350....:16360....:16370....:16380
Abbassa_amh_h2tg000119l     14689 TTCCATTTTTTCATTATAACATCTTAGGATAAAAGCACCAACATCAAACAGTAACAATGT
Abbassa_amhΔy_h1tg000212l   13812 TTCCATTTTTTCATTATAACATCTTAGGATAAAAGCACCAACATC-AACAGTAACAATGT
Abbassa_amh_5'UTR              29 ------------------------------------------------------------
Abbassa_amhy_h1tg000212l    14730 TTCTGTATTT---------CTTTTTAAGCTTTCATCACTTACAG-----AATAGAACAGC

                                 ....:16390....:16400....:16410....:16420....:16430....:16440
Abbassa_amh_h2tg000119l     14749 TTCCATGAAAAACCGAGGGCGTTTTGGAAAGTAGGTTTGCCATTATTGCACCTCAAACCT
Abbassa_amhΔy_h1tg000212l   13871 TTCCATGAAAAACCGAGGGCGTTTTGGAAAGTAGGTTTGCCATTATTGCACCTCAAACCT
Abbassa_amh_5'UTR              29 ------------------------------------------------------------
Abbassa_amhy_h1tg000212l    14776 TTAGCTGAAAGATTGCGATCCCGTTGG---------------TCATTCCCTCGCCCACCT

                                 ....:16450....:16460....:16470....:16480....:16490....:16500
Abbassa_amh_h2tg000119l     14809 GCCCTGGCTTCACCAATCAGAGA---GCACTATACACATCCTGTAGGTC----ACGCGTG
Abbassa_amhΔy_h1tg000212l   13931 GCCCTGGCTTCACCAATCAGAGA---GCACTATACACATCCTGTAGGTC----ACGCGTG
Abbassa_amh_5'UTR              29 ------------------------------------------------------------
Abbassa_amhy_h1tg000212l    14821 GGGAT--CTCCAGGGGTGGGAGAGGGGCATCAGGAGCACCACAGAGGCCCCGGACACGGG

                                 ....:16510....:16520....:16530....:16540....:16550....:16560
Abbassa_amh_h2tg000119l     14862 TCCTTGAATGTTACCCCCAATTCAAAGAGGGATTCAAACAAGACACCGAACCTCCTCCTT
Abbassa_amhΔy_h1tg000212l   13984 TCCTTGAATGTTACCCCCAATTCAAAGAGGGATTCAAACAAGACACCGAACCTCCTCCTT
Abbassa_amh_5'UTR              29 ------------------------------------------------------------
Abbassa_amhy_h1tg000212l    14879 TTACTGCA---------------ACAGCGGGAGCACCTCAGAGGACTGGAGATACT----

                                 ....:16570....:16580....:16590....:16600....:16610....:16620
Abbassa_amh_h2tg000119l     14922 AGAGACGCAAATCTAGACAGAGGCATTTGTGAGATGAGGAAGATGCTGAAAGTCGTTGCC
Abbassa_amhΔy_h1tg000212l   14044 AGAGACGCAAATCTAGACAGAGGCATTTGTGAGATGAGGAAGATGCTGAAAGTCGTTGCC
Abbassa_amh_5'UTR              29 ------------------------------------------------------------
Abbassa_amhy_h1tg000212l    14920 -GGAAAGCGAAACAAACAAGTGGTTTATGTCACACCACCACATTACTGA-------TGCC

                                 ....:16630....:16640....:16650....:16660....:16670....:16680
Abbassa_amh_h2tg000119l     14982 ATTTCACAGCTTCT--CGAGATCAGATCGCTCATAAAGCTCGGACTGATTGTCGGGGGTG
Abbassa_amhΔy_h1tg000212l   14104 ATTTCACAGCTTCT--CGAGATCAGATCGCTCATAAAGCTCGGACTGATTGTCGGGGGTG
Abbassa_amh_5'UTR              29 ------------------------------------------------------------
Abbassa_amhy_h1tg000212l    14972 TCACTAAAACTGTTAACAATTTTATGTCAGCTGTCAGTCAAATATTGACCACCAACTATG

                                 ....:16690....:16700....:16710....:16720....:16730....:16740
Abbassa_amh_h2tg000119l     15040 GGGGTGT----------------------ACGTGCCGGCAGGACTGTGTCTGAGGAGATC
Abbassa_amhΔy_h1tg000212l   14162 GGGGTGT----------------------ACGTGCCGGCAGGACTGTGTCTGAGGAGATC
Abbassa_amh_5'UTR              29 ------------------------------------------------------------
Abbassa_amhy_h1tg000212l    15032 AGAAACTGTTTGTTTTTTGTCTCACAAAGACACATCGAGGCTACGGCTTCTGCACACTTC

                                 ....:16750....:16760....:16770....:16780....:16790....:16800
Abbassa_amh_h2tg000119l     15078 TGATGAGGGCAGCTGATACATTTTACCTTG-------AAATACACTGACTGAAGGGTGGC
Abbassa_amhΔy_h1tg000212l   14200 TGATGAGGGCAGCTGATACATTTTACCTTG-------AAATACACTGACTGAAGGGTGGC
Abbassa_amh_5'UTR              29 ------------------------------------------------------------
Abbassa_amhy_h1tg000212l    15092 TGCTGCGGGCACTTTAAGCTTGTTCCACAGTTCTCACAGGAAACCTGCCAGAAGAACAGG
```

```
                                       ....:16810....:16820....:16830....:16840....:16850....:16860
Abbassa_amh_h2tg000119l         15131 CCTGGTCCTGAGTTGGGTCGTAGAGGGACTTTGTCATGCCACCTGAAGGTAAATTAACGT
Abbassa_amhΔy_h1tg000212l       14253 CCTGGTCCTGAGTTGGGTCGTAGAGGGACTTTGTCATGCCACCTGAAGGTAAATTAACGT
Abbassa_amh_5′UTR                  29 ------------------------------------------------------------
Abbassa_amhy_h1tg000212l        15152 CTATTTTTTTCCTCTAGCCGCTAGCCTACTTTAGCTCTTCAC-----------TCACCGC

                                       ....:16870....:16880....:16890....:16900....:16910....:16920
Abbassa_amh_h2tg000119l         15191 AACATGCAGCAGATTAGGAAACTTGTTCTTTCAAGCAGTGTGGGGAGTTCCTGAGTTAAG
Abbassa_amhΔy_h1tg000212l       14313 AACATGCAGCAGATTAGGAAACTTGTTCTTTCAAGCAGTGTGGGGAGTTCCTGAGTTAAG
Abbassa_amh_5′UTR                  29 ------------------------------------------------------------
Abbassa_amhy_h1tg000212l        15201 GTCA--CTGCAGTTTAAGAGACTGGGACAAACAATACTGATCAGCCGTTCT--AATTCAG

                                       ....:16930....:16940....:16950....:16960....:16970....:16980
Abbassa_amh_h2tg000119l         15251 ACCTTTGGCAGGATTTCGAGTAAACAGCCT----TGACTGCAAAGCTCT-----------
Abbassa_amhΔy_h1tg000212l       14373 ACCTTTGGCAGGATTTCGAGTAAACAGCCT----TGACTGCAAAGCTCT-----------
Abbassa_amh_5′UTR                  29 ------------------------------------------------------------
Abbassa_amhy_h1tg000212l        15257 CGCTGTCACAGCACTCCTCAGATTTAGCTTGCTGTGACTGCATACATTTACAGCTTTGAG

                                       ....:16990....:17000....:17010....:17020....:17030....:17040
Abbassa_amh_h2tg000119l         15296 -GAAAGATTATCTACGAATAATGTACCAGGATGCAGTATGATGCAGAACTGCACAGCTTT
Abbassa_amhΔy_h1tg000212l       14418 -GAAAGATTATCTACGAATAATGTACCAGGATGCAGTATGATGCAGAACTGCACAGCTTT
Abbassa_amh_5′UTR                  29 ------------------------------------------------------------
Abbassa_amhy_h1tg000212l        15317 CGAAGGCCTGTTCACTGGGATTCTATTAGGAACAAAAAT-------AACAGCGCCGC---

                                       ....:17050....:17060....:17070....:17080....:17090....:17100
Abbassa_amh_h2tg000119l         15355 AAAGCCAAATCACACCCCAGAAATACTTTTAAAAACTATAAAACTGCCTAAAATCTG-TA
Abbassa_amhΔy_h1tg000212l       14477 AAAGCCAAATCACACCCCAGAAATACTTTTAAAAACTATAAAACTGCCTAAAATCTG-TA
Abbassa_amh_5′UTR                  29 ------------------------------------------------------------
Abbassa_amhy_h1tg000212l        15367 -AAACACCATCACA-CACAGGGATGCTCTC-----TTACGCAAATGCCACCAATGCGCTA

                                       ....:17110....:17120....:17130....:17140....:17150....:17160
Abbassa_amh_h2tg000119l         15414 TGTAATGAATGGAAT------ACCTCCCACCTTCCTCAAAGTGCTCTGACAGGCACGTAC
Abbassa_amhΔy_h1tg000212l       14536 TGTAATGAATGGAAT------ACCTCCCACCTTCCTCAAAGTGCTCTGACAGGCACGTAC
Abbassa_amh_5′UTR                  29 ------------------------------------------------------------
Abbassa_amhy_h1tg000212l        15420 CATGTTGAGCAGAGGTGTGGAAACCAGTGTCATTCGCAGTGTGTAATATCCAACTTATGA

                                       ....:17170....:17180....:17190....:17200....:17210....:17220
Abbassa_amh_h2tg000119l         15468 ACCCCC------------GCCTCCGCCCGCCGGTGTCCCTCCAGATTAAAGATTCAAAGG
Abbassa_amhΔy_h1tg000212l       14590 ACCCCC------------GCCTCCGCCCGCCGGTGTCCCTCCAGATTAAAGATTCAAAGG
Abbassa_amh_5′UTR                  29 ------------------------------------------------------------
Abbassa_amhy_h1tg000212l        15480 TCTCACAGAGGATTGGGGATTTTAACCAAATGCAGTTACTCCCT-----AAATCCGGTAA

                                       ....:17230....:17240....:17250....:17260....:17270....:17280
Abbassa_amh_h2tg000119l         15516 TTAGACTGGGCTGTGAGAAAAGCCACCAGGTGAGCTTGTGTGTAAGATTCTTAAACCATA
Abbassa_amhΔy_h1tg000212l       14638 TTAGACTGGGCTGTGAGAAAAGCCACCAGGTGAGCTTGTGTGTAAGATTCTTAAACCATA
Abbassa_amh_5′UTR                  29 ------------------------------------------------------------
Abbassa_amhy_h1tg000212l        15535 TCAGATTAGG----AAAGGAAGCCACTTGCCATCCT----TGCAGAACCCCTTAACGCCT

                                       ....:17290....:17300....:17310....:17320....:17330....:17340
Abbassa_amh_h2tg000119l         15576 AAACCCTGCAGCACAACAGGAGGAGAGTAGTATTCATCAATAGCTGCGATTTCCTTGTTT
Abbassa_amhΔy_h1tg000212l       14698 AAACCCTGCAGCACAACAGGAGGAGAGTAGTATTCATCAATAGCTGCGATTTCCTTGTTT
Abbassa_amh_5′UTR                  29 ------------------------------------------------------------
Abbassa_amhy_h1tg000212l        15587 CCTTTCTGCGATTTAAAGAGAACACAAAGTTCTCAGTCAGTTTCTGCCTTTAGCATTCTT

                                       ....:17350....:17360....:17370....:17380....:17390....:17400
Abbassa_amh_h2tg000119l         15636 TAA---------GTGAAGGAGAGCACCGTGCTGTCTGACTAGTAGTATTAAAAACTGACA
Abbassa_amhΔy_h1tg000212l       14758 TAA---------GTGAAGGAGAGCACCGTGCTGTCTGACTAGTAGTATTAAAAACTGACA
Abbassa_amh_5′UTR                  29 ------------------------------------------------------------
Abbassa_amhy_h1tg000212l        15647 TGAGACTGACATGTGAACAGTAG-GCAGTGCTGCGCA---------ATGACGCCCTGACG
```

```
                                   ....:17410....:17420....:17430....:17440....:17450....:17460
Abbassa_amh_h2tg000119l      15687 GAAATCTGAAGACACAATCAAAAGCATAACATGTAACT-GCAATCGTATCAAACACCAA-
Abbassa_amhΔy_h1tg000212l    14809 GAAATCTGAAGACACAATCAAAAGCATAACATGTAACT-GCAATCGTATCAAACACCAA-
Abbassa_amh_5'UTR               29 ------------------------------------------------------------
Abbassa_amhy_h1tg000212l     15697 GTCATCTTAG--------TAAAAACAGCTGCTGTGAATAGTGTTTGTAGGAGACTCTGCT

                                   ....:17470....:17480....:17490....:17500....:17510....:17520
Abbassa_amh_h2tg000119l      15745 GAATAAGAAAATCA----AAACGCCATCGACAATAATTACAAAATACCTCTCAGTGCTGG
Abbassa_amhΔy_h1tg000212l    14867 GAATAAGAAAATCA----AAACACCATCGACAATAATTACAAAATACCTCTCAGTGCTGG
Abbassa_amh_5'UTR               29 ------------------------------------------------------------
Abbassa_amhy_h1tg000212l     15749 GTAAATGAACATCATTGTAAAGGACATCTGCAATAAATCTCATGCATCCATC--------

                                   ....:17530....:17540....:17550....:17560....:17570....:17580
Abbassa_amh_h2tg000119l      15801 TAACAAATGGACTAGTCTAATTCTAAAATGACTTTTTCTCTTCACTTGCACACGAGTGCA
Abbassa_amhΔy_h1tg000212l    14923 TAACAAATGGACTAGTCTAATTCTAAAATGACTTTTTCTCTTCACTTGCACACGAGTGCA
Abbassa_amh_5'UTR               29 ------------------------------------------------------------
Abbassa_amhy_h1tg000212l     15801 TAGCAGAGAATCATCTTTAATCCCATG-----TTCTAGCCTGCACTGTTAGACACTTCAA

                                   ....:17590....:17600....:17610....:17620....:17630....:17640
Abbassa_amh_h2tg000119l      15861 CGGGACAGGA--------------GTTTTATTTT------TCTATAAAGCTTTGATTATT
Abbassa_amhΔy_h1tg000212l    14983 CGGGACAGGA--------------GTTTTATTTT------TCTATAAAGCTTTGATTATT
Abbassa_amh_5'UTR               29 ------------------------------------------------------------
Abbassa_amhy_h1tg000212l     15856 TGTAGTATGACATTTGCAATGTGGGTTTTATTTTAGCTCACCTAACACGTTCTGGCT---

                                   ....:17650....:17660....:17670....:17680....:17690....:17700
Abbassa_amh_h2tg000119l      15901 ACACTAATATTCCCTTAGATATGCGCTGGCAAGTGAAAAATTAAAAAAAAACCAA-----
Abbassa_amhΔy_h1tg000212l    15023 ACACTAATATTCCCTTAGATATGCGCTGGCAAGTGAAAAATTAAAAAA--ACCAA-----
Abbassa_amh_5'UTR               29 ------------------------------------------------------------
Abbassa_amhy_h1tg000212l     15913 ---------TTCAACAAAGAAAAATTAGCAGGTTAAAAATGTAACAGTCTTTAATGTGG

                                   ....:17710....:17720....:17730....:17740....:17750....:17760
Abbassa_amh_h2tg000119l      15956 --ACAAAAACACACACGTACAAACTGAGCTTTAGTGTAATCCCACTATCAGAG-------
Abbassa_amhΔy_h1tg000212l    15076 --ACAAAA--ACACACGTACAAACTGAGCTTTAGTGTAATCCCACTATCAGAG-------
Abbassa_amh_5'UTR               29 ------------------------------------------------------------
Abbassa_amhy_h1tg000212l     15964 GGTTAAATTTACACAAGTTTAAAT--ACATTTGATTTTATCAGGTGGTCAAAAATGAAAA

                                   ....:17770....:17780....:17790....:17800....:17810....:17820
Abbassa_amh_h2tg000119l      16007 ----GAAAAAAGGAAGCCT-GTCTGATCT--CACTCACACACACACACACTAAGACAAGT
Abbassa_amhΔy_h1tg000212l    15125 ----GAAAAAAGGAAGCCT-GTCTGATCTCACACACACACACACACACACTAAGACAAGT
Abbassa_amh_5'UTR               29 ------------------------------------------------------------
Abbassa_amhy_h1tg000212l     16022 GAATGAAAAGAAAAGGCCTAGCCTTCTCT--TA--------------AGGTAACAGAGAC

                                   ....:17830....:17840....:17850....:17860....:17870....:17880
Abbassa_amh_h2tg000119l      16060 GTTTCACATCAGCTCACGCGGCGAGGTACGGCCTCGTCAGTAAGACGTCTACAGGCGGTG
Abbassa_amhΔy_h1tg000212l    15180 GTTTCACATCAGCTCACGCGGCGAGGTACGGCCTCGTCAGTAAGACGTCTACAGGCGGTG
Abbassa_amh_5'UTR               29 ------------------------------------------------------------
Abbassa_amhy_h1tg000212l     16066 TTTTTACCTTGG-------GAAGGGGGAGGCTATTAGCTGCTATA-----ATAGATGATG

                                   ....:17890....:17900....:17910....:17920....:17930....:17940
Abbassa_amh_h2tg000119l      16120 TGTGTAAAGAGGCTGACGCGCAGTTCTGTATGCACCGGTGAAGATTTTGA-------TTG
Abbassa_amhΔy_h1tg000212l    15240 TGTGTAAAGAGGCTGACGCGCAGTTCTGTATGCACCGGTGAAGATTTTGA-------TTG
Abbassa_amh_5'UTR               29 ------------------------------------------------------------
Abbassa_amhy_h1tg000212l     16114 T-------TATTTTAATGCATGCTTTATTATTGGAAGGTGTGGATGTTAAACCAGCCTTA

                                   ....:17950....:17960....:17970....:17980....:17990....:18000
Abbassa_amh_h2tg000119l      16173 TAAGGTCTGA----ACTGCCCTCGCTTGGAAACAGAAGCACCATTTTTCCTGCAGAATCT
Abbassa_amhΔy_h1tg000212l    15293 TAAGGTCTGA----ACTGCCCTCGCTTGGAAACAGAAGCACCATTTTTCCTGCAGAATCT
Abbassa_amh_5'UTR               29 ------------------------------------------------------------
Abbassa_amhy_h1tg000212l     16167 TCACACATGATAGCTATGTGTCAGCATATTTATAAAGGCATCTTTT------CAAGCTTT
```

```
                                      ....:18010....:18020....:18030....:18040....:18050....:18060
Abbassa_amh_h2tg000119l        16229 GGGTTGGTAAAAAA----ATTAAACACCAAAAACAAACAAAAA--AAACAACAATAAATC
Abbassa_amhΔy_h1tg000212l      15349 GGGTTGGTAAAAAA----ATTAAACACCAAAAACAAACAAAAA--AACCAACAATAAATC
Abbassa_amh_5'UTR                 29 ------------------------------------------------------------
Abbassa_amhy_h1tg000212l       16221 GGCTTAAAGAGAAACACTATTATATAATGTACATATATAAATATCTAACCATGACAATTA

                                      ....:18070....:18080....:18090....:18100....:18110....:18120
Abbassa_amh_h2tg000119l        16283 AAAGAAGTGAAAATACCTGAAACAAGTGATAACAGTTCTAGTACTG----------GAAT
Abbassa_amhΔy_h1tg000212l      15403 AAAGAAGTGAAAATACCTGAAACAAGTGATAACAGTTCTAGTACTG----------GAAT
Abbassa_amh_5'UTR                 29 ------------------------------------------------------------
Abbassa_amhy_h1tg000212l       16281 CATAACAATTACATAGCCGAAACAGCTGAACAC---TCTTGCACTGCCACTGCAAAGAGA

                                      ....:18130....:18140....:18150....:18160....:18170....:18180
Abbassa_amh_h2tg000119l        16333 CTGGTGGCGTTTGAGCCGC-----------------------------------------
Abbassa_amhΔy_h1tg000212l      15453 CTGGTGGCGTTTGAGCCGC-----------------------------------------
Abbassa_amh_5'UTR                 29 ------------------------------------------------------------
Abbassa_amhy_h1tg000212l       16338 TTAGGGGCTGCAGAGCGGCGTAACAAGTTAGTATTAGTTAACAGTAATAACCTTGGCTAT

                                      ....:18190....:18200....:18210....:18220....:18230....:18240
Abbassa_amh_h2tg000119l        16352 --------------CGCACCTTTGTGTAATAAACACACAGCCT-------CTGAGTTTGT
Abbassa_amhΔy_h1tg000212l      15472 --------------CGCACCTTTGTGTAATAAACACACAGCCT-------CTGAGTTTGT
Abbassa_amh_5'UTR                 29 ------------------------------------------------------------
Abbassa_amhy_h1tg000212l       16398 CACAGGTTAGTGTATACTTCTTTGGGTGATGGATGCCGAACTTTTCACAGCTATCTTTGA

                                      ....:18250....:18260....:18270....:18280....:18290....:18300
Abbassa_amh_h2tg000119l        16391 GGCTGCGGGGCATGTGTGTGTGTGTGTG-----------TGTGTGTGT------------
Abbassa_amhΔy_h1tg000212l      15511 GGCTGCGGGGCATGTGTGTGTGTGTGTG-----------TGTGTGTGTGTGTGTGTGTGT
Abbassa_amh_5'UTR                 29 ------------------------------------------------------------
Abbassa_amhy_h1tg000212l       16458 GAGTAACAGAGTCATTATCGTATTTTTGGTACTGCAAGATGTGTGCTTAATAATTTTATA

                                      ....:18310....:18320....:18330....:18340....:18350....:18360
Abbassa_amh_h2tg000119l        16428 ---------------------------TTTCAGGTGCTGATGTGCAGCCATGTTAAGGTC
Abbassa_amhΔy_h1tg000212l      15560 GTGTGTGTGTGTGTGTGTGTGTGTGTGTTTCAGGTGCTGATGTGCAGCCATGTTAAGGTC
Abbassa_amh_5'UTR                 29 ------------------------------------------------------------
Abbassa_amhy_h1tg000212l       16518 ATATAGCAATAATAATTTTCATAAATGTTCCAAAGA-----GCGCAGAAATGTCTACAGC

                                      ....:18370....:18380....:18390....:18400....:18410....:18420
Abbassa_amh_h2tg000119l        16461 CTCTGGTGGCAATGCTTTGCTCTGCTCAGCTTCAAGCCTCCTGTTACAGGGTGAA----T
Abbassa_amhΔy_h1tg000212l      15620 CTCTGGTGGCAATGCTTTGCTCTGCTCAGCTTCAAGCCTCCTGTTACAGGGTGAA----T
Abbassa_amh_5'UTR                 29 ------------------------------------------------------------
Abbassa_amhy_h1tg000212l       16573 CGGTTTCTCCAAGTCTAGGAAATGCACA----CTAAACTGCTCAAGCTGGACAAACAGCT

                                      ....:18430....:18440....:18450....:18460....:18470....:18480
Abbassa_amh_h2tg000119l        16517 TTAAGGGTCACAATGCCAGCACCAAAACACCCCTTTCACATATATATGTATATATATTTA
Abbassa_amhΔy_h1tg000212l      15676 TTAAGGGTCACAATGCCAGCACCAAAACACCCCTTTCACATATATATGTATATATATTTA
Abbassa_amh_5'UTR                 29 ------------------------------------------------------------
Abbassa_amhy_h1tg000212l       16629 CTACAGCTCAGAGGACCAAAAATAAACTGGACTTTGGAT-CGTCTCTTTAAATA---TCA

                                      ....:18490....:18500....:18510....:18520....:18530....:18540
Abbassa_amh_h2tg000119l        16577 TATACTTATATATATTTATATAGACTGTATATAGTTACTTATTTTATATACCTTTCTGTT
Abbassa_amhΔy_h1tg000212l      15736 TATACTTATATATATTTATATAGACTGTATATAGTTACTTATTTTATATACCTTTCTGTT
Abbassa_amh_5'UTR                 29 ------------------------------------------------------------
Abbassa_amhy_h1tg000212l       16685 CATGCTCAAATATTATGA---------TATATGAGAATTTTTATTATGCAACACTATA--

                                      ....:18550....:18560....:18570....:18580....:18590....:18600
Abbassa_amh_h2tg000119l        16637 TATGATGGAGATGTACAA---TTAAAGAAAACTTATGTACAAAAACAGTGTGTCTTGTTC
Abbassa_amhΔy_h1tg000212l      15796 TATGATGGAGATGTACAA---TTAAAGAAAACTTATGTACAAAAACAGTGTGTCTTGTTC
Abbassa_amh_5'UTR                 29 ------------------------------------------------------------
Abbassa_amhy_h1tg000212l       16734 ------GGATTTCCACAGTATTTAGGGCTGGTTGATATGGAATCACA-CGTGTCTTGTGT
```

```
                                 ....:18610....:18620....:18630....:18640....:18650....:18660
Abbassa_amh_h2tg000119l    16694 ACCAACAGCGGGCC---GATGTATGTTTAAAACATTAACAGGGGC---------------
Abbassa_amhΔy_h1tg000212l  15853 ACCAACAGCGGGCC---GATGTATGTTTAAAACAATAACAGGGGCCCCCGTGGCCGTCGC
Abbassa_amh_5’UTR             29 ------------------------------------------------------------
Abbassa_amhy_h1tg000212l   16787 CCCATTGTCTGCCTCCAGGTTTCCATATGGGACGGCTTCA--------------------

                                 ....:18670....:18680....:18690....:18700....:18710....:18720
Abbassa_amh_h2tg000119l    16736 -----CCCCGTGGCCGTCGCTCGGGCCCCGAGCCGGCTGCACGAGAGCACTTTCATCCAA
Abbassa_amhΔy_h1tg000212l  15910 TCGGGCCCCGTGGCCGTCGCTCGGGCCCCGAGCCGGCTGCACGAGAGCACTTTCATCCAA
Abbassa_amh_5’UTR             29 ------------------------------------------------------------
Abbassa_amhy_h1tg000212l   16827 -----CCCGGTGGTGG------GGATCTGAAGAAGTCGGAGCGATAA----------GAA

                                 ....:18730....:18740....:18750....:18760....:18770....:18780
Abbassa_amh_h2tg000119l    16791 AGGTGGCGTCTACAGTTGTACACGATGCATTC--AAAGAACGAGACAATAGCATATTAAA
Abbassa_amhΔy_h1tg000212l  15970 AGGTGGCGTCTACAGTTGTACACGATGCATTC--AAAGAACGAGACAATAGCATATTAAA
Abbassa_amh_5’UTR             29 ------------------------------------------------------------
Abbassa_amhy_h1tg000212l   16866 GGAGGAAGAAGCCAGCTGCTAATAAAGCAATCCTGAAACCCAAAATCAAAGCATAAAACA

                                 ....:18790....:18800....:18810....:18820....:18830....:18840
Abbassa_amh_h2tg000119l    16849 TATGAGAAAGGAAATGCCAAAAAACGGCTTTACAAATGTCCCTGTGTACAGCACCAAGTA
Abbassa_amhΔy_h1tg000212l  16028 TATGAGAAAGGAAATGCCAAAAAACGGCTTTACAAATGTCCCTGTGTACAGCACCAAGTA
Abbassa_amh_5’UTR             29 ------------------------------------------------------------
Abbassa_amhy_h1tg000212l   16926 TATACTGTAGCTTATGTAGCTGAATGGC--TACGAA--------CACAGAACTTTAAGTA

                                 ....:18850....:18860....:18870....:18880....:18890....:18900
Abbassa_amh_h2tg000119l    16909 CTTGACTTGTTTTTGATGTTTTCATTACAGAATAAATAAGATCAGGCATAGTCCAAACTT
Abbassa_amhΔy_h1tg000212l  16088 CTTGACTTGTTTTTGATGTTTTCATTACAGAATAAATAAGATCAGGCATAGTCCAAACTT
Abbassa_amh_5’UTR             29 ------------------------------------------------------------
Abbassa_amhy_h1tg000212l   16976 C-------ACTTTGCATTTAATTATTACAAGTTTGAGAAGGCTATATTTCATCC--TTTT

                                 ....:18910....:18920....:18930....:18940....:18950....:18960
Abbassa_amh_h2tg000119l    16969 AGCACCATTGTTTTCAGACTGTGAGTCAAGTACACACACACTAATGCATTCTACAATGTC
Abbassa_amhΔy_h1tg000212l  16148 AGCACCATTGTTTTCAGACTGTGAGTCAAGTACACACACACTAATGCATTCTACAATGTC
Abbassa_amh_5’UTR             29 ------------------------------------------------------------
Abbassa_amhy_h1tg000212l   17027 AACAGTTTAACTTTAACACT------------CAGAAACATTATTCCAATTAGAAATATT

                                 ....:18970....:18980....:18990....:19000....:19010....:19020
Abbassa_amh_h2tg000119l    17029 AACACAATTCAGGATTT------------AAAAAAAAAAAAAAAAAAAAAAAAAGGCAAA
Abbassa_amhΔy_h1tg000212l  16208 AACACAATTCAGGATTTAAAAAAAAAAAAAAAAAAAAAAAAAAAAAAAAAAAAAAGGCAAA
Abbassa_amh_5’UTR             29 ------------------------------------------------------------
Abbassa_amhy_h1tg000212l   17075 TTCAAAAT-----------------------------GACAAAA----------------

                                 ....:19030....:19040....:19050....:19060....:19070....:19080
Abbassa_amh_h2tg000119l    17077 TGGAAAGACCGAC-----------------------------------------------
Abbassa_amhΔy_h1tg000212l  16268 TGGAAAGACCGACAGCATAAATATAATCATCTAAATTATCCACAACTATACAGTCAGAGA
Abbassa_amh_5’UTR             29 ------------------------------------------------------------
Abbassa_amhy_h1tg000212l   17090 ------------------------------------------------------------

                                 ....:19090....:19100....:19110....:19120....:19130....:19140
Abbassa_amh_h2tg000119l    17090 ------------------------------------------------------------
Abbassa_amhΔy_h1tg000212l  16328 ATGAATCACATTCACAGTACAAACATTGTTTACAAAAACAAATTACACCGTTTTTTAAAA
Abbassa_amh_5’UTR             29 ------------------------------------------------------------
Abbassa_amhy_h1tg000212l   17090 ------------------------------------------------------------

                                 ....:19150....:19160....:19170....:19180....:19190....:19200
Abbassa_amh_h2tg000119l    17090 ------------------------------------------------------------
Abbassa_amhΔy_h1tg000212l  16388 AGCAAACAAAGAGGTGTGAAACAAGTCCAGTAGTTTGTTGTGGTTATGGTCAACTATGTT
Abbassa_amh_5’UTR             29 ------------------------------------------------------------
Abbassa_amhy_h1tg000212l   17090 ------------------------------------------------------------
```

```
                                 ....:19210....:19220....:19230....:19240....:19250....:19260
Abbassa_amh_h2tg000119l    17090 ------------------------------------------------------------
Abbassa_amhΔy_h1tg000212l  16448 TTCCCAGTGTGTGAGGATTTGCATGCATGACTGTGTGTGTGTATGTGTGTTTAAACTAAA
Abbassa_amh_5'UTR             29 ------------------------------------------------------------
Abbassa_amhy_h1tg000212l   17090 ------------------------------------------------------------

                                 ....:19270....:19280....:19290....:19300....:19310....:19320
Abbassa_amh_h2tg000119l    17090 ------------------------------------------------------------
Abbassa_amhΔy_h1tg000212l  16508 CGCAGTAAATTTTCACATATCAGTCCACTTTCATTTAAACCTGCATTTCCTAACAGCACG
Abbassa_amh_5'UTR             29 ------------------------------------------------------------
Abbassa_amhy_h1tg000212l   17090 ------------------------------------------------------------

                                 ....:19330....:19340....:19350....:19360....:19370....:19380
Abbassa_amh_h2tg000119l    17090 ------------------------------------------------------------
Abbassa_amhΔy_h1tg000212l  16568 GATAAATATGAATGCATGGATCTCTGCTGTTATGCTTAGCTTACAAAGAGGAATATTTCT
Abbassa_amh_5'UTR             29 ------------------------------------------------------------
Abbassa_amhy_h1tg000212l   17090 ------------------------------------------------------------

                                 ....:19390....:19400....:19410....:19420....:19430....:19440
Abbassa_amh_h2tg000119l    17090 ------------------------------------------------------------
Abbassa_amhΔy_h1tg000212l  16628 GCTGCCGTTTGTGTGACTTCACAAACCGACGGGAAAGAAACGGTTAATAAATACACGCAA
Abbassa_amh_5'UTR             29 ------------------------------------------------------------
Abbassa_amhy_h1tg000212l   17090 ------------------------------------------------------------

                                 ....:19450....:19460....:19470....:19480....:19490....:19500
Abbassa_amh_h2tg000119l    17090 ------------------------------------------------------------
Abbassa_amhΔy_h1tg000212l  16688 ACTCACTGAACTCTGTTGTTCTTGGACATGTGGATTCAGCTGTTCGCTGAACCTTAACAG
Abbassa_amh_5'UTR             29 ------------------------------------------------------------
Abbassa_amhy_h1tg000212l   17090 ------------------------------------------------------------

                                 ..
Abbassa_amh_h2tg000119l    17090 --
Abbassa_amhΔy_h1tg000212l  16748 AG
Abbassa_amh_5'UTR             29 --
Abbassa_amhy_h1tg000212l   17090 --
```
