## supplementary figure 2 for "High-quality genome assembly for the genetically improved Abbassa Nile tilapia enables the reconstruction of X and Y haplotypes"

**Supplementary Figure 2** – Alignment of GIFT *amh*, *amhy*, and *amhΔy* open reading frame (ORF) and their encoded amino acid sequences. Coding sequence conservation is shaded and encoded amino acids are marked as alternating bold to demarcate exon boundaries. Missense single nucleotide polymorphisms contributing to amino acid change is marked in blue on the encoded amino acid sequence.

```
                   ....:...10....:...20....:...30....:...40....:...50....:...60
GIFT_amh        1  ATGTTGGGTCTGCTCGTTCTTTACAGCGAGGCGCTGACACTCTGCTGGACCCTGCAACCG
GIFT_amhy       1  ATGTTGGGTCTGCTCGTTCTTTACAGCGAGGCGCTGACACTCTGCTGGACCCTGCAACCG
GIFT_amhΔy      1  ATGTTGGGTCTGCTCGTTCTTTACAGCGAGGCGCTGACACTCTGCTGGACCCTGCAACCG
                   _M__L__G__L__L__V__L__Y__S__E__A__L__T__L__C__W__T__L__Q__P_
                   _M__L__G__L__L__V__L__Y__S__E__A__L__T__L__C__W__T__L__Q__P_
                   _M__L__G__L__L__V__L__Y__S__E__A__L__T__L__C__W__T__L__Q__P_

                   ....:...70....:...80....:...90....:..100....:..110....:..120
GIFT_amh       61  GCCCAGGACCCCACAGTAACCGAGTACTCACTCCCATCAGCGAAGACCCCATCATCACCA
GIFT_amhy      61  GCCCAGGACCCCACAGTAACCGAGTACTCACTCCCATCAGCGAAGACCCCATCATCACCA
GIFT_amhΔy     61  GCCCAGGACCCCACAGTAACCGAGTACTCACTCCCATCAGCGAAGACCCCATCATCACCA
                   _A__Q__D__P__T__V__T__E__Y__S__L__P__S__A__K__T__P__S__S__P_
                   _A__Q__D__P__T__V__T__E__Y__S__L__P__S__A__K__T__P__S__S__P_
                   _A__Q__D__P__T__V__T__E__Y__S__L__P__S__A__K__T__P__S__S__P_

                   ....:..130....:..140....:..150....:..160....:..170....:..180
GIFT_amh      121  TCATCATCCTCAGCAGCAGCGCCTCATGCTGCACCATGCTTCGTGGAGGACATCTTTGCA
GIFT_amhy     121  TCATCATCCTCAGCAGCAGCGCCTCATGCTGCACCATGCTTCGTGGAGGACATCTTTGCA
GIFT_amhΔy    121  TCATCATCCTCAGCAGCAGCGCCTCATGCTGCACCATGCTTCGTGGAGGACATCTTTGCA
                   _S__S__S__S__A__A__A__P__H__A__A__P__C__F__V__E__D__I__F__A_
                   _S__S__S__S__A__A__A__P__H__A__A__P__C__F__V__E__D__I__F__A_
                   _S__S__S__S__A__A__A__P__H__A__A__P__C__F__V__E__D__I__F__A_

                   ....:..190....:..200....:..210....:..220....:..230....:..240
GIFT_amh      181  GCGTTGCGTGATGGTGTGGGGGACAGCGGCGAACTGACAAACAGCAGTTTGGTTCTGTTT
GIFT_amhy     181  GCGTTGCGTGATGGTGTGGGGGACAGCGGCGAACTGACAAACAGCAGTTTGGTTCTGTTT
GIFT_amhΔy    181  GCGTTGCGTGAGGGTGTGGGGGACAGCGGCGAACTGACAAACAGCAGTTTGGTTCTGTTT
                   _A__L__R__D__G__V__G__D__S__G__E__L__T__N__S__S__L__V__L__F_
                   _A__L__R__D__G__V__G__D__S__G__E__L__T__N__S__S__L__V__L__F_
                   _A__L__R__E__G__V__G__D__S__G__E__L__T__N__S__S__L__V__L__F_

                   ....:..250....:..260....:..270....:..280....:..290....:..300
GIFT_amh      241  GGATTCTGCTCGCAGTCTGCCCGCTCATCAGCCTCGGTCTCGTTAGACCTCGCTAACAAG
GIFT_amhy     241  GGATTCTGCTCGCAGTCTGCCCGCTCATCAGCCTCGGTCTCGTTAGACCTCGCTAACAAG
GIFT_amhΔy    241  GGATTCTGCTCGCAGTCTGCCCGCTCATCAGCCTCGGTCTCGTTAGACCTCGCTAACAAG
                   _G__F__C__S__Q__S__A__R__S__S__A__S__V__S__L__D__L__A__N__K_
                   _G__F__C__S__Q__S__A__R__S__S__A__S__V__S__L__D__L__A__N__K_
                   _G__F__C__S__Q__S__A__R__S__S__A__S__V__S__L__D__L__A__N__K_

                   ....:..310....:..320....:..330....:..340....:..350....:..360
GIFT_amh      301  AAGAGCAGCTTGGAGGTTCTGCACCCAGCTGCAGTACACGTATCAGAGGAAGAGGAGCAA
GIFT_amhy     301  AAGAGCAGCTTGGAGGTTCTGCACCCAGCTGCAGTACACGTATCAGAGGAAGAGGAGCAA
GIFT_amhΔy    301  AAGAGCAGCTTGGAGGTTCTGCACCCAGCTGCAGTACACGTATCAGAGGAAGAGGAGCAA
                   _K__S__S__L__E__V__L__H__P__A__A__V__H__V__S__E__E__E__E__Q_
                   _K__S__S__L__E__V__L__H__P__A__A__V__H__V__S__E__E__E__E__Q_
                   _K__S__S__L__E__V__L__H__P__A__A__V__H__V__S__E__E__E__E__Q_

                   ....:..370....:..380....:..390....:..400....:..410....:..420
GIFT_amh      361  GGAACAATCACGTTGACCTTTGACCTCCCACGGCCTCCATCGCTCATGACAAACCCTGTG
GIFT_amhy     361  GGAACAATCACGTTGACCTTTGACCTCCCACGGCCTCCATCGCTCATGACAAACCCTGTG
GIFT_amhΔy    361  GGAACAATCACGTTGACCTTTGACCTCCCACGGCCTCCTTCGCTCATGACAAACCCTGTG
                   _G__T__I__T__L__T__F__D__L__P__R__P__P__S__L__M__T__N__P__V_
                   _G__T__I__T__L__T__F__D__L__P__R__P__P__S__L__M__T__N__P__V_
                   _G__T__I__T__L__T__F__D__L__P__R__P__P__S__L__M__T__N__P__V_

                   ....:..430....:..440....:..450....:..460....:..470....:..480
GIFT_amh      421  CTGCTCTTGGTCTTTGAAAATCCACTGGCACGAGGAGACCTGGAAGTTGCTTTCACTAGT
GIFT_amhy     421  CTGCTCTTGGTCTTTGAAAATCCACTGGCACGAGGAGACCTGGAAGTTGCTTTCACTAGT
GIFT_amhΔy    421  CTGCTCTTGGTCTTTGAAAGTCCACTGGCACGAGGAGACCTGGAAGTTGCTTTCACTAGT
                   _L__L__L__V__F__E__N__P__L__A__R__G__D__L__E__V__A__F__T__S_
                   _L__L__L__V__F__E__N__P__L__A__R__G__D__L__E__V__A__F__T__S_
                   _L__L__L__V__F__E__S__P__L__A__R__G__D__L__E__V__A__F__T__S_

                   ....:..490....:..500....:..510....:..520....:..530....:..540
GIFT_amh      481  CAGTTTCTGCAGCCTAACACGCAGGCTGTGTGCATTTCAGGAGACACACAGTACGTACTG
```

```
GIFT_amhy      481 CAGTTTCTGCAGCCTAACACGCAGGCTGTGTGCATTTCAGGAGACACACAGTACGTACTG
GIFT_amhΔy     481 CAGTTTCTGCAGCCTAACACGCAGGCTGTGTGCATTTCAGGAGACACACAGTACGTACTG
                   _Q__F__L__Q__P__N__T__Q__A__V__C__I__S__G__D__T__Q__Y__V__L_
                   _Q__F__L__Q__P__N__T__Q__A__V__C__I__S__G__D__T__Q__Y__V__L_
                   _Q__F__L__Q__P__N__T__Q__A__V__C__I__S__G__D__T__Q__Y__V__L_

                   ....:..550....:..560....:..570....:..580....:..590....:..600
GIFT_amh       541 CTGACAGGAAAATCATCAGAGGGGAGTGTTAATGACAGGTGGCAGATTACGGCTCAGACA
GIFT_amhy      541 CTGACAGGAAAATCATCAGAGGGGAGTGTTAATGACAGGTGGCAGATTACGGCTCAGACA
GIFT_amhΔy     541 CTGACAGGAAAATCATCAGAGGGGAGTGTTAATGACAGGTGGCAGATTACGGCTCAGACA
                   _L__T__G__K__S__S__E__G__S__V__N__D__R__W__Q__I__T__A__Q__T_
                   _L__T__G__K__S__S__E__G__S__V__N__D__R__W__Q__I__T__A__Q__T_
                   _L__T__G__K__S__S__E__G__S__V__N__D__R__W__Q__I__T__A__Q__T_

                   ....:..610....:..620....:..630....:..640....:..650....:..660
GIFT_amh       601 AAACTCCCTCATATGAAGCAAAACCTAAAAAGCATCTTGATTGGTGAAAAATCAGGAAGT
GIFT_amhy      601 AAACTCCCTCATATGAAGCAAAACCTAAAAAGCATCTTGATTGGTGAAAAATCAGGAAGT
GIFT_amhΔy     601 AAACTCCCTCATATGAAGCAAAACCTAAAAAGCATCTTGATTGGTGAAAAATCAGGAAGT
                   _K__L__P__H__M__K__Q__N__L__K__S__I__L__I__G__E__K__S__G__S_
                   _K__L__P__H__M__K__Q__N__L__K__S__I__L__I__G__E__K__S__G__S_
                   _K__L__P__H__M__K__Q__N__L__K__S__I__L__I__G__E__K__S__G__S_

                   ....:..670....:..680....:..690....:..700....:..710....:..720
GIFT_amh       661 AACATCAGCATGAGTCCACTTCTACTTTTCTCCGGGGGAACGGGAACTGATACGAGATGT
GIFT_amhy      661 AACATCAGCATGAGTCCACTTCTACTTTTCTCCGGGGGAACGGGAACTGATACGAGATGT
GIFT_amhΔy     661 AACATCAGCATGAGTCCACTTCTACTTTTCTCCGGGGGAACGGGAACTGATACGAGATGT
                   _N__I__S__M__S__P__L__L__L__F__S__G__G__T__G__T__D__T__R__C_
                   _N__I__S__M__S__P__L__L__L__F__S__G__G__T__G__T__D__T__R__C_
                   _N__I__S__M__S__P__L__L__L__F__S__G__G__T__G__T__D__T__R__C_

                   ....:..730....:..740....:..750....:..760....:..770....:..780
GIFT_amh       721 GCTTCAGGCTCGCCCCCGGCATCTCTGCAAACCTCCTTCCTTTGTGAGATGAAACGCTTC
GIFT_amhy      721 GCTTCAGGCTCGCCCCCGGCATCTCTGCAAACCTCCTTCCTTTGTGAGATGAAACGCTTC
GIFT_amhΔy     721 GCTTCAGGCTCGCCCCCGGCATCTCTGCAAACCTCCTTCCTTTGTGAATGTCGATGA---
                   _A__S__G__S__P__P__A__S__L__Q__T__S__F__L__C__E__M__K__R__F_
                   _A__S__G__S__P__P__A__S__L__Q__T__S__F__L__C__E__M__K__R__F_
                   _A__S__G__S__P__P__A__S__L__Q__T__S__F__L__C__E__C__R__*__-_

                   ....:..790....:..800....:..810....:..820....:..830....:..840
GIFT_amh       781 CTGGGTGCTGTTCTCCCTCAGGAACACTTCACGTCCCCTCCACTTCCTCTGGACTCCTTA
GIFT_amhy      781 CTGGGTGCTGTTCTCCCTCAGGAACACTTCACGTCCCCTCCACTTCCTCTGGACTCCTTA
GIFT_amhΔy     768 ------------------------------------------------------------
                   _L__G__A__V__L__P__Q__E__H__F__T__S__P__P__L__P__L__D__S__L_
                   _L__G__A__V__L__P__Q__E__H__F__T__S__P__P__L__P__L__D__S__L_
                   _-__-__-__-__-__-__-__-__-__-__-__-__-__-__-__-__-__-__-__-_

                   ....:..850....:..860....:..870....:..880....:..890....:..900
GIFT_amh       841 CAGTCTCTGCCTCCCCTCTCGCTTGGCTTATCCTCCAGCGAGACCCTGCTGGCAGTAATG
GIFT_amhy      841 CAGTCTCTGCCTCCCCTCTCGCTTGGCTTATCCTCCAGCGAGACCCTGCTGGCAGTAATG
GIFT_amhΔy     768 ------------------------------------------------------------
                   _Q__S__L__P__P__L__S__L__G__L__S__S__S__E__T__L__L__A__V__M_
                   _Q__S__L__P__P__L__S__L__G__L__S__S__S__E__T__L__L__A__V__M_
                   _-__-__-__-__-__-__-__-__-__-__-__-__-__-__-__-__-__-__-__-_

                   ....:..910....:..920....:..930....:..940....:..950....:..960
GIFT_amh       901 ATCAACTCCACAGCTCCCACAGTCTTTGGCTTCACGAGCTGGGGCTCCGTGTTGCCGGTG
GIFT_amhy      901 ATCAACTCCACAGCTCCCACAGTCTTTGGCTTCACGAGCTGGGGCTCCGTGTTGCCGGTG
GIFT_amhΔy     768 ------------------------------------------------------------
                   _I__N__S__T__A__P__T__V__F__G__F__T__S__W__G__S__V__L__P__V_
                   _I__N__S__T__A__P__T__V__F__G__F__T__S__W__G__S__V__L__P__V_
                   _-__-__-__-__-__-__-__-__-__-__-__-__-__-__-__-__-__-__-__-_

                   ....:..970....:..980....:..990....:.1000....:.1010....:.1020
GIFT_amh       961 TGCCACGGAGAGCTGGCCCTGTCTGCTGCACTGTTAGAGGAGCTCAGACAGAGACTGGAC
GIFT_amhy      961 TGCCACGGAGAGCTGGCCCTGTCTGCTGCACTGTTAGAGGAGCTCAGACAGAGACTGGAC
GIFT_amhΔy     768 ------------------------------------------------------------
```

```
                    _C__H__G__E__L__A__L__S__A__A__L__L__E__E__L__R__Q__R__L__D_
                    _C__H__G__E__L__A__L__S__A__A__L__L__E__E__L__R__Q__R__L__D_
                    _-__-__-__-__-__-__-__-__-__-__-__-__-__-__-__-__-__-__-__-_

                    ....:.1030....:.1040....:.1050....:.1060....:.1070....:.1080
GIFT_amh       1021 CAGACTTTGGTGCAAATGACAGAAATAATCAGAGAGGAAGAGGTTTCACCGGGAGCCAAG
GIFT_amhy      1021 CAGACTTTGGTGCAAATGACAGAAATAATCAGAGAGGAAGAGGTTTCACCGGGAGCCAAG
GIFT_amhΔy      768 ------------------------------------------------------------
                    _Q__T__L__V__Q__M__T__E__I__I__R__E__E__E__V__S__P__G__A__K_
                    _Q__T__L__V__Q__M__T__E__I__I__R__E__E__E__V__S__P__G__A__K_
                    _-__-__-__-__-__-__-__-__-__-__-__-__-__-__-__-__-__-__-__-_

                    ....:.1090....:.1100....:.1110....:.1120....:.1130....:.1140
GIFT_amh       1081 GAGAGCCTGGGGAGGCTCAAAGAACTGAGTGCGTTACAGGAGAAAGAACATGCCACAGGA
GIFT_amhy      1081 GAGAGCCTGGGGAGGCTCAAAGAACTGAGTGCGTTACAGGAGAAAGAACATGCCACAGGA
GIFT_amhΔy      768 ------------------------------------------------------------
                    _E__S__L__G__R__L__K__E__L__S__A__L__Q__E__K__E__H__A__T__G_
                    _E__S__L__G__R__L__K__E__L__S__A__L__Q__E__K__E__H__A__T__G_
                    _-__-__-__-__-__-__-__-__-__-__-__-__-__-__-__-__-__-__-__-_

                    ....:.1150....:.1160....:.1170....:.1180....:.1190....:.1200
GIFT_amh       1141 GGGAGTCAGTTCCGTGCGTTTCTTCTGCTGAAGGCTCTGCAGACGGTGGCCCAAACGTAC
GIFT_amhy      1141 GGGAGTCAGTTCCGTGTGTTTCTTCTGCTGAAGGCTCTGCAGACGGTGGCCCAAACGTAC
GIFT_amhΔy      768 ------------------------------------------------------------
                    _G__S__Q__F__R__A__F__L__L__L__K__A__L__Q__T__V__A__Q__T__Y_
                    _G__S__Q__F__R__V__F__L__L__L__K__A__L__Q__T__V__A__Q__T__Y_
                    _-__-__-__-__-__-__-__-__-__-__-__-__-__-__-__-__-__-__-__-_

                    ....:.1210....:.1220....:.1230....:.1240....:.1250....:.1260
GIFT_amh       1201 GACGCGCAAAGAAAACTGCGGGCCACCAGAGCAGACCCCAGTTCGTCAGTGAGGGGCGGC
GIFT_amhy      1201 GACGCGCAAAGAAAACTGCGGGCCACCAGAGCAGACCCCAGTTCGTCAGTGAGGGGCGGC
GIFT_amhΔy      768 ------------------------------------------------------------
                    _D__A__Q__R__K__L__R__A__T__R__A__D__P__S__S__S__V__R__G__G_
                    _D__A__Q__R__K__L__R__A__T__R__A__D__P__S__S__S__V__R__G__G_
                    _-__-__-__-__-__-__-__-__-__-__-__-__-__-__-__-__-__-__-__-_

                    ....:.1270....:.1280....:.1290....:.1300....:.1310....:.1320
GIFT_amh       1261 GTCTGTGGGCTGAAGGCTCTCACCGTGTCCCTGACAAAGCTTCTTGTCGGCCCAAGCAGC
GIFT_amhy      1261 GTCTGTGGGCTGAAGGCTCTCACCGTGTCCCTGACAAAGCTTCTTGTCGGCCCAAGCAGC
GIFT_amhΔy      768 ------------------------------------------------------------
                    _V__C__G__L__K__A__L__T__V__S__L__T__K__L__L__V__G__P__S__S_
                    _V__C__G__L__K__A__L__T__V__S__L__T__K__L__L__V__G__P__S__S_
                    _-__-__-__-__-__-__-__-__-__-__-__-__-__-__-__-__-__-__-__-_

                    ....:.1330....:.1340....:.1350....:.1360....:.1370....:.1380
GIFT_amh       1321 GCAAACATTAACAATTGCCACGGCTCCTGCGCGTTCCCTCTGACCAACGGCAACAACCAC
GIFT_amhy      1321 GCAAACATTAACAATTGCCACGGCTCCTGCGCGTTCCCTCTGACCAACGGCAACAACCAC
GIFT_amhΔy      768 ------------------------------------------------------------
                    _A__N__I__N__N__C__H__G__S__C__A__F__P__L__T__N__G__N__N__H_
                    _A__N__I__N__N__C__H__G__S__C__A__F__P__L__T__N__G__N__N__H_
                    _-__-__-__-__-__-__-__-__-__-__-__-__-__-__-__-__-__-__-__-_

                    ....:.1390....:.1400....:.1410....:.1420....:.1430....:.1440
GIFT_amh       1381 GCCATCCTGCTCAACTCCCACATCGAGACCGGCAACGCGGATGAGCGTTCGCCCTGCTGT
GIFT_amhy      1381 GCCATCCTGCTCAACTCCCACATCGAGACCGGCAACGCGGATGAGCGTTCGCCCTGCTGT
GIFT_amhΔy      768 ------------------------------------------------------------
                    _A__I__L__L__N__S__H__I__E__T__G__N__A__D__E__R__S__P__C__C_
                    _A__I__L__L__N__S__H__I__E__T__G__N__A__D__E__R__S__P__C__C_
                    _-__-__-__-__-__-__-__-__-__-__-__-__-__-__-__-__-__-__-__-_
```

```
                    ....:.1450....:.1460....:.1470....:.1480....:.1490....:.1500
GIFT_amh      1441 GTGCCCGTGGCATACGAAGCCCTGGAGGTTGTGGACTGGAACGCAGATGGGACCTTCATC
GIFT_amhy     1441 GTGCCCGTGGCATACGAAGCCCTGGAGGTTGTGGACTGGAACGCAGATGGGACCTTCATC
GIFT_amhΔy     768 ------------------------------------------------------------
                   _V__P__V__A__Y__E__A__L__E__V__V__D__W__N__A__D__G__T__F__I_
                   _V__P__V__A__Y__E__A__L__E__V__V__D__W__N__A__D__G__T__F__I_
                   _-__-__-__-__-__-__-__-__-__-__-__-__-__-__-__-__-__-__-__-_

                   ....:.1510....:.1520....:.1530....:.1540....:
GIFT_amh      1501 TCCATCAAGCCAGATGCGGTTGCGAGGGAGTGTGGATGCCGCTAG
GIFT_amhy     1501 TCCATCAAGCCAGATGCGGTTGCGAGGGAGTGTGGATGCCGCTAG
GIFT_amhΔy     768 ---------------------------------------------
                   _S__I__K__P__D__A__V__A__R__E__C__G__C__R__*_
                   _S__I__K__P__D__A__V__A__R__E__C__G__C__R__*_
                   _-__-__-__-__-__-__-__-__-__-__-__-__-__-__-_
```
