## supplementary figure 1 for "High-quality genome assembly for the genetically improved Abbassa Nile tilapia enables the reconstruction of X and Y haplotypes"

**Supplementary Figure 1** – Alignment of Abbassa *amh, amhy,* and *amhΔy* open reading frame (ORF) and their encoded amino acid sequences. Coding sequence conservation is shaded and encoded amino acids are marked as alternating bold to demarcate exon boundaries. Missense single nucleotide polymorphisms contributing to amino acid change is marked in blue on the encoded amino acid sequence.

```
                          ....:...10....:...20....:...30....:...40....:...50....:...60
Abbassa_amh          1    ATGTTGGGTCTGCTCGTTCTTTACAGCGAGGCGCTGACACTCTGCTGGACCCTGCAACCG
Abbassa_amhy         1    ATGTTGGGTCTGCTCGTTCTTTACAGCGAGGCGCTGACACTCTGCTGGACCCTGCAACCG
Abbassa_amhΔy        1    ATGTTGGGTCTGCTCGTTCTTTACAGCGAGGCGCTGACACTCTGCTGGACCCTGCAACCG
                          _M__L__G__L__L__V__L__Y__S__E__A__L__T__L__C__W__T__L__Q__P_
                          _M__L__G__L__L__V__L__Y__S__E__A__L__T__L__C__W__T__L__Q__P_
                          _M__L__G__L__L__V__L__Y__S__E__A__L__T__L__C__W__T__L__Q__P_

                          ....:...70....:...80....:...90....:..100....:..110....:..120
Abbassa_amh          61   GCCCAGGACCCCACAGTAACCGAGTACTCACTCCCATCAGCGAAGACCCCATCATCACCA
Abbassa_amhy         61   GCCCAGGACCCCACAGTAACCGAGTACTCACTCCCATCAGCGAAGACCCCATCATCACCA
Abbassa_amhΔy        61   GCCCAGGACCCCACAGTAACCGAGTACTCACTCCCATCAGCGAAGACCCCATCATCACCA
                          _A__Q__D__P__T__V__T__E__Y__S__L__P__S__A__K__T__P__S__S__P_
                          _A__Q__D__P__T__V__T__E__Y__S__L__P__S__A__K__T__P__S__S__P_
                          _A__Q__D__P__T__V__T__E__Y__S__L__P__S__A__K__T__P__S__S__P_

                          ....:..130....:..140....:..150....:..160....:..170....:..180
Abbassa_amh          121  TCATCATCCTCAGCAGCAGCGCCTCATGCTGCACCATGCTTCGTGGAGGACATCTTTGCA
Abbassa_amhy         121  TCATCATCCTCAGCAGCAGCGCCTCATGCTGCACCATGCTTCGTGGAGGACATCTTTGCA
Abbassa_amhΔy        121  TCATCATCCTCAGCAGCAGCGCCTCATGCTGCACCATGCTTCGTGGAGGACATCTTTGCA
                          _S__S__S__S__A__A__A__P__H__A__A__P__C__F__V__E__D__I__F__A_
                          _S__S__S__S__A__A__A__P__H__A__A__P__C__F__V__E__D__I__F__A_
                          _S__S__S__S__A__A__A__P__H__A__A__P__C__F__V__E__D__I__F__A_

                          ....:..190....:..200....:..210....:..220....:..230....:..240
Abbassa_amh          181  GCGTTGCGTGATGGTGTGGGGGACAGCGGCGAACTGACAAACAGCAGTTTGGTTCTGTTT
Abbassa_amhy         181  GCGTTGCGTGATGGTGTGGGGGACAGCGGCGAACTGACAAACAGCAGTTTGGTTCTGTTT
Abbassa_amhΔy        181  GCGTTGCGTGAGGGTGTGGGGGACAGCGGCGAACTGACAAACAGCAGTTTGGTTCTGTTT
                          _A__L__R__D__G__V__G__D__S__G__E__L__T__N__S__S__L__V__L__F_
                          _A__L__R__D__G__V__G__D__S__G__E__L__T__N__S__S__L__V__L__F_
                          _A__L__R__E__G__V__G__D__S__G__E__L__T__N__S__S__L__V__L__F_

                          ....:..250....:..260....:..270....:..280....:..290....:..300
Abbassa_amh          241  GGATTCTGCTCGCAGTCTGCCCGCTCATCAGCCTCGGTCTCGTTAGACCTCGCTAACAAG
Abbassa_amhy         241  GGATTCTGCTCGCAGTCTGCCCGCTCATCAGCCTCGGTCTCGTTAGACCTCGCTAACAAG
Abbassa_amhΔy        241  GGATTCTGCTCGCAGTCTGCCCGCTCATCAGCCTCGGTCTCGTTAGACCTCGCTAACAAG
                          _G__F__C__S__Q__S__A__R__S__S__A__S__V__S__L__D__L__A__N__K_
                          _G__F__C__S__Q__S__A__R__S__S__A__S__V__S__L__D__L__A__N__K_
                          _G__F__C__S__Q__S__A__R__S__S__A__S__V__S__L__D__L__A__N__K_

                          ....:..310....:..320....:..330....:..340....:..350....:..360
Abbassa_amh          301  AAGAGCAGCTTGGAGGTTCTGCACCCAGCTGCAGTACACGTATCAGAGGAAGAGGAGCAA
Abbassa_amhy         301  AAGAGCAGCTTGGAGGTTCTGCACCCAGCTGCAGTACACGTATCAGAGGAAGAGGAGCAA
Abbassa_amhΔy        301  AAGAGCAGCTTGGAGGTTCTGCACCCAGCTGCAGTACACGTATCAGAGGAAGAGGAGCAA
                          _K__S__S__L__E__V__L__H__P__A__A__V__H__V__S__E__E__E__E__Q_
                          _K__S__S__L__E__V__L__H__P__A__A__V__H__V__S__E__E__E__E__Q_
                          _K__S__S__L__E__V__L__H__P__A__A__V__H__V__S__E__E__E__E__Q_

                          ....:..370....:..380....:..390....:..400....:..410....:..420
Abbassa_amh          361  GGAACAATCACGTTGACCTTTGACCTCCCACGGCCTCCATCGCTCATGACAAACCCTGTG
Abbassa_amhy         361  GGAACAATCACGTTGACCTTTGACCTCCCACGGCCTCCATCGCTCATGACAAACCCTGTG
Abbassa_amhΔy        361  GGAACAATCACGTTGACCTTTGACCTCCCACGGCCTCCTTCGCTCATGACAAACCCTGTG
                          _G__T__I__T__L__T__F__D__L__P__R__P__P__S__L__M__T__N__P__V_
                          _G__T__I__T__L__T__F__D__L__P__R__P__P__S__L__M__T__N__P__V_
                          _G__T__I__T__L__T__F__D__L__P__R__P__P__S__L__M__T__N__P__V_

                          ....:..430....:..440....:..450....:..460....:..470....:..480
Abbassa_amh          421  CTGCTCTTGGTCTTTGAAAATCCACTGGCACGAGGAGACCTGGAAGTTGCTTTCACTAGT
Abbassa_amhy         421  CTGCTCTTGGTCTTTGAAAATCCACTGGCACGAGGAGACCTGGAAGTTGCTTTCACTAGT
Abbassa_amhΔy        421  CTGCTCTTGGTCTTTGAAAGTCCACTGGCACGAGGAGACCTGGAAGTTGCTTTCACTAGT
                          _L__L__L__V__F__E__N__P__L__A__R__G__D__L__E__V__A__F__T__S_
                          _L__L__L__V__F__E__N__P__L__A__R__G__D__L__E__V__A__F__T__S_
                          _L__L__L__V__F__E__S__P__L__A__R__G__D__L__E__V__A__F__T__S_

                          ....:..490....:..500....:..510....:..520....:..530....:..540
Abbassa_amh          481  CAGTTTCTGCAGCCTAACACGCAGGCTGTGTGCATTTCAGGAGACACACAGTACGTACTG
```

```
Abbassa_amhy      481 CAGTTTCTGCAGCCTAACACGCAGGCTGTGTGCATTTCAGGAGACACACAGTACGTACTG
Abbassa_amhΔy     481 CAGTTTCTGCAGCCTAACACGCAGGCTGTGTGCATTTCAGGAGACACACAGTACGTACTG
                      _Q__F__L__Q__P__N__T__Q__A__V__C__I__S__G__D__T__Q__Y__V__L_
                      _Q__F__L__Q__P__N__T__Q__A__V__C__I__S__G__D__T__Q__Y__V__L_
                      _Q__F__L__Q__P__N__T__Q__A__V__C__I__S__G__D__T__Q__Y__V__L_

                      ....:..550....:..560....:..570....:..580....:..590....:..600
Abbassa_amh       541 CTGACAGGAAAATCATCAGAGGGGAGTGTTAATGACAGGTGGCAGATTACGGCTCAGACA
Abbassa_amhy      541 CTGACAGGAAAATCATCAGAGGGGAGTGTTAATGACAGGTGGCAGATTACGGCTCAGACA
Abbassa_amhΔy     541 CTGACAGGAAAATCATCAGAGGGGAGTGTTAATGACAGGTGGCAGATTACGGCTCAGACA
                      _L__T__G__K__S__S__E__G__S__V__N__D__R__W__Q__I__T__A__Q__T_
                      _L__T__G__K__S__S__E__G__S__V__N__D__R__W__Q__I__T__A__Q__T_
                      _L__T__G__K__S__S__E__G__S__V__N__D__R__W__Q__I__T__A__Q__T_

                      ....:..610....:..620....:..630....:..640....:..650....:..660
Abbassa_amh       601 AAACTCCCTCATATGAAGCAAAACCTAAAAAGCATCTTGATTGGTGAAAAATCAGGAAGT
Abbassa_amhy      601 AAACTCCCTCATATGAAGCAAAACCTAAAAAGCATCTTGATTGGTGAAAAATCAGGAAGT
Abbassa_amhΔy     601 AAACTCCCTCATATGAAGCAAAACCTAAAAAGCATCTTGATTGGTGAAAAATCAGGAAGT
                      _K__L__P__H__M__K__Q__N__L__K__S__I__L__I__G__E__K__S__G__S_
                      _K__L__P__H__M__K__Q__N__L__K__S__I__L__I__G__E__K__S__G__S_
                      _K__L__P__H__M__K__Q__N__L__K__S__I__L__I__G__E__K__S__G__S_

                      ....:..670....:..680....:..690....:..700....:..710....:..720
Abbassa_amh       661 AACATCAGCATGAGTCCACTTCTACTTTTCTCCGGCGGAACGGGAACTGATACGAGATGT
Abbassa_amhy      661 AACATCAGCATGAGTCCACTTCTACTTTTCTCCGGCGGAACGGGAACTGATACGAGATGT
Abbassa_amhΔy     661 AACATCAGCATGAGTCCACTTCTACTTTTCTCCGGGGGAACGGGAACTGATACGAGATGT
                      _N__I__S__M__S__P__L__L__L__F__S__G__G__T__G__T__D__T__R__C_
                      _N__I__S__M__S__P__L__L__L__F__S__G__G__T__G__T__D__T__R__C_
                      _N__I__S__M__S__P__L__L__L__F__S__G__G__T__G__T__D__T__R__C_

                      ....:..730....:..740....:..750....:..760....:..770....:..780
Abbassa_amh       721 GCTTCAGGCTCGCCCCCGGCATCTCTGCAAACCTCCTTCCTTTGTGAGATGAAACGCTTC
Abbassa_amhy      721 GCTTCAGGCTCGCCCCCGGCATCTCTGCAAACCTCCTTCCTTTGTGAGATGAAACGCTTC
Abbassa_amhΔy     721 GCTTCAGGCTCGCCCCCGGCATCTCTGCAAACCTCCTTCCTTTGTGAATGTCGATGA---
                      _A__S__G__S__P__P__A__S__L__Q__T__S__F__L__C__E__M__K__R__F_
                      _A__S__G__S__P__P__A__S__L__Q__T__S__F__L__C__E__M__K__R__F_
                      _A__S__G__S__P__P__A__S__L__Q__T__S__F__L__C__E__C__R__*__-_

                      ....:..790....:..800....:..810....:..820....:..830....:..840
Abbassa_amh       781 CTGGGTGCTGTTCTCCCTCAGGAACACTTCACGTCCCCTCCACTTCCTCTGGACTCCTTA
Abbassa_amhy      781 CTGGGTGCTGTTCTCCCTCAGGAACACTTCACGTCCCCTCCACTTCCTCTGGACTCCTTA
Abbassa_amhΔy     768 ------------------------------------------------------------
                      _L__G__A__V__L__P__Q__E__H__F__T__S__P__P__L__P__L__D__S__L_
                      _L__G__A__V__L__P__Q__E__H__F__T__S__P__P__L__P__L__D__S__L_
                      _-__-__-__-__-__-__-__-__-__-__-__-__-__-__-__-__-__-__-__-_

                      ....:..850....:..860....:..870....:..880....:..890....:..900
Abbassa_amh       841 CAGTCTCTGCCTCCCCTCTCGCTTGGCTTATCCTCCAGCGAGACCCTGCTGGCAGTAATG
Abbassa_amhy      841 CAGTCTCTGCCTCCCCTCTCGCTTGGCTTATCCTCCAGCGAGACCCTGCTGGCAGTAATG
Abbassa_amhΔy     768 ------------------------------------------------------------
                      _Q__S__L__P__P__L__S__L__G__L__S__S__S__E__T__L__L__A__V__M_
                      _Q__S__L__P__P__L__S__L__G__L__S__S__S__E__T__L__L__A__V__M_
                      _-__-__-__-__-__-__-__-__-__-__-__-__-__-__-__-__-__-__-__-_

                      ....:..910....:..920....:..930....:..940....:..950....:..960
Abbassa_amh       901 ATCAACTCCACAGCTCCCACAGTCTTTGGCTTCACGAGCTGGGGCTCCGTGTTGCCGGTG
Abbassa_amhy      901 ATCAACTCCACAGCTCCCACAGTCTTTGGCTTCACGAGCTGGGGCTCCGTGTTGCCGGTG
Abbassa_amhΔy     768 ------------------------------------------------------------
                      _I__N__S__T__A__P__T__V__F__G__F__T__S__W__G__S__V__L__P__V_
                      _I__N__S__T__A__P__T__V__F__G__F__T__S__W__G__S__V__L__P__V_
                      _-__-__-__-__-__-__-__-__-__-__-__-__-__-__-__-__-__-__-__-_

                      ....:..970....:..980....:..990....:.1000....:.1010....:.1020
Abbassa_amh       961 TGCCACGGAGAGCTGGCCCTGTCTGCTGCACTGTTAGAGGAGCTCAGACAGAGACTGGAC
Abbassa_amhy      961 TGCCACGGAGAGCTGGCCCTGTCTGCTGCACTGTTAGAGGAGCTCAGACAGAGACTGGAC
Abbassa_amhΔy     768 ------------------------------------------------------------
```

```
                     _C__H__G__E__L__A__L__S__A__A__L__L__E__E__L__R__Q__R__L__D_
                     _C__H__G__E__L__A__L__S__A__A__L__L__E__E__L__R__Q__R__L__D_
                     _-__-__-__-__-__-__-__-__-__-__-__-__-__-__-__-__-__-__-__-_

                     ....:.1030....:.1040....:.1050....:.1060....:.1070....:.1080
Abbassa_amh      1021 CAGACTTTGGTGCAAATGACAGAAATAATCAGAGAGGAAGAGGTTTCACTGGGAGCCAAG
Abbassa_amhy     1021 CAGACTTTGGTGCAAATGACAGAAATAATCAGAGAGGAAGAGGTTTCACTGGGAGCCAAG
Abbassa_amhΔy     768 ------------------------------------------------------------
                     _Q__T__L__V__Q__M__T__E__I__I__R__E__E__E__V__S__L__G__A__K_
                     _Q__T__L__V__Q__M__T__E__I__I__R__E__E__E__V__S__L__G__A__K_
                     _-__-__-__-__-__-__-__-__-__-__-__-__-__-__-__-__-__-__-__-_

                     ....:.1090....:.1100....:.1110....:.1120....:.1130....:.1140
Abbassa_amh      1081 GAGAGCCTGGGGAGGCTCAAAGAACTGAGTGCGTTACAGGAGAAAGAACATGCCACAGGA
Abbassa_amhy     1081 GAGAGCCTGGGGAGGCTCAAAGAACTGAGTGCGTTACAGGAGAAAGAACATGCCACAGGA
Abbassa_amhΔy     768 ------------------------------------------------------------
                     _E__S__L__G__R__L__K__E__L__S__A__L__Q__E__K__E__H__A__T__G_
                     _E__S__L__G__R__L__K__E__L__S__A__L__Q__E__K__E__H__A__T__G_
                     _-__-__-__-__-__-__-__-__-__-__-__-__-__-__-__-__-__-__-__-_

                     ....:.1150....:.1160....:.1170....:.1180....:.1190....:.1200
Abbassa_amh      1141 GGGAGTCAGTTCCGTGTGTTTCTTCTGCTGAAGGCTCTGCAGACGGTGGCCCAAACGTAC
Abbassa_amhy     1141 GGGAGTCAGTTCCGTGTGTTTCTTCTGCTGAAGGCTCTGCAGACGGTGGCCCAAACGTAC
Abbassa_amhΔy     768 ------------------------------------------------------------
                     _G__S__Q__F__R__V__F__L__L__L__K__A__L__Q__T__V__A__Q__T__Y_
                     _G__S__Q__F__R__V__F__L__L__L__K__A__L__Q__T__V__A__Q__T__Y_
                     _-__-__-__-__-__-__-__-__-__-__-__-__-__-__-__-__-__-__-__-_

                     ....:.1210....:.1220....:.1230....:.1240....:.1250....:.1260
Abbassa_amh      1201 GACGCGCAAAGAAAACTGCGGGCCACCAGAGCAGACCCCAGTTCGTCAGTGAGGGGCGGC
Abbassa_amhy     1201 GACGCGCAAAGAAAACTGCGGGCCACCAGAGCAGACCCCAGTTCGTCAGTGAGGGGCGGC
Abbassa_amhΔy     768 ------------------------------------------------------------
                     _D__A__Q__R__K__L__R__A__T__R__A__D__P__S__S__S__V__R__G__G_
                     _D__A__Q__R__K__L__R__A__T__R__A__D__P__S__S__S__V__R__G__G_
                     _-__-__-__-__-__-__-__-__-__-__-__-__-__-__-__-__-__-__-__-_

                     ....:.1270....:.1280....:.1290....:.1300....:.1310....:.1320
Abbassa_amh      1261 GTCTGTGGGCTGAAGGCTCTCACCGTGTCCCTGACAAAGCTTCTTGTCGGCCCAAGCAGC
Abbassa_amhy     1261 GTCTGTGGGCTGAAGGCTCTCACCGTGTCCCTGACAAAGCTTCTTGTCGGCCCAAGCAGC
Abbassa_amhΔy     768 ------------------------------------------------------------
                     _V__C__G__L__K__A__L__T__V__S__L__T__K__L__L__V__G__P__S__S_
                     _V__C__G__L__K__A__L__T__V__S__L__T__K__L__L__V__G__P__S__S_
                     _-__-__-__-__-__-__-__-__-__-__-__-__-__-__-__-__-__-__-__-_

                     ....:.1330....:.1340....:.1350....:.1360....:.1370....:.1380
Abbassa_amh      1321 GCAAACATTAACAATTGCCACGGCTCCTGCACGTTCCCTCTGACCAACGGCAACAACCAC
Abbassa_amhy     1321 GCAAACATTAACAATTGCCACGGCTCCTGCGCGTTCCCTCTGACCAACGGCAACAACCAC
Abbassa_amhΔy     768 ------------------------------------------------------------
                     _A__N__I__N__N__C__H__G__S__C__T__F__P__L__T__N__G__N__N__H_
                     _A__N__I__N__N__C__H__G__S__C__A__F__P__L__T__N__G__N__N__H_
                     _-__-__-__-__-__-__-__-__-__-__-__-__-__-__-__-__-__-__-__-_

                     ....:.1390....:.1400....:.1410....:.1420....:.1430....:.1440
Abbassa_amh      1381 GCCATCCTGCTCAACTCCCACATCGAGACCGGCAACGCGGATGAGCGTTCGCCCTGCTGT
Abbassa_amhy     1381 GCCATCCTGCTCAACTCCCACATCGAGACCGGCAACGCGGATGAGCGTTCGCCCTGCTGT
Abbassa_amhΔy     768 ------------------------------------------------------------
                     _A__I__L__L__N__S__H__I__E__T__G__N__A__D__E__R__S__P__C__C_
                     _A__I__L__L__N__S__H__I__E__T__G__N__A__D__E__R__S__P__C__C_
                     _-__-__-__-__-__-__-__-__-__-__-__-__-__-__-__-__-__-__-__-_

                     ....:.1450....:.1460....:.1470....:.1480....:.1490....:.1500
Abbassa_amh      1441 GTGCCCGTGGCATACGAAGCCCTGGAGGTTGTGGACTGGAACGCAGATGGGACCTTCATC
Abbassa_amhy     1441 GTGCCCGTGGCATACGAAGCCCTGGAGGTTGTGGACTGGAACGCAGATGGGACCTTCATC
Abbassa_amhΔy     768 ------------------------------------------------------------
                     _V__P__V__A__Y__E__A__L__E__V__V__D__W__N__A__D__G__T__F__I_
                     _V__P__V__A__Y__E__A__L__E__V__V__D__W__N__A__D__G__T__F__I_
```

```
                         _-__-__-__-__-__-__-__-__-__-__-__-__-__-__-__-__-__-__-__-_

                         ....:.1510....:.1520....:.1530....:.1540....:
Abbassa_amh        1501  TCCATCAAGCCAGATGCGGTTGCGAGGGAGTGTGGATGCCGCTAG
Abbassa_amhy       1501  TCCATCAAGCCAGATGCGGTTGCGAGGGAGTGTGGATGCCGCTAG
Abbassa_amhΔy       768  ---------------------------------------------
                         _S__I__K__P__D__A__V__A__R__E__C__G__C__R__*_
                         _S__I__K__P__D__A__V__A__R__E__C__G__C__R__*_
                         _-__-__-__-__-__-__-__-__-__-__-__-__-__-__-_
```
