## supplementary table 4 for "High-quality genome assembly for the genetically improved Abbassa Nile tilapia enables the reconstruction of X and Y haplotypes"

**Supplementary Table 3** - Abbassa and GIFT substitutions and indels in *amhy* and *amhΔy* compared to *amh*. Substitutions and indels in noncoding (gene promoter, 5’ UTR, and 3’ UTR) and coding regions of the *amhy* and *amhΔy* compared to *amh* in both the Abbassa and GIFT genome sequence.

| **Strain** | **Gene** | **Annotation** | **Position (bp)** | **Description** | **Nucleotide change** | **Amino acid change** |
| --- | --- | --- | --- | --- | --- | --- |
| Abbassa | *amhy* | Gene promoter | 279-281 | 3 bp insertion | TCT insertion | - |
|  |  | Gene promoter | 300-1600 | Several SNPs and indels (see Additional File 3) | - | - |
|  |  | Gene promoter | >1600 | 117 bp deletion | - | - |
|  |  | Gene promoter | > 1600 | 236 bp deletion | - | - |
|  |  | 5’ UTR | 54 | SNP | G>A | - |
|  |  | 5’ UTR | 72 | SNP | G>A | - |
|  |  | 5’ UTR | 74 | SNP | C>T | - |
|  |  | 5’ UTR | 144 | SNP | G>C | - |
|  |  | 5’ UTR | 252 | SNP | C>T | - |
|  |  | 5’ UTR | 260 | SNP | T>C | - |
|  |  | 5’ UTR | 329 | SNP | T>C | - |
|  |  | Exon VII | 211 | Missense SNP | A>G | Thr (T) > (A) Ala |
|  | *amhΔy* | Gene promoter | 241-406 | 165 bp deletion | - | - |
|  |  | Gene promoter | 300-1650 | Several SNPs and indels (see Additional File 3) | - | - |
|  |  | Gene promoter | >1650 | 479 bp deletion | - | - |
|  |  | 5’ UTR | 22 | SNP | C>T | - |
|  |  | 5’ UTR | 29 | SNP | T>C | - |
|  |  | Exon II | 111 | Missense SNP | T>G | Asp (D) > (E) Glu |
|  |  | Exon III | 66 | SNP | A>T | - |
|  |  | Exon III | 107 | Missense SNP | A>G | Asn (N) > (S) Ser |
|  |  | Exon V | 81 | SNP | C>G | - |
|  |  | Exon VI | - | Premature stop codon | 10 bp insertion | M K R > C R * |
| GIFT | *amhy* | Gene promoter | 279-281 | 3 bp insertion | TCT insertion | - |
|  |  | Gene promoter | 300-1700 | Several SNPs and indels (see Additional File 3) | - | - |
|  |  | Gene promoter | >1700 | 456 bp deletion | - | - |
|  |  | 5’ UTR | 54 | SNP | G>A | - |
|  |  | 5’ UTR | 72 | SNP | G>A | - |
|  |  | 5’ UTR | 74 | SNP | C>T | - |
|  |  | 5’ UTR | 144 | SNP | G>C | - |
|  |  | 5’ UTR | 252 | SNP | C>T | - |
|  |  | 5’ UTR | 260 | SNP | T>C | - |
|  |  | Exon VII | 17 | Missense SNP | C>T | Ala (A) > (V) Val |
|  | *amhΔy* | Gene promoter | 241-406 | 165 bp deletion | - | - |
|  |  | Gene promoter | 410-1600 | Several SNPs and indels (see Additional File 3) | - | - |
|  |  | Gene promoter | > 1600 | 472 bp | - | - |
|  |  | 5’ UTR | 22 | SNP | C>T | - |
|  |  | Exon II | 111 | Missense SNP | T>G | Asp (D) > (E) Glu |
|  |  | Exon III | 66 | SNP | A>T | - |
|  |  | Exon III | 107 | Missense SNP | A>G | Asn (N) > (S) Ser |
|  |  | Exon V | 81 | SNP | C>G | - |
|  |  | Exon VI | - | Premature stop codon | 10 bp insertion | M K R > C R * |

**Supplementary Table 1 – Abbassa and GIFT substitutions and indels in *amhy* and *amhΔy* compared to *amh*.** Substitutions and indels in noncoding (gene promoter, 5’ UTR, and 3’ UTR) and coding regions of the *amhy* and *amhΔy* compared to *amh* in both the Abbassa and GIFT genome sequence.
